# Constructing a Timescale of Biotic Recovery across the Cretaceous–Paleogene Boundary, Corral Bluffs, Denver Basin, Colorado

**DOI:** 10.1101/636951

**Authors:** Anthony J. Fuentes, William C. Clyde, Ken Weissenburger, Antoine Bercovici, Tyler R. Lyson, Ian M. Miller, Jahandar Ramezani, Mark D. Schmitz, Kirk R. Johnson

## Abstract

The Cretaceous–Paleogene (K–Pg) boundary interval represents one of the most significant mass extinctions and ensuing biotic recoveries in Earth history. Earliest Paleocene fossil mammal faunas corresponding to the Puercan North American Land Mammal Age (NALMA) are thought to be highly endemic and potentially diachronous, necessitating precise chronostratigraphic controls at key fossil localities to constrain recovery dynamics in continental biotas following the K–Pg mass extinction. The Laramide synorgenic sedimentary deposits within the Denver Basin preserve one of the most continuous and fossiliferous records of the K–Pg boundary interval in North America. However, poor exposure in much of the Denver Basin makes it difficult to correlate between outcrops. In order to constrain fossil localities across the basin, previous studies have relied upon chronostratigraphic methods such as magnetostratigraphy. Here we present a new high-resolution magnetostratigraphy of 10 lithostratigraphic sections spanning the K–Pg boundary interval at Corral Bluffs located east of Colorado Springs in the southern part of the Denver Basin. Fossil localities from Corral Bluffs have yielded limited dinosaur remains, mammal fossils assigned to the Puercan NALMA, and numerous fossil leaf localities. Palynological analysis identifying the K–Pg boundary in three sections and two independent but nearly identical ^206^Pb/^238^U age estimates for the same volcanic ash, provide key temporal calibration points. Our paleomagnetic analysis has identified clear polarity reversal boundaries from Chron C30n to Chron C28r across the sections. It is now possible to place the fossil localities at Corral Bluffs within the broader basin-wide chronostratigraphic framework and evaluate them in the context of K–Pg boundary extinction and recovery.

## INTRODUCTION

The Cretaceous-Paleogene (K–Pg) boundary is associated with one of the most significant periods of biological turnover in Earth history. It corresponds to one of the three largest mass extinctions and is followed by a profound adaptive radiation of modern taxa, most notably mammals, in the Paleocene (Schulte et al., 2010; Halliday et al., 2017). While the drivers and signature of the K–Pg mass extinction is the subject of active and extensive discussion in the literature (Alvarez et al., 1980; Courtillot et al., 1986; Schulte et al., 2010; Schoene et al., 2015, 2019; Sprain et al., 2019), the recovery dynamics in the subsequent earliest Paleocene have received much less attention. A more comprehensive understanding of the recovery after the rapid and catastrophic end Cretaceous mass extinction may provide insights into how the modern biosphere will respond to the ongoing anthropogenic driven extinction event.

Numerous vertebrate fossil localities spanning the Late Cretaceous to Paleocene have been identified within western North American interior basins (Eberle and Lillegraven, 1998; Clemens, 2002; Hunter and Archibald, 2002; Eberle, 2003; Lofgren et al., 2004; Middleton et al., 2004; Wilson, 2013). The mammalian faunas comprising the Puercan North American Land Mammal Age (NALMA) are especially significant as this interval coincides with the onset of biotic recovery immediately following the K–Pg mass extinction (Lofgren et al., 2004). While the Western Interior of North America includes the best record of early Paleocene mammalian fossil localities in the world, the total number of localities and overall availability of fossil material remains limited. The identification of new localities and continued sampling of existing localities is critical to isolating the factors driving the recovery following the K–Pg mass extinction. Moreover, temporal constraints provided by high-resolution chronostratigraphic studies are critical to pace the recovery within continental strata. Puercan mammalian faunas are highly endemic making it difficult to identify biostratigraphic patterns across the region, further necessitating the use of chronostratigraphic methods that are independent of biostratigraphy in order to properly deconvolve the regional scale drivers of recovery (Lofgren et al., 2004). Magnetostratigraphy is commonly and effectively used for this purpose. This method correlates an observed pattern of paleomagnetic polarity reversals in a stratigraphic sequence to the independently dated Geomagnetic Polarity Time Scale (GPTS; Gradstein et al., 2012; Ogg, 2012), thereby constraining a stratigraphic section to a well-calibrated time interval.

The Laramide synorgenic sediment sequence within the Denver Basin represents one of the best terrestrial records of the K–Pg boundary in the world with observed osmium and iridium anomalies, shocked quartz, extinction of flora and fauna, and a spike in abundance of fern spores that are commonly used as indicators of the boundary (Nichols and Fleming, 2002; Barclay, 2003; Raynolds and Johnson, 2003; Zaiss et al., 2014). Several important megafloral and vertebrate paleontological localities have been identified within the basin; however, the Denver Basin is relatively under-sampled in comparison to coeval sites in the San Juan Basin of New Mexico and Williston Basin of North Dakota and Montana (Archibald, 1982; Williamson, 1996; Clemens, 2002; Williamson et al., 2012; Wilson, 2013; Smith et al., 2018). There are 231 Puercan-aged mammal localities globally recorded in the Paleobiology Database, 185 of which are in the Rocky Mountain region of North America, and of these, 101 sites are from New Mexico, North Dakota, and Montana (Paleobiology Database accessed on 5/1/2019).

Previous discoveries of an *in-situ* rainforest at Castle Rock and the anomalously high mammalian diversity of the Littleton Local Fauna, both in the Denver Basin, allude to the incredible potential this region has in yielding informative fossil material to help elucidate the post K–Pg extinction recovery dynamics within the central Western Interior of North America (Johnson and Ellis, 2002; Ellis et al., 2003; Middleton et al., 2004; Dahlberg et al., 2016). A barrier to developing a coherent stratigraphic framework for existing and new fossil localities in the Denver Basin is the disparate nature of exposure making it difficult to tie localities to one another via lithostratigraphy. Fortunately, magnetostratigraphy calibrated through palynology and radiometric ages can surmount this barrier and provide an independent chronology of biologic turnover in the Denver Basin that can be compared to other basins.

Much of the previous paleontological and chronostratigraphic work in the Denver Basin has been focused in the central portion of the basin in and around Denver, highlighting the need for more detailed, expansive prospecting, and stratigraphic correlation in other parts of the basin (Hicks et al., 2003; Clyde et al., 2016). Previous studies have identified Late Cretaceous through Paleocene fossil leaf localities and Puercan mammal localities at the Jimmy Camp Creek and Corral Bluffs field site (hereafter referred to as “Corral Bluffs”), located east of Colorado Springs in the southernmost part of the basin (Johnson et al., 2003; Eberle, 2003; Figure 1). This study seeks to construct a high-resolution chronostratigraphy for the Corral Bluffs area so that existing and future Late Cretaceous-Early Paleocene fossil localities can be placed in a precise temporal context. This will be achieved through compiling a magnetostratigraphic framework calibrated by U–Pb geochronology and a palynologically defined K–Pg boundary. This new chronology for Corral Bluffs will allow the lithostratigraphic units and associated fossil localities within this field area to be tied into the broader basin-wide chronostratigraphic framework, which will ultimately help better constrain the timing and drivers of the post-K–Pg extinction recovery in this region.

**Figure 1:**
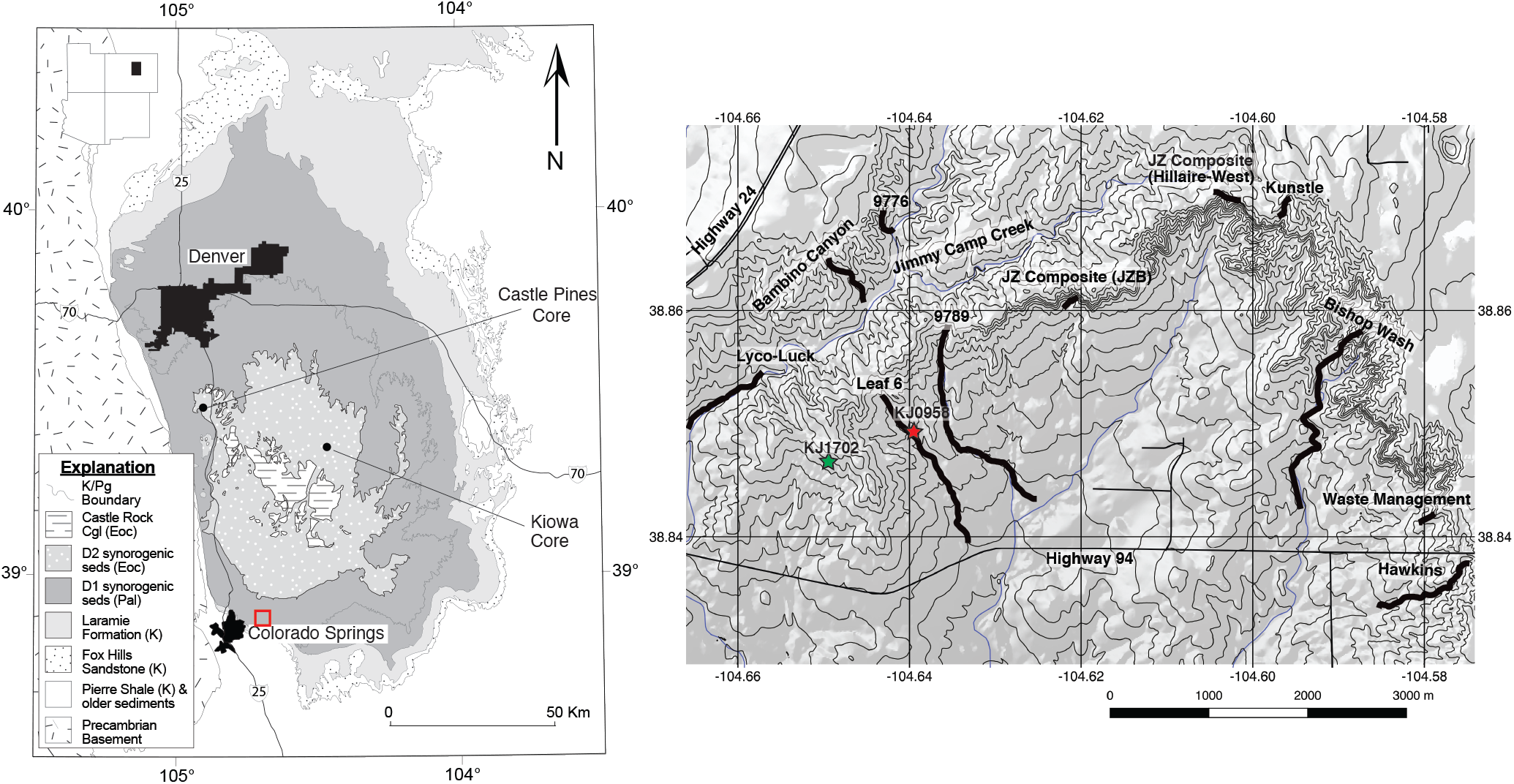
Geologic map (left) of the Denver Basin modified from Clyde et al. 2016 showing the locations of the Castle Pines and Kiowa Cores in relation to Corral Bluffs/Jimmy Camp Creek area (red square). Map (right) of the Corral Bluffs/Jimmy Camp Creek field area showing the ten stratigraphic sections studied here (black lines with black text labels) and location of ash samples KJ0958 (red star) and KJ1702 (green star).

### Extinction and Recovery across the K–Pg Boundary

Mass extinctions are typified by an initial catastrophic loss of taxa that is followed by a low-diversity “recovery” interval dominated by surviving taxa (Erwin, 2001; Hull et al., 2015). The detailed patterns of biotic recovery following the K–Pg mass extinction, however, are complicated and subject to regional biogeographic and taxonomic variability. For example, there is ongoing debate as to whether distance from the Chicxulub impact site can serve as proxy for the severity and timing of biotic recovery across the K–Pg boundary (Witts et al., 2016; Lowery et al., 2018). Consistent with this idea, are numerous studies of paleofloral assemblages suggesting that North America, which is relatively close to the Chixculub impact site, experienced higher rates of extinction, and delayed recovery, compared to southern hemisphere sites (Erwin, 2001; Nichols and Fleming, 2002, Iglesias et al., 2007; Clyde et al., 2014). However, a recent study of the diversity of benthic marine organisms within the Chicxulub Crater itself concluded that biologic productivity and diversity had recovered within 30 kyr, in contrast to sites in the other parts of the Gulf of Mexico that took 300 kyr or more to return to pre-impact levels (Lowery et al., 2018). Such heterogeneous patterns of K–Pg extinction and recovery have been documented for numerous other taxonomic groups and biogeographic regions suggesting this is more the rule than the exception (Jablonski, 1998; Håkansson and Thomsen, 1999; Kiessling and Baron-Szabo, 2004; Vajda and McLoughlin, 2007; Hull and Norris, 2011; Mizukami et al., 2013; Vajda and Bercovici, 2014; Witts et al., 2016; Donovan et al., 2018).

Floras of the Late Cretaceous are largely dominated by angiosperms, and the K–Pg boundary in the Western Interior of North America is associated with a ~57–78% loss of plant diversity (Johnson, 2002) as well as a ~33% extinction in the pollen record (Nichols and Johnson, 2002). The immediate recovery is characterized by the fern spike interval representing the first few thousand years after the bolide impact during which pioneer plants recolonize the devastated landscape, followed by a low-diversity interval dominated by dicotyledonous angiosperms (Johnson, 2002; Barclay, 2003; Vajda and Bercovici, 2014; Clyde et al. 2016; Fastovsky and Bercovici, 2016). Regional heterogeneity is also observed in the vegetation of the Western Interior as localities in Saskatchewan, Canada record a mass-kill of standing vegetation but no clear evidence of mass extinction of paleoflora (McIver, 1999; Johnson and Ellis, 2002; Vajda and McLoughlin, 2007), whereas the palynological record shows a similar signature to that of the Western Interior (Sweet and Braman, 1992; 2001; Sweet et al., 1990; 1999).

Despite ongoing debate as to the origination of crown placental mammals, the most important diversification of eutherian mammals seems to coincide with the K–Pg boundary (O’Leary et al., 2013; Grossnickle and Newham, 2016; Davies et al., 2017). The Western Interior presents the best-documented record of the K–Pg extinction on land with the highest density of vertebrate fossil localities spanning the boundary (Clemens, 2002). As noted for other groups, the recovery dynamics of mammals from the Western Interior of North America are heterogenous as the speed and severity of turnover at the K–Pg boundary vary depending on the spatiotemporal resolution of the study (Wilson, 2013; Longrich et al., 2016; Close et al., 2019). In general, however, rapid and severe declines in mammalian taxonomic diversity are reported across North America during this interval (Wilson, 2014; Longrich et al., 2016). Mammalian taxa with more widely spread geographic ranges and simpler morphologies corresponding to an omnivorous feeding mode are associated with a higher likelihood of crossing the K–Pg boundary (Wilson, 2013; Longrich et al., 2016).

The Pu1 (*Protongulatum/Ectoconus*) land mammal zone is characterized primarily by low ecological and morphological diversity in the western interior, mirroring the diversity trends of the paleoflora (Wilson, 2014). Opportunistic immigrant taxa spearheaded by archaic ungulates are common in the earliest Paleocene, possibly from refugia either to the north or south of the Western Interior (Lofgren et al., 2004; Wilson, 2014). Assemblages from the Hell Creek region of Montana indicate that recovery of mammal communities in this area took between 320 kyr to 1 myr with eutherian taxa taking 400 kyr to 1 myr to recover (Wilson, 2014; Smith et al., 2018). Lillegraven and Eberle (1999) noted that the eutherian faunas in the Ferris Formation of the Hanna Basin, the only locality currently where Pu1-3 are in direct superposition, do not show significant losses at the K–Pg but instead radiate to fill vacant niche space. The onset of the Pu2 (*Ectoconus/Taeniolabis taoensis*) interval zone has been noted to coincide with an increase in both body size and morphologic complexity of mammals, most notably the rapid diversification of archaic ungulates (Lillegraven and Eberle, 1999; Lofgren et al., 2004). The Pu2 and Pu3 (*Taeniolabis taoensis/Periptychus carinidens)* interval zones also coincide with increasing north to south provinciality of mammalian assemblages (Lofgren et al., 2004). The provinciality of Pu2/3 faunas is often confounded by stratigraphic concerns (e.g. reworking, incision by channels) at important localities (Lillegraven and Eberle, 1999). The Pu3 interval continues the diversity trends established during the Pu2 interval with considerable compositional overlap between the two intervals (Eberle and Lillegraven, 1998). While details of the onset and tempo of initial recovery for North American mammals are still largely unresolved, by the Pu2/Pu3 transition, mammal communities had largely recovered across the Western Interior and began to expand rapidly into new available niche space (Wilson, 2014).

The global patterns and rates of extinction and recovery during and following the K–Pg mass extinction appear to be more stochastic than simply a function of proximity to the impact site. Recovery dynamics may instead be primarily governed by biotic factors like evolution of novel adaptations and immigration, and high-resolution temporal frameworks are needed to better understand the tempo and mode of these processes.

## METHODS

### Paleomagnetism

#### Field Sampling

We collected 548 oriented hand samples from 126 sites in 10 stratigraphic sections across the Corral Bluffs field area over the course of the 2017 and 2018 field seasons (Figure 1). Thirty-two samples from ten previously unanalyzed sites from the Leaf 6 section collected during a 2009 field reconnaissance were also included in this analysis, for a total of 136 sites analyzed. Sampling was guided by the previous magnetostratigraphic work of Hicks et al. (2003) who identified polarity reversals corresponding to the C30n–C29r and C29r–C29n boundaries in Jimmy Camp Creek (our Lyco-Luck section) and the JZ section. Our sampling was limited to fine-grained lithologies such as mudstones, siltstone, or muddy, fine-grained sandstones. Overburden was removed before sampling to minimize the effects of modern weathering. Slumps and channels were avoided for sampling although site CB1804 from the 9776 section was later determined to be sampled from slumped material and was not included in the final data table (Table S1).

A GNSS differential global positioning system (DGPS; Trimble June T41S Model) was used to collect waypoints at each sample site and these data were post-processed using a nearby Continuously Operating Reference Station. These DGPS data were used in conjunction with hand levelling to determine stratigraphic levels within a section. Correlation between sections is challenging. Local lithologic controls are available from the 9789 section to the Bishop Wash section in the form of traceable sandstone bodies. The westernmost sections (Lyco-Luck, Bambino Canyon, Leaf 6, and 9776), however, are difficult to correlate lithostratigraphically due to low relief, variable dips, and large covered areas. In order to account for the heterogeneity in available lithostratigraphic tracers across the sections, elevation from the DGPS measurements is used as the primary stratigraphic framework for this study (Table S1 and S2). Elevation error for DGPS measurements like these is typically less than 0.5 meters so the superposition of almost all sample sites was resolved correctly. In cases where the superposition of closely spaced sites was not properly resolved by DGPS (e.g. because of steep overhanging cliffs causing poor GPS reception) or where we had clearly erroneous or missing DGPS measurements, we estimated elevation values (signified by italics in Tables S1 and S2). These estimated values were measured using hand levelling or interpolated using field notes and Google Earth to calculate the proportional elevation difference between two bracketing sites with definitive DGPS values and the unknown site. The difference between the two bracketing DGPS elevations was multiplied by the proportion calculated from the Google Earth analysis and added to the elevation of the lower sample to derive the estimated elevation for the unknown site. Site DB0902 was sampled from the same outcrop as CB1735 but during a different field season and was estimated to lie no more than 1 meter below CB1735 (Table S1). Stratigraphic positions for CB1805 and CB1809 could not be estimated because they are located in a subsidiary gully to the main Bambino section and therefore are not included in the magnetostratigraphic results. The estimate stratigraphic thickness of the Lyco-Luck section within Jimmy Camp Creek appears to be more compressed relative to the other sections probably due to its relatively long horizontal distance, small change in elevation, and relatively steep 3° dip.

We acknowledge that the stratigraphic thicknesses determined using DGPS elevation are minimum estimates of true thicknesses, however this method allows us to use a completely independent and objective frame of reference to identify shared polarity reversals across sections that could not be correlated lithostratigraphically due to outcrop cover. Also, dips in the field area are generally < 3° and difficult to measure accurately, so elevation serves as a reasonable proxy for stratigraphic level. Our identification and correlation of the paleomagnetic reversals at Corral Bluffs within an absolute elevation framework will allow for future work to more comprehensively model stratigraphic thicknesses using the Chron boundaries as stratigraphic datums.

#### Laboratory Methods

Paleomagnetic hand samples were cut into 8 cm^3^ cubes, retaining the oriented surface as one cube face, and analyzed at the University of New Hampshire (UNH) Paleomagnetics Laboratory. Samples were measured using a 2G SQUID cryogenic magnetometer shielded from the background magnetic field. Demagnetization protocol was determined using data from pilot samples and previous studies (Hicks et al., 2003; Clyde et al., 2016). Samples were first subjected to stepwise alternating field (AF) demagnetization via a Molspin tumbling alternating field demagnetizer at 3 mT steps up to 30 mT, 5 mT steps between 30 mT and 60 mT, and finally 10 mT steps between 60 mT and 100 mT until the natural remnant magnetization (NRM) fell below the detection level of the magnetometer or the NRM no longer decreased in intensity. Samples that did not demagnetize according to the AF protocol were then subjected to additional thermal demagnetization in an ASC Model TD48 SC thermal demagnetizer at 25, 80, 115, 135, 160, 250, 300, 350, 400, 450, 500, 525, 560, 580,640, 690 °C temperature steps.

#### Data Analysis

After completion of the demagnetization protocol, the sample data were analyzed using the PuffinPlot paleomagnetic data analysis program (Lurcock and Wilson, 2012). Samples with three or more sequential steps exhibiting linear or quasi-linear decay to the origin were characterized using principal component analysis (Kirschvink, 1980). For these samples, only those with a maximum angular deviation (MAD) of ≤ 20° were included (Figure 2). A Fisher mean was calculated for samples that displayed an initial decay followed by strong clustering of vector end points and no further decay (Fisher, 1953; Figure 2). Some samples with overlapping unblocking spectra display a demagnetization path that was best characterized by a great circle (Figure 2).

**Figure 2:**
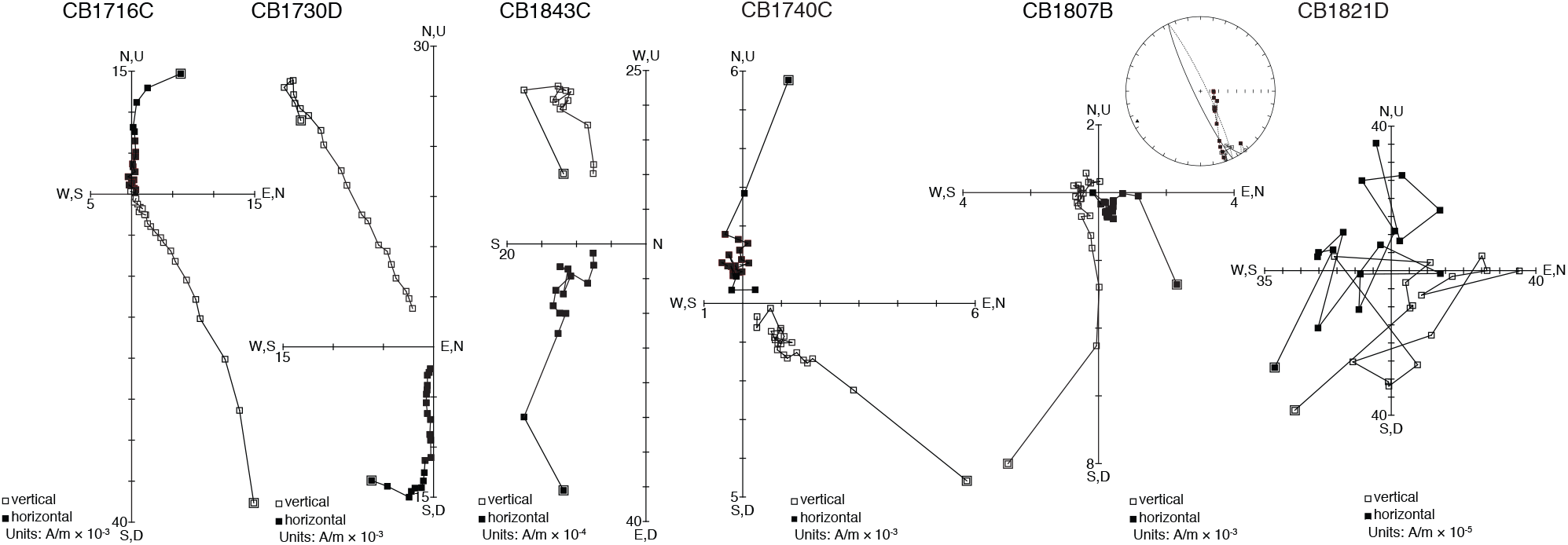
Representative vector end point diagrams for selected samples. CB1716C is a normal polarity sample characterized by a PCA with a MAD = 3.5°. CB1730D is a reversed sample characterized by a PCA with a MAD = 2.1°. CB1843C is a reverse sample characterized by a fisher mean due to strong clustering of vector endpoints (Fisher, 1953). CB1740C is a normal polarity sample exhibiting strong clustering of vector endpoints and is characterized by a fisher mean. CB1807B is a reverse polarity sample exhibiting a direction best characterized by the great circle analysis of McFadden and McElhinny (1998). Equal area projection for CB1807B with a great circle defining sample ChRM is inset adjacent to that sample’s vector end point diagram. CB1821D is a sample exhibiting chaotic demagnetization behavior and was not able to be characterized using a PCA, fisher mean, or great circle analysis.

Alpha sites for this study are described using Fisher statistics and defined as sites that pass the Watson test for randomness (Watson, 1956; Figure 3) and included three or more samples whose characteristic remanent magnetization (ChRM) was defined by either a principle component analysis (PCA) or Fisher mean (or some combination). Beta sites are sites that have three or more sample directions but include at least one sample characterized by great circle analysis (McFadden and McElhinny, 1988). Well characterized sample directions from sites that do not meet the alpha or beta criteria are still reported (Table S2) and considered in the magnetostratigraphic analysis.

**Figure 3:**
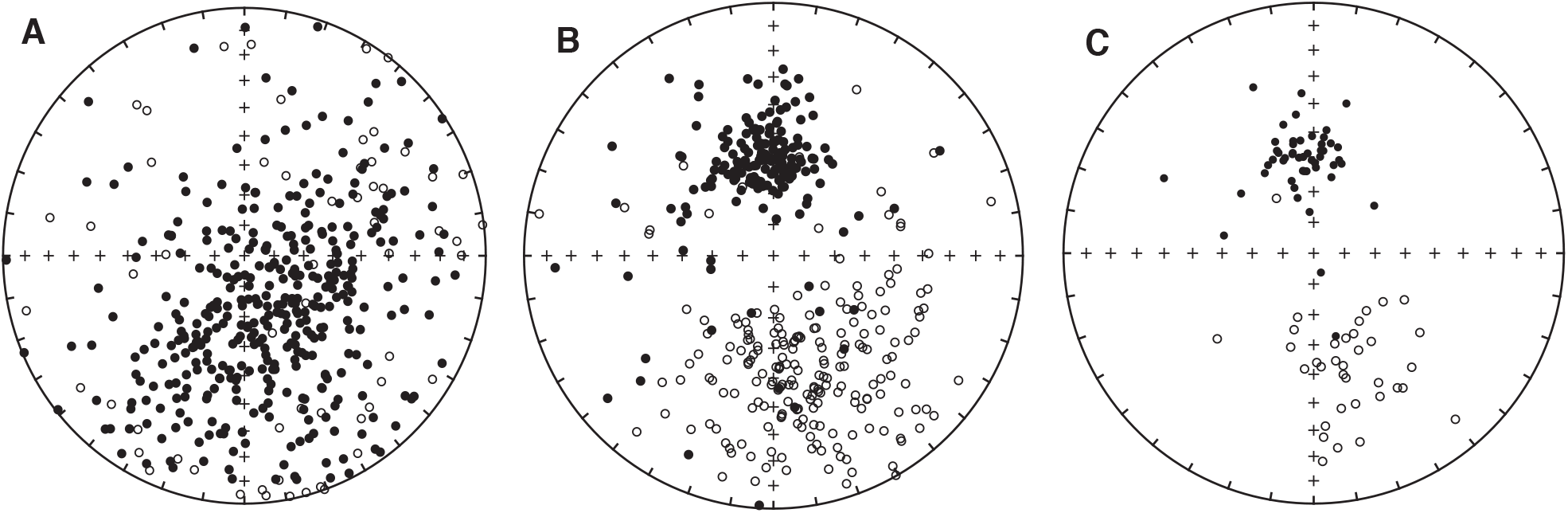
Equal area projections for paleomagnetic sample and alpha site directions. **A.** Equal area projections of all NRM paleomagnetic sample directions, **B.** ChRM directions for paleomagnetic samples with coherent demagnetization behavior, **C.** Alpha site mean ChRM directions.

#### Isothermal Remanent Magnetization

Seven samples from sites spanning the field area were selected to undergo isothermal remanent magenetization (IRM) experiments in order to better understand the mineralogy governing the magnetic remanence. Samples were selected from sites within sections across the field area laterally as well as from the lower sections (Lyco-Luck) to the highest section stratigraphically (Kunstle), in order to provide a comprehensive spread in lithologies and ages. Additionally, this helped to test whether poorly behaved samples in the stratigraphically lower Lyco-Luck section differed significantly in magnetic mineralogy relative to well-behaved sites that are stratigraphically higher. The IRM samples were first cut into cubes < 1 cm^3^ and then the x-axis of each cube was subjected to a magnetic field of increasing strength within an ASC IM10 impulse magnet at discreet steps from 0 T up to 1.1 T and measured in the 2G magnetometer after each step (Figure 4). Upon reaching the 1.1 T step, the samples were then subjected to a backfield applied in the opposite direction in steps of 100 and 300 mT in order to isolate the proportion of remanence held by the soft and hard components. From the backfield magnetization, it is possible to determine the contributions to bulk remanence of hard antiferromagnetic components (e.g. hematite, goethite, etc.) and soft ferrimagnetic components (e.g. magnetite, maghemite, etc.) through the calculation of the S-Ratio (Stober and Thompson, 1979; Bloemendal et al., 1992; Maxbauer et al., 2016). The S-Ratio is defined as half of the difference between the saturation isothermal remanent magnetization (SIRM) and the IRM at 300 mT (IRM__300mT_), divided by the SIRM (Maxbauer et al., 2016). Following the three axis IRM method of Lowrie (1990), a field of 1.1 T was then applied to the x-axis, 0.4 T to y-axis, and 0.12 T to the z-axis of the samples to separate the magnetization of mineral fractions of different coercivities along each axis. The samples were then subjected to thermal demagnetization at the following temperature steps: 25, 50, 86, 109, 132, 150, 209, 250, 275, 300, 325, 350, 375, 400, 500, 540, 560, 580, 600, 621, 650, and 680 °C. The magnetic intensities of the three orthogonal axes were measured at each step and compiled into three-axis demagnetization curves.

**Figure 4:**
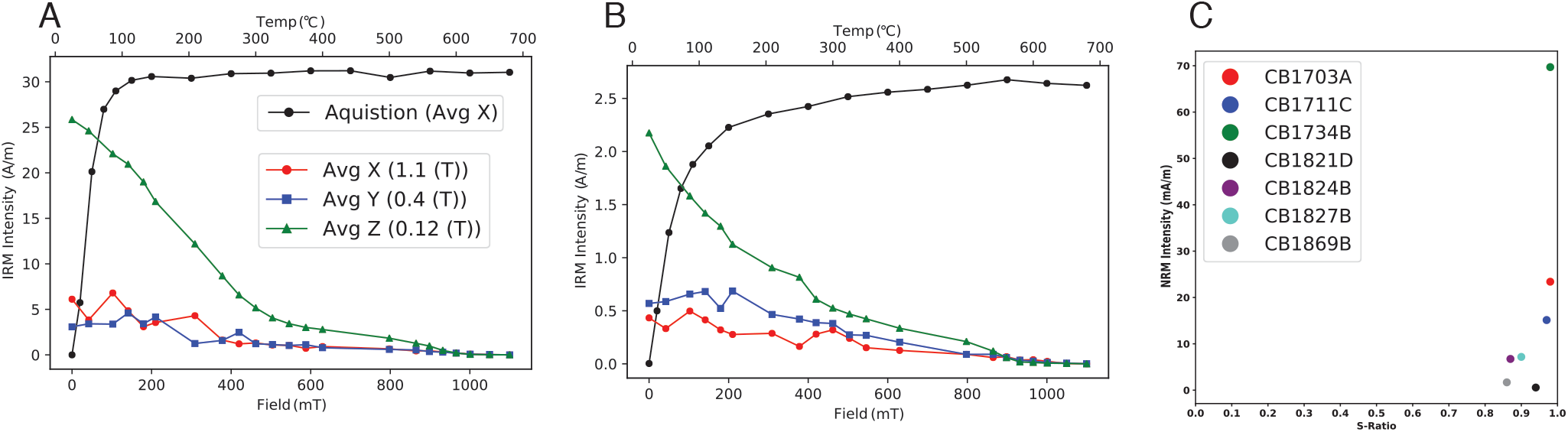
IRM acquisition and three-axis demagnetization curves for two representative samples. **A.** CB1703A showing sample dominated by magnetite. **B.** CB1869B showing sample dominated by magnetite but influenced by minor amounts hematite. **C.** NRM intensity (mA/m) vs S-Ratio for all IRM samples showing that remanence in all samples was carried dominantly by magnetite. See Table S4 and Figure S5 for complete IRM data.

### U–Pb Geochronology

Two samples of volcanic ash from separate sites in the Corral Bluffs field area were collected and analyzed for U–Pb geochronology. Sample KJ0958 (locally known as the “Feral Llama Ash”) was collected within the Leaf 6 stratigraphic section at 38.846403°N, 104.649671°W, 1900.4 m elevation (NAD83 datum). Sample KJ1702 (locally referred to as the “Stock Tank Ash”) was collected ~940 meters to the W/SW of KJ0958 at 38.849187°N, 104.639530°W, 1930.9 m elevation (NAD83 datum). Both ash deposits are expressed as thin (< 1-2 cm), gray, lensy tonsteins within 10–20 cm thick lignite deposits, similar to ashes previously dated from the central part of the basin (Clyde et al., 2016). The limited occurrence of outcrop between the two ash locations does not expose the proper stratigraphic level to allow tracing the ashes directly, but sufficiently exposes adjacent horizons to show the gradual change of apparent dip from the Leaf 6 location (about 2° east) to the Stock Tank location (about 4-5° east). This simple change in apparent dip is consistent with a single ash layer having produced the measured elevation difference between the two age-dated locations. Based on the similar mode of occurrence of the two tephras, the similar succession of coal-bearing and arkosic strata at the two ash locations, and the structural dip model between the two locations, we interpret these ashes to represent the same volcanic deposit (Figure S1). This interpretation is reinforced by the observation that samples KJ0958 and KJ1702 have yielded U–Pb ages that are equivalent within uncertainty (see Results below).

Sample KJ0958 was processed and analyzed using the chemical abrasion isotope dilution thermal ionization mass spectrometry (CA-IDTIMS) method at the Massachusetts Institute of Technology Isotope Laboratory. Sample KJ1702 was processed and analyzed using LA-ICPMS and CA-IDTIMS at the Boise State University Isotope Geology Laboratory. Detailed analytical methods for both samples can be found in the Supplement. The calculated U-Pb date uncertainties are reported in ± *X*/*Y*/*Z* Ma format to account for internal versus fully propagated uncertainties (including tracer calibration and U decay constant errors; see Supplement).

### Palynological Analysis

Palynological analysis was conducted on a total of 46 samples from three sections to identify the placement of the K–Pg boundary in the study area: Bishop Wash (ADB1701 and ADB1702), Leaf 6 (ADB1703), and 9789 (ADB1802). Samples were processed using standard palynological procedure by Global GeoLab Ltd., Canada. Approximately 20 grams of bulk rock was treated with dilute hydrochloric acid (HCl), macerated in 75% hydrofluoric acid (HF), and cleaned of fluorosilicate byproducts in hot HCl. Organic residue was sieved, the fraction between 10 µm and 200µm was kept, dyed with safranin red, and mounted in polyester resin on one or two microscope slides, depending on recovery. Slides are reposited at the Denver Museum of Nature & Science (DMNS). Organic matter recovery ranges from excellent to fair, with some barren samples and some samples showing a significant amount of older reworked palynomorphs. The position of the K–Pg is indicated by the major decrease of the relative abundance of Cretaceous indicators (K-taxa) without subsequent recovery (Bercovici et al., 2009; Table S8). Paleocene samples are characterized by the appearance of several pollen species belonging to the *Momipites* genus.

## RESULTS

### Paleomagnetism

#### Demagnetization Behavior

The majority of samples responded favorably to the AF demagnetization protocol, with only one site (CB1703B) needing additional thermal steps to be fully characterized. Thermal demagnetization applied after AF demagnetization did not prove to be effective for the vast majority of samples as the remaining high coercivity component typically did not exhibit clear demagnetization behavior when subjected to these additional thermal steps. This was likely due to the presence of goethite which was observed in some IRM samples (see below). Of the 426 samples that were analyzed, 185 displayed a linear decay to the origin with characteristic remanent magnetizations (ChRMs) defined by PCAs with MADs less than 20°, 121 samples exhibited initial decay followed by strong clustering of vector endpoints that were calculated using Fisher means, and 69 samples were characterized by great circle analysis due to the overlapping unblocking spectra between the components of the natural remanent magnetization of the samples (Figure 2,3). The 51 remaining samples were not able to be characterized due to highly chaotic demagnetization behavior.

#### Site Calculations

Of the 134 sites for which three or more samples were analyzed, 84 sites fit the criteria for alpha sites and 23 sites are considered beta sites (Table S1). Some sites did not meet the alpha or beta site criteria but still included samples with stable demagnetization so these sample ChRM directions are plotted in their respective sections’ magnetostratigraphic plots to help further guide our interpretation. Polarity determinations were not possible for sites CB1818, 1819, 1822, 1849, 1852, 1862, 1874, and 1876 due to highly chaotic behaviors of the samples from these sites and were not considered further for analysis.

#### Reversal Test

The reversal test is a standard field stability test in paleomagnetism that tests whether the two polarities in a paleomagnetic data set are antipodal to one another as one would expect under the Geocentric Axial Dipole (GAD) model of the Earth’s magnetic field (Cox and Doell, 1960; Heslop and Roberts, 2018). The mean declination and inclination of the normal polarity alpha sites is 351.9/58.4 and the overall mean for all normal polarity alpha and beta sites is 349.1/57.8. The mean declination and inclination for the reverse polarity alpha sites is 162.7/-46.9 and the mean for all reverse polarity sites is 164.4/-48.3. The 83 alpha sites of both normal and reversed polarities narrowly passed the parametric bootstrap reversal test of Tauxe et al. (2018) at the 95% confidence level (Figure S2). The narrow passage of the reversal test is likely due to a small but persistent present-day normal polarity overprint causing reversed polarity samples to have slightly shallower inclinations than normal polarity samples.

#### Polarity Determinations and Reversal Placement

Polarity determinations for most sites is very clear so paleomagnetic reversals are positioned stratigraphically halfway between two superpositionally adjacent alpha or beta sites of opposite polarity (Figure S3). The 9776 section exhibits a polarity oscillation close to a reverse-to-normal polarity reversal near the base of the section (Figure S3). Sampling of this interval was undertaken along a short ridge and no slumping was apparent and it is likely that the oscillation in polarity corresponds to an instability in the field near the paleomagnetic reversal. In this case, the paleomagnetic reversal was placed above the highest reverse site and below the subsequent normal site.

Nine of the 10 sections recorded at least one polarity reversal. The lowest sections (Lyco-Luck, Leaf 6, 9789; Figure S3) exhibit a normal polarity interval that switches to reverse polarity at an average elevation of 1887 m. Most of the stratigraphically higher sections (Bambino Canyon, 9776, 9789, JZ, Bishop Wash, Waste Management; Figure S3) record a reversed polarity interval at the base of the sections that switches to a normal polarity interval at an average elevation of 1952 m. The Hawkins section is entirely characterized by reverse polarity (Figure S3). The highest section (Kunstle; Figure S3) has a normal polarity zone at the base that transitions to reverse polarity at ~2039 m elevation. The 9789 section includes two paleomagnetic reversals, from normal to reverse polarity at ~1880 m elevation and from reverse to normal at 1960.8 m elevation (Figure S3).

#### Paleomagnetic Pole

The paleomagnetic pole calculated from the alpha sites is 135.3°/81.3° (α95 = 4.7°) and does not overlap with the unflattened or flattened paleomagnetic poles of North America from 70 to 60 Ma (Besse and Courtillot, 2003; Torsvik et al., 2012; Figure S4). Several factors may be causing this discrepancy. Inclination shallowing and a weak but persistent present-day overprint causing shallower than expected directions for reversed sites are likely the predominate contributors to the somewhat “far-sided” pole for this study.

#### Isothermal Remanent Magnetization Experiments

The IRM acquisition curves for all samples from Corral Bluffs are characterized by a steep increase in intensity up to ~300 mT which is consistent with low coercivity species such as magnetite, titanomagnetite, and maghemite (Figure 4, S5). The S-ratios calculated from the backfield steps (Maxbauer et al., 2016) are all greater than 0.5 which is also indicative of a majority of the bulk remanence held by low coercivity components (Figure 4). CB1824B, CB1827B, and CB1869B show small increases in intensity past 300 mT which is indicative of the presence of higher coercivity minerals such as hematite and goethite in these samples. The backfield data for these samples shows that, although these samples are influenced by a high coercivity component, they all have S-ratios greater than 0.8 so are still dominated by a low coercivity phase.

The three-axis IRM demagnetization curves indicate that the soft axes (0.12 T) in all samples initially have by far the highest intensity which is consistent with remanence held by low coercivity minerals (Figure S5). There is a considerable drop in intensity along the 0.12 T axes of all the samples from 200–500 °C followed by generally another significant drop from 540–580 °C at which point the intensity of the soft axes of the samples are essentially null. It is difficult to distinguish between the acquisition and demagnetization properties of magnetite and either titanomagnetite or titanomaghemite. Both titanomagnetite and titanomaghemite have coercivities that are approximate to that of magnetite and can have unblocking temperatures ranging from room temperature to > 600 °C (Özdemir and Banerjee, 1984; Lowrie, 1990). Given the fluvial depositional environment of the sediment at the field site, it is likely that the primary low coercivity component is detrital magnetite/titanomagnetite eroded and deposited during the uplift of the Front Range. Maghemite is also likely present in these rocks as it often forms as a product of oxidation of magnetite, either through weathering or pedogenic processes (van Velzen and Dekkers, 1999). CB1821D, CB1824B, and CB1869B show a slight dip in intensity along the hard axis (1.1 T) between 50–150 °C which is evidence of a small amount of goethite within these samples (Figure S5 D, E, F). CB1824B, CB1827B, and CB1869B have the smallest S-ratios and also show a final decrease in intensity along the hard axis above the 580 °C step (Figure 4 A, B; S5 E, F, G). This is characteristic of hematite, which is the primary hard coercivity component within these samples. The minor amounts of goethite and hematite in these samples indicates that these facies likely underwent weak pedogenesis soon after deposition..

### U–Pb Geochronology

#### Sample KJ0958

All seven zircons selected and analyzed by the CA-IDTIMS method from the Corral Bluffs ash bed define a statistically coherent cluster of ^206^Pb/^238^U dates with a weighted mean date of 66.253 ± 0.031/0.045/0.084 Ma and a mean squared weighted deviation (MSWD) of 0.36. This date best represents the age of zircon crystallization and a good approximation for the age of Feral Llama ash deposition (Figure 5; Table S5).

**Figure 5:**
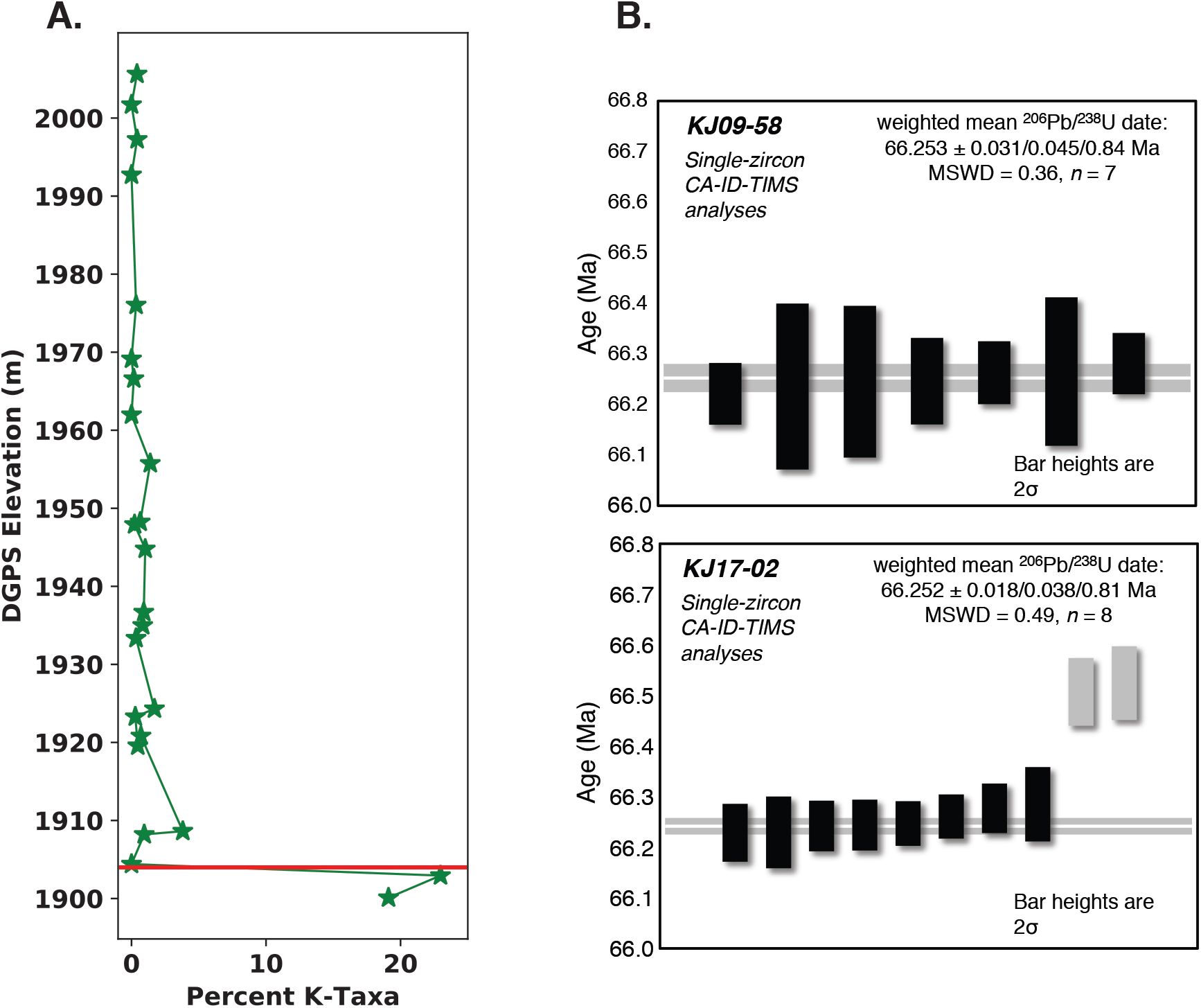
Palynological and ^206^Pb /^238^U age constraints. **A.** Percent of Cretaceous palynomorph counts per sample plotted by DGPS elevation for the Bishop Wash section. Red line represents placement of K-Pg boundary in this section. **B.** Distribution of ^206^Pb/^238^U ages of analyzed zircons from the Feral Llama (KJ09-58) and Stock Tank (KJ17-02) ash beds of the Corral Bluffs sections of the Denver Basin. Vertical bars represent individual zircon analyses with the bar height proportional to 2σ analytical uncertainty of the ^206^Pb/^238^U dates. Solid bars are analyses used in date calculation. Horizontal line and its shaded envelope signify the weighted mean date and it 95% confidence level uncertainty, respectively. See Tables S6 and S7 for complete U–Pb isotopic data and the Supplement for explanation of stated uncertainties.

#### Sample KJ1702

Most of the zircon population of sample KJ1702 consists of large (majority greater than ~100 µm), generally euhedral grains, many of which possess dark inherited cores. Thirty-five of the 81 laser ablation inductively coupled plasma mass spectrometry (LA-ICPMS) analyses yielded ^206^Pb/^238^U dates between 61–77 Ma, and the majority of the remaining analysis yielded dates greater than 400 Ma (Table S6).

Thirteen zircon grains were selected from the youngest LA-ICPMS population for CA-IDTIMS analysis. Two grains (z7 and z8) yielded reversely discordant data, assumed to be laboratory error, and were discarded. Two grains (z5 and z11) yielded older, concordant dates of ~66.5 Ma and were discarded. One grain (z4) was dissolved during chemical abrasion and was lost. The remaining eight CA-ID-TIMS analyses resulted in a weighted mean ^206^Pb/^238^U date of 66.252 ± 0.018/0.038/0.081Ma (MSWD = 0.49), which is interpreted to represent the age of crystallization preceding eruption of the sample (Figure 5; Table S7).

### Palynology

Palynological assemblages of the lower portions the Leaf 6, 9789, and Bishop Wash sections correspond to the upper Maastrichtian *Wodehouseia spinata* Assemblage Zone (Nichols, 1994; Bercovici et al., 2012). The sharp decline in relative abundance without subsequent recovery (Bercovici et al., 2009) of typical Cretaceous taxa, or “K-taxa” (Nichols et al., 2002), is indicative of the terrestrial floral demise associated with the K–Pg boundary and the Chicxulub impact (Nichols and Johnson, 2002, 2008; Vajda and Bercovici, 2014; Fastovsky and Bercovici, 2016). The identified K-taxa from Corral Bluffs are *Aquilapollenites trialatus*, *A. calvus*, *A. turbidus*, *A. collaris*, *A. conatus*, *A. marmarthensis*, *A. mtshedlishvili*, *A. quadrilobus*, *A. reductus*, *Ephedripites multipartitus*, *Erdtmanipollis* spp., *Libopollis jarzeni*, *Liliacidites altimurus*, *L. complexus*, *Retibrevitricolporites beccus*, *Striatellipollis striatellus*, *Tricolpites microreticulatus*, and *Tschudypollis* spp. In addition to the above pollen taxa, the epiphyllous fungal thallus *Trichopeltinites* sp. is also present, and sometimes very common with relative abundance of up to 59.8%. *Trichopeltinites* has been reported to disappear at the K–Pg boundary, presumably along with its macrofloral host (Nichols and Johnson, 2008), and thus has been treated here as a K-taxa. The relative abundance drop in K-taxa and the palynologically defined K–Pg boundary is clearly identified between samples ADB1701-0a and ADB1701-01 at Bishop Wash, ADB1703-04d and ADB1703-04e at Leaf 6, and ADB1802-06 and ADB1802-07 at 9789 (Figure 5, 6, S6; Table S8). Samples from above show the appearance of typical Paleocene *Momipites*, and low diversity assemblages often dominated by palm pollen (*Arecipites* spp.).

**Figure 6:**
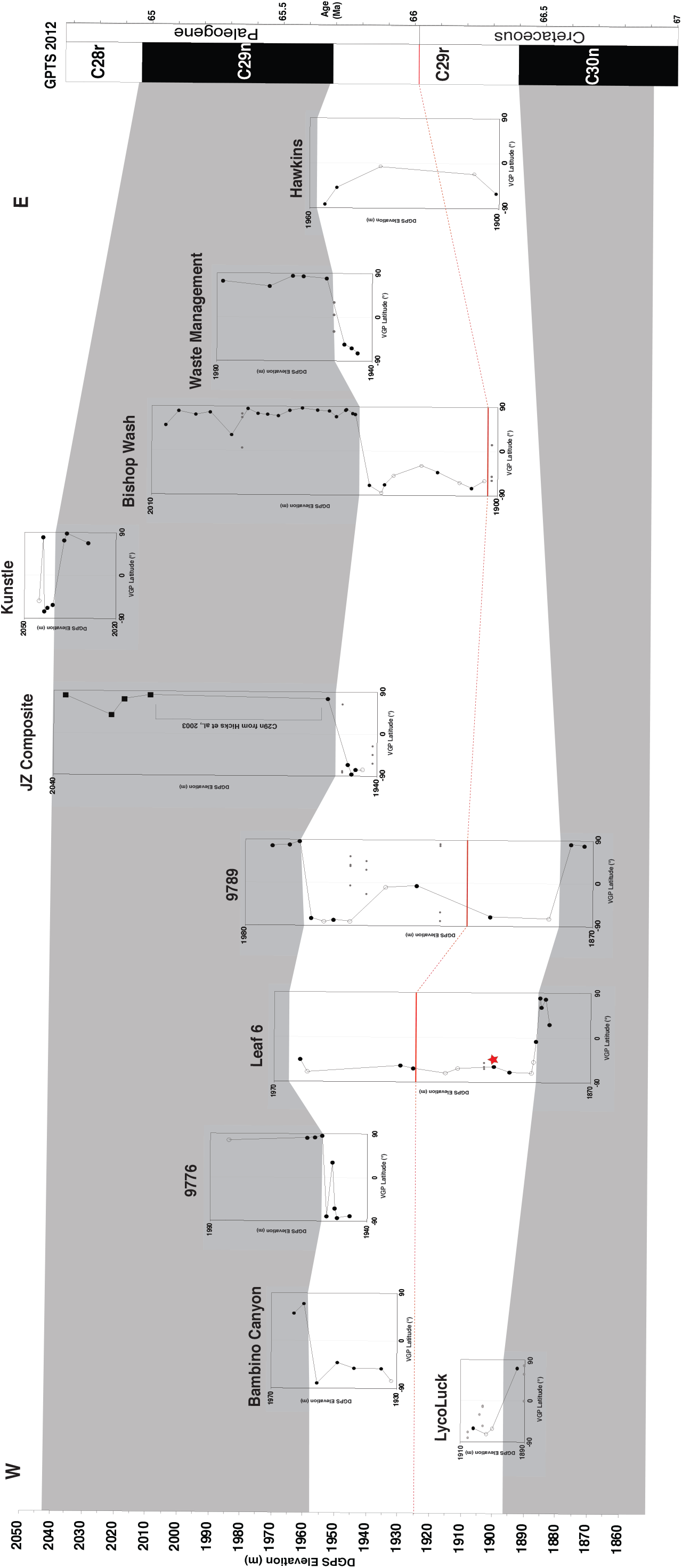
East to west fence diagram of Corral Bluffs with magnetostratigraphic sections plotted by elevation and correlated to the GPTS 2012 (Gradstein et al., 2012). Paleomagnetic section names labeled above respective VGP plots. Shaded regions correspond to the normal polarity Chrons 30n and C29n. Solid red line is the K–Pg boundary where determined through palynological analysis. Dashed red line represents inferred level of K–Pg across the field area. Bracketed zone in JZ section was previously identified as C29n by Hicks et al., (2003). Red star represents location of ^206^Pb/^238^U dated Feral Llama ash (KJ0958; 66.253 ± 0.031/0.045/0.084 Ma).

## DISCUSSION

### Correlations of Paleomagnetic Reversals to GPTS

Biostratigraphic indicators from the Corral Bluffs field area such as fragmentary dinosaur remains, numerous Late Cretaceous and early Paleocene leaf localities (Johnson et al., 2003), and mammal fossils tentatively assigned to the Pu2/3 NALMA stages (Eberle, 2003), constrain the D1 strata in this part of the Denver Basin to the time interval from the Latest Cretaceous to the early Paleocene, as demonstrated by the basin-wide chronostratigraphy of Clyde et al. (2016). The integrated palynological and high-precision U–Pb geochronological results reported here and tied to specific stratigraphic sections provide further support for this and help constrain the correlation of the magnetostratigraphic data to the GPTS and identify a precise position of the K–Pg boundary across the field area.

East of Jimmy Camp Creek, at Corral Bluffs, are the Leaf 6, 9789, and Bishop Wash sections, where the three palynologically determined K–Pg boundary sites from this study are located at approximately 1925, 1910, and 1904 m elevation respectively (Figure 5, 6, S8; Table S8). Fragmentary ceratopsian dinosaur material (DMNH Loc. 4195) was also identified at 1924.2 m in the Leaf 6 section. A ^206^Pb /^238^U age of 66.253 ± 0.031/0.045/0.084 Ma was obtained from zircons within the “Feral Llama” ash (KJ0958) at an elevation of 1900.4 m within the Leaf 6 section. An identical (within uncertainty) ^206^Pb /^238^U age of 66.252 ± 0.018/0.038/0.081 Ma was independently obtained for sample KJ1702 from the “Stock Tank” ash at a site ~1 km to the west in an area where no paleomagnetic data were collected (Figure 1, 5, S1). KJ1702 provides temporal constraints to nearby fossil localities in this area and supports our stratigraphic model indicating that the two ashes are actually from the same volcanic deposit, which allows for these localities to be directly correlated to the Leaf 6 section (Figure 1, S1).

Hicks et al. (2003) previously identified paleomagnetic reversals corresponding to the interval from C31n and C30n–C29r in the Jimmy Camp Creek section, the top of which overlaps with our Lyco-Luck section. The normal polarity zone within our Lyco-Luck section is interpreted to correspond to the upper portion C30n with a subsequent reversed interval interpreted as C29r (Figure 6). Further up section in Jimmy Camp Creek, the Bambino Canyon and 9776 sections record basal reverse intervals followed by normal polarity intervals (Figure 5 and 6). Due to their clear location stratigraphically higher than the Lyco-Luck section, the reverse polarity intervals are interpreted to correlate to C29r and the normal polarities to C29n.

At the base of the Leaf 6 and 9789 sections are normal polarity zones corresponding to C30n on the basis of their position below reversed intervals containing the K–Pg boundary correlating to C29r on the GPTS (Figure 6). The 9789 section records a second polarity reversal from reversed polarity to normal polarity near the top of the section and is correlated to the C29r–C29n reversal based on its position above the C30n–C29r reversal and the high density of sampling through the C29r interval exhibiting uniform polarity until the overlying C29r–C29n reversal (Figure 6).

Continuing eastward, the JZ composite section, comprising the JZB and Hillaire-West subsections (Figure S3; Table S1), overlaps with the JZ section of Hicks et al. 2003. We sampled within the gap between the highest reversed site and lowest normal site in the JZ section of Hicks to increase the resolution in this section. We also sampled higher via the Hillaire-West subsection. Our data agree with the determination of Hicks et al. (2003), that the transition from the reverse polarity zone to the normal polarity zone represents the C29r–C29n boundary based upon stratigraphic relationships (Figure 6). The gap in our sampling was previously shown to represent C29n by Hicks et al. (2003) and was not resampled as the C29n–C28r reversal was clearly higher than their JZ section.

The stratigraphically highest section in our field area is the Kunstle section to the east of the Hillaire-West subsection. This section records a normal polarity interval through the base of the section, followed by a reverse polarity interval and ending with mixed polarity at the very top (Figure S3). Traverses on marker beds stratigraphically tie the Kunstle section to the adjacent Hillaire-West subsection, indicating that the normal interval at the base of the Kunstle section likewise corresponds to C29n and the overlying reverse polarity zone correlates to C28r. Site CB1854 is a normal polarity site located above three reverse polarity sites and below another reverse polarity site, indicating that this site may either have a strong normal overprint or may have captured a short interval of instability in the Earth’s magnetic field at this time, possibly related to the next reversal. No slumping was apparent across the ridge where CB1854 was sampled and the sites are clearly in superposition.

The Bishop Wash section records the third palynologically determined K–Pg boundary near the base of the section at an elevation of ~1904 m, within a reversed interval assigned to C29r (Figure 6). The high resolution of sampling through this section indicates that the subsequent normal polarity zone must represent C29n.

The Waste Management Boathouse section records a paleomagnetic reversal from reversed to normal and given the stratigraphic position of the section relative to the Bishop Wash section it is interpreted that this reversal represents the C29r–C29n boundary (Figure 6). The westernmost section, the Hawkins section, is directly across highway 94 from the Waste Management Boathouse section and is characterized by a reversed zone throughout the entire section. The Hawkins section is stratigraphically below the adjacent Waste Management section where we identified the C29r–C29n reversal, therefore the reverse zone making up the entirety of this section is interpreted to be the C29r subchron (Figure 6).

The patterns of reversals observed at Corral Bluffs and Jimmy Camp Creek are correlated here to the interval C30n–C28r, with both the top and bottom of C29r observed within the 9789 section. The chron boundaries occur at very similar elevations across the entire field area, which agrees well with the nearly flat-lying beds observed in the field (Figure 6). The proportional thicknesses of the chron boundaries and the K–Pg boundary matches well with the GPTS 2012 timescale of Gradstein et al. (2012), which is supportive of the Corral Bluffs area preserving a near continuous depositional sequence that is not disrupted by significant unconformities.

### Correlations to Basin-Wide Magnetostratigraphic Framework

The pattern of paleomagnetic reversals identified at Corral Bluffs and Jimmy Camp Creek from this study and Hicks et al. (2003) is interpreted to represent the interval from Chrons C31n, and C30n–C28r. This represents the longest surface exposure in the Denver Basin for which there are chronostratigraphic constraints. The Kiowa core matches closely with these results as it likewise records Chrons C31n, and C30n–C28r (Hicks et al., 2003; Clyde et al., 2016). The short reversed interval C28r is not recorded in the thicker Castle pines core where it is thought to be cut out by a thick channel sandstone (Figure 7; Hicks et al., 2003). It also possible to correlate the three surface sections of Clyde et al. (2016) from the central part of the basin to our sites as these sections span the interval from C29r–C28n.

**Figure 7:**
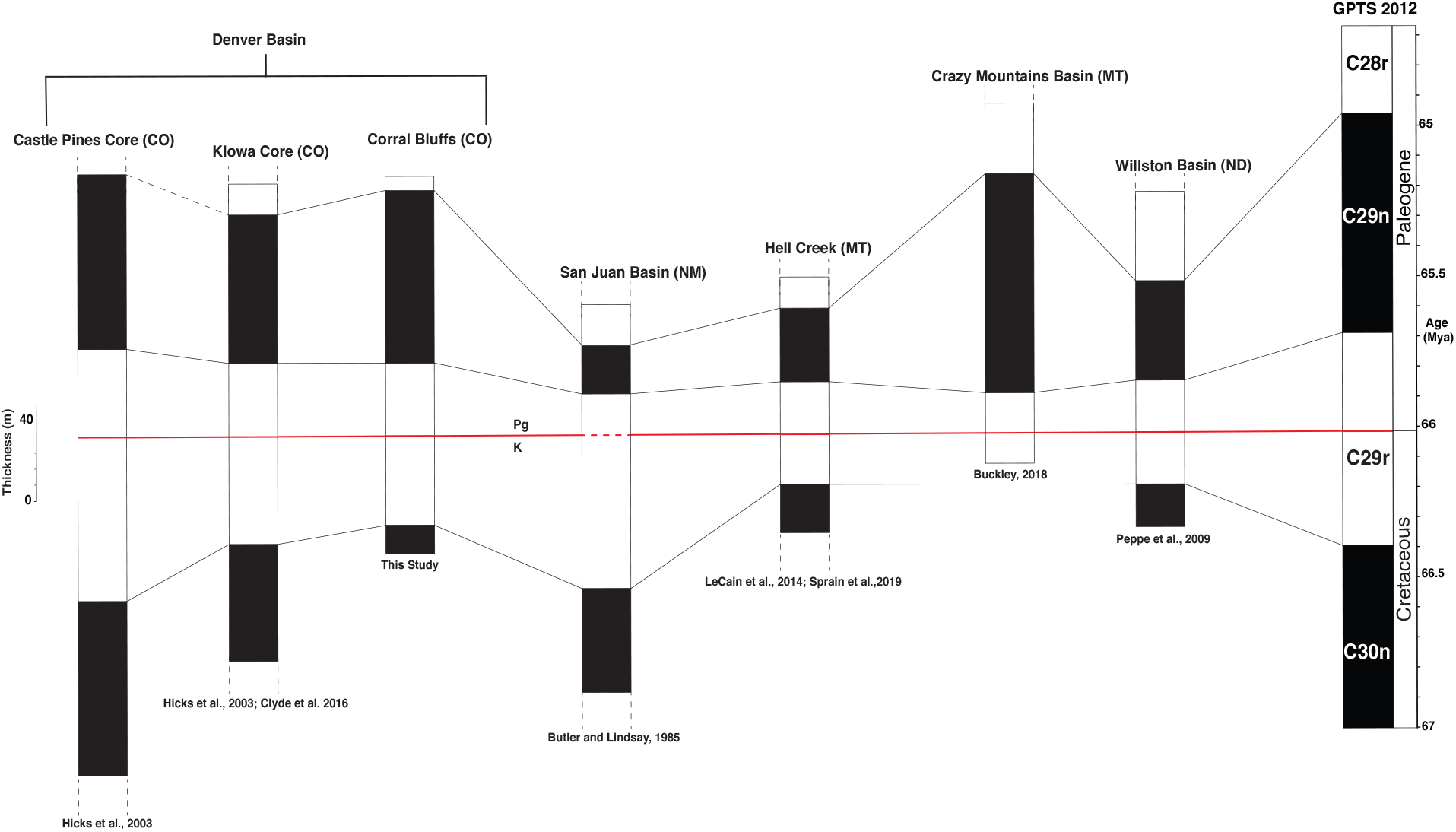
Magnetostratigraphic correlations of the K–Pg boundary interval (Top of Chron C30n–C28r) from the Castle Pines core, Kiowa core (Hicks et al., 2003) and Corral Bluffs to studies in the San Juan Basin (Butler and Lindsay, 1985; Peppe et al., 2013), Hell Creek region of Montana (LeCain et al., 2014; Sprain et al., 2018), and the Williston Basin in North Dakota (Peppe et al., 2009). Solid red line represents the K-Pg boundary as defined in the respective studies. K–Pg boundary dashed in San Juan Basin magnetostratigraphy represents uncertainty in the placement of boundary (Peppe et al. 2013). Dashed vertical lines indicate that sections continue further in those studies than is shown here. Dashed line from Castle Pines core to Kiowa indicates that the C29n–C28r boundary is not preserved in the Castle Pines Core and therefore the top of C29n is approximated. Solid black lines represent correlations of the paleomagnetic reversal boundaries across the studies to the GPTS 2012 (Gradstein et al., 2012). All sections plotted to same stratigraphic scale with approximate thickness scale shown (m).

### Paleontological implications

Given the reported provinciality of mammalian faunas during the Puercan (Lofgren et al., 2004), it is critical to the understanding of regional recovery dynamics to have independent age controls at key fossil localities in order to correlate coeval faunas across the Western Interior. It is now possible to tie our new magnetostratigraphy at Corral Bluffs to other key K-Pg boundary sites in the Western Interior (Butler and Lindsay, 1985; Swisher et al., 1993; Peppe et al., 2009; LeCain et al., 2014; Sprain et al., 2018; Buckley, 2018; Figure 7).

The Pu1 localities within the Denver Basin occur within C29r, including the Alexander locality which is thought to represent an intermediate between Pu1 and Pu2 faunas (Hicks et al., 2003; Eberle, 2003; Middleton et al. 2004). The Hell Creek region in Garfield and McCone Counties, Montana has yielded Pu1 and Pu3 mammal fossils and is among the best paleontologically sampled K–Pg boundary intervals with extensive chronostratigraphic controls (Swisher et al., 1993; Clemens, 2013; LeCain et al., 2014; Wilson, 2014; Sprain et al., 2018). Swisher et al. (1993) and LeCain et al. (2014) identified paleomagnetic reversals interpreted to represent C30r–C28n within the Hell Creek and Fort Union formations in Garfield and McCone Counties. Sprain et al. (2018) further refined the magnetostratigraphy of the Hell Creek region across 14 sections that are interpreted to span C30n through C29n. Pu1 assemblages within the Hell Creek region of Montana are located within C29r. The Hanna Basin, which is the only locality that preserves Lancian through Pu3 localities in direct superposition, does not as of yet have a magnetostratigraphy in which to place these localities (Lillegraven and Eberle, 1999).

The Pu2/3 fauna of Corral Bluffs described by Eberle (2003) were sampled from the C29n interval of Hicks et al. (2003), which corresponds to our JZ composite section and our new results support this determination. The Nacimiento Formation in the San Juan Basin is the type locality for the Puercan and is currently known to possess assemblages interpreted to correspond to Pu2 through Pu3 located within a normal polarity zone also interpreted as C29n (Butler and Lindsay, 1985; Williamson et al., 1996; Lofgren et al., 2004; Peppe et al., 2013). The Pu3 localities from the Hell Creek are also interpreted to lie within C29n (Sprain et al., 2018). Magnetostratigraphic results from the eastern Williston Basin in North Dakota complicate the associations observed elsewhere (Hunter and Archibald, 2002; Peppe et al., 2009). The Pu2 fauna from the PITA Flats locality is interpreted to occur within C29r based upon their proximity to the K–Pg boundary (Hunter and Archibald, 2002; Fox and Scott, 2011). The Pu2 Hiatt local fauna from Makoshika State Park in eastern Montana is located within C29r (Hunter et al., 1997). Pu2 assemblages from the Rav W-1 horizon of the Ravenscrag Formation in southwestern Saskatchewan similarly occur within C29r (Fox and Scott, 2011). Although it is possible that the variable occurrences of Pu2 within C29r and C29n could indicate increased provinciality across the Western Interior at that time as has been argued elsewhere, it is also possible that Pu2 simply begins within C29r and extends into C29n. Additional high-resolution chronostratigraphic constraints on sections that clearly record the Pu1/Pu2 boundary will be needed to resolve this.

## CONCLUSION

The magnetostratigraphic results reported here indicate that the Corral Bluffs area of the Denver Basin preserves a relatively continuous K–Pg boundary stratigraphic sequence spanning Chron C30n to C28r. These magnetostratigraphic results combined with three palynologically determined K–Pg boundary sites, and two independent ^206^Pb /^238^U ages for the same ash bed, sampled ~1 km apart, provide one of the very few precisely dated temporal frameworks for a fossil-bearing, continental, latest Cretaceous to early Paleocene boundary sequence. It is now possible to correlate the fossil localities in this area to contemporaneous sites in the basin, across the region, and around the globe using the GPTS (Gradstein et al., 2012; Ogg, 2012). Our results indicate that the Corral Bluffs Pu2/3 fauna lies within C29n, similar to what is found in the San Juan Basin of New Mexico and the Hell Creek area of Montana. The Corral Bluffs area represents an important candidate for future paleontological investigation due to this new high-resolution temporal framework and its previously noted presence of early Paleocene vertebrates and well-preserved Late Cretaceous and Early Paleocene megaflora. Corral Bluffs and Jimmy Camp Creek now represents the most extensive surface exposure in the Denver Basin for which there are chronostratigraphic controls, allowing for the opportunity to interpret fossil collections from this area in the context of K–Pg boundary extinction and recovery.

## Supporting information

Fuentes et al., Supplement

## ACKNOWLEDGEMENTS

Funding for this project was provided by a Geologic Society of America Graduate Student Research Grant and a Colorado Scientific Society Memorial Funds Grant to A.J.F and L. Levin Appel, M. Cleworth, Lyda Hill Philanthropies, David B. Jones Foundation, M. L. and S. R. Kneller, T. and K. Ryan, and J. Tucker as part of the Denver Museum of Nature & Science No Walls Community Initiative. Geochronology at MIT was supported by NSF grant EAR 0643158 to S. Bowring. The funding sources were not involved in the study design, data collection, interpretation, or submission for publication.

We thank Norwood Properties, City of Colorado Springs, Waste Management, J. Hawkins, J. Hilaire, J. Carner, W. Pendleton, the Bishop Family, and H. Kunstle for land access; R. M. Hess, R. Barclay, and S. Milito for help with field work; B. Snellgrove for logistics; and J. Johnson for discussion.

