## Supplementary material for "Constructing a Timescale of Biotic Recovery across the Cretaceous–Paleogene Boundary, Corral Bluffs, Denver Basin, Colorado": Fuentes et al., Supplement


**Supplementary Text**

**U–Pb Geochronology Analytical Methods**

**Supplementary Figures**

**Figure S1–Ash Beds Cross Section**

**Figure S2–Reversal Test**

**Figure S3–Virtual Geomagnetic Pole Plots**

**Figure S4–Paleomagnetic Pole**

**Figures S5–IRM Curves**

**Figure S6–Pollen Stratigraphic Sections Plots**

**Supplementary Tables**

**Table S1–Paleomagnetic Site Data**

**Table S2–Paleomagnetic Sample Data**

**Table S3–Isothermal Remanent Magnetization (IRM) Acquisition Data**

**Table S4–IRM Demagnetization Data**

**Table S5–U–Pb Isotopic Data for Sample KJ0958**

**Table S6–LA_ICPMS U-Pb Geochronologic Analyses for Sample KJ1702**

**Table S7–U-Th-Pb Isotopic Data for Sample KJ1702**

**Table S8–Pollen Counts and Sample Elevation Data**

**Supplementary References**

### Supplementary Text

### U–Pb Analytical Methods

#### *Sample KJ0958*

*Sample preparation*

Sample KJ0958 was processed and analyzed for U-Pb geochronology by the CA-IDTIMS method at the Massachusetts Institute of Technology Isotope Laboratory following the procedures outlined in Ramezani et al. (2011) that are largely similar to those applied to sample KJ1702. Zircon grains were selected based on their aspect ratios and morphologic characteristics with preference given to prismatic, multi-faceted crystals that contain glass (melt) inclusions parallel to their long axis. This selection strategy has proven highly effective in screening out reworked zircons, as well as those containing xenocrystic cores, and thus eliminating in many cases the need for extensive imaging and/or a-priori age screening by microbeam U-Pb techniques. Selected zircons for analysis were pre-treated by a chemical abrasion technique modified after Mattinson (2005), which involved thermal annealing in a furnace at 900°C for 60 hours, followed by partial dissolution in 29M HF at 210°C in high-pressure vessels for 12 hours. The 210°C leach temperature may result in significant loss of material (and thus lower analytical precision) in some zircons, but recent experiments (Widmann et al., 2019) have underscored its effectiveness in overcoming the subtle and often undetected Pb loss in zircon that may compromise the accuracy of high-precision U-Pb dates.

*CA-IDTIMS analysis*

Pre-treated and thoroughly fluxed/rinsed zircons were spiked with the EARTHTIME ET535 mixed $^{205}$Pb-$^{233}$U-$^{235}$U isotopic tracer (Condon et al. 2015; McLean et al. 2015) prior to complete dissolution and Pb and U purification by ion-exchange columns chemistry. Isotopic measurements were made on a VG Sector 54 multi-collector thermal ionization mass spectrometer equipped with a Daly photomultiplier ion counting system at MIT. Pb isotopes were measured as mono-atomic ions in a peak hopping mode on the ion counter and were corrected for a mass-dependent isotope fractionation of 0.24% ± 0.05% per atomic mass unit (2σ). U isotopes were measured as dioxide ions in a static mode using three Faraday collectors, while subjected to a within-run mass fractionation correction using the $^{233}$U/$^{235}$U ratio of the spike and a sample $^{238}U/^{235}U$ ratio of 137.818 ± 0.045 (Hiess et al., 2012), as well as an oxide correction based on an $^{18}O/^{16}O$ ratio of 0.00205 ± 0.00005.

Complete Pb and U isotopic data are given in Table S5 and the age results are illustrated on the date distribution plots of Figure 5. Data reduction, calculation of dates and propagation of uncertainties used the Tripoli and ET_Redux applications and algorithms (Bowring et al. 2011; McLean et al. 2011). The zircon $^{206}Pb/^{238}U$ dates were corrected for initial $^{230}Th$ disequilibrium based on a magma Th/U ratio of 2.8 ± 1.0 (2σ). The sample date was calculated based on the weighted mean $^{206}Pb/^{238}U$ date of all seven zircon analyses without any outlier exclusions and is reported at 95% confidence level in the ±*X*/*Y*/*Z* Ma format. The use of ET U-Pb tracer allows direct comparisons between both ash bed ages reported here and with the comprehensive Denver Basin chronostratigraphy of Clyde et al. (2016) without the need to consider the external (systematic) uncertainties.

***Sample KJ1702***

*Sample preparation*

Sample KJ1702 was processed and analysed using LA-ICPMS and CA-IDTIMS at Boise State University Isotope Geology Laboratory. Zircon crystals were separated by conventional density and magnetic methods. The entire zircon separate was placed in a muffle furnace at 900°C for 60 hours in quartz beakers to anneal minor radiation damage; annealing enhances cathodoluminescence (CL) emission (Nasdala et al., 2002), promotes more reproducible interelement fractionation during laser ablation inductively coupled plasma mass spectrometry (LA-ICPMS) (Allen and Campbell, 2012), and prepares the crystals for subsequent chemical abrasion (Mattinson, 2005). Following annealing, individual grains were hand-picked and mounted, polished and imaged by cathodoluminence (CL) on a scanning electron microscope. From these compiled images, the location of spot analyses for LA-ICPMS were selected.

*LA-ICPMS analysis*

LA-ICPMS analysis utilized an X-Series II quadrupole ICPMS and New Wave Research UP-213 Nd:YAG UV (213 nm) laser ablation system. In-house analytical protocols, standard materials, and data reduction software were used for acquisition and calibration of U-Pb dates and a suite of high field strength elements (HFSE) and rare earth elements (REE). Zircon was ablated with a laser spot of 25 μm wide using fluence and pulse rates of ~5 J/cm$^2$ and 10 Hz, during a 45 second analysis (15 sec gas blank, 30 sec ablation) that excavated a pit ~25 μm deep. Ablated material was carried by a 1.2 L/min He gas stream to the nebulizer flow of the plasma. Quadrupole dwell times were 5 ms for Si and Zr, 200 ms for $^{49}$Ti and $^{207}$Pb, 80 ms for $^{206}$Pb, 40 ms for $^{202}$Hg, $^{204}$Pb, $^{208}$Pb, $^{232}$Th, and $^{238}$U and 10 ms for all other HFSE and REE; total sweep duration is 950 ms. Background count rates for each analyte were obtained prior to each spot analysis and subtracted from the raw count rate for each analyte. For concentration calculations, background-subtracted count rates for each analyte were internally normalized to $^{29}$Si and calibrated with respect to NIST SRM-610 and -612 glasses as the primary standards. Ablations pits that appear to have intersected glass or mineral inclusions were identified based on Ti and P signal excursions, and associated sweeps were generally discarded. U-Pb dates from these analyses are considered valid if the U-Pb ratios appear to have been unaffected by the inclusions. Signals at mass 204 were normally indistinguishable from zero following subtraction of mercury backgrounds measured during the gas blank (<100 cps $^{202}$Hg), and thus dates are reported without common Pb correction. Rare analyses that appear contaminated by common Pb were rejected based on mass 204 greater than baseline. Temperature was calculated from the Ti-in-zircon thermometer (Watson et al., 2006). Because there are no constraints on the activity of $TiO_2$ in the source rocks, an average value in crustal rocks of 0.8 was used.

For U-Pb and $^{207}$Pb/$^{206}$Pb dates, instrumental fractionation of the background-subtracted ratios was corrected and dates were calibrated with respect to interspersed measurements of zircon standards and reference materials. The primary standard Plešovice zircon (Sláma et al., 2008) was used to monitor time-dependent instrumental fractionation based on two analyses for every 10 analyses of unknown zircon. A polynomial fit to the primary standard analyses versus time yields each sample-specific fractionation factor. A secondary bias correction is subsequently applied to unknowns on the basis of the residual age bias as a function of radiogenic Pb count rate in standard materials including Seiland, Zirconia, and Plesovice zircon. A polynomial fit to the secondary standard analyses with Pb count rate yields each sample-specific bias correction. Radiogenic isotope ratio and age error propagation for all analyses includes uncertainty contributions from counting statistics and background subtraction. Because the non-detrital zircon analyses are not interpreted individually, uncertainties from the standard calibrations are not propagated into the errors on each date. These uncertainties are the local standard deviations of the polynomial fits to the interspersed primary standard measurements versus time for the time-dependent, relatively larger U/Pb fractionation factor, and the standard errors of the means of the consistently time-invariant and smaller $^{207}Pb/^{206}Pb$ fractionation factor. These uncertainties are 0.96-1.25% (2σ) for $^{206}Pb/^{238}U$ and 0.56-0.77% (2σ) for $^{207}Pb/^{206}Pb$. Additional details of methodology and reproducibility are reported in Rivera et al. (2013).

*CA-IDTIMS analysis*

Following CL imaging, zircon grains chosen for IDTIMS analysis were subjected to a modified version of the chemical abrasion method of Mattinson (2005). Single grains were then transferred to 3 ml Teflon PFA beakers and loaded into 300 µl Teflon PFA microcapsules. Microcapsules were placed in a large-capacity Parr vessel and the grains partially dissolved in 120 µl of 29 M HF for 12 hours at 190°C. The contents of the microcapsules were returned to 3 ml Teflon PFA beakers, HF removed, and the residual grains immersed in 3.5 M $HNO_3$, ultrasonically cleaned for an hour, and fluxed on a hotplate at 80°C for an hour. The $HNO_3$ was removed and grains were rinsed twice in ultrapure $H_2O$ before being reloaded into the 300 µl Teflon PFA microcapsules (rinsed and fluxed in 6 M HCl during sonication and washing of the grains) and spiked with the EARTHTIME mixed $^{233}U$-$^{235}U$-$^{205}Pb$ tracer solution. Zircon was dissolved in Parr vessels in 120 µ l of 29 M HF with a trace of 3.5 M $HNO_3$ at 220°C for 48 hours, dried to fluorides, and re-dissolved in 6 M HCl at 180°C overnight. U and Pb were separated from the zircon matrix using an HCl-based anion-exchange chromatographic procedure (Krogh, 1973), eluted together and dried with 2 µl of 0.05 N $H_3PO_4$.

Pb and U were loaded on a single outgassed Re filament in 5 µl of a silica-gel/phosphoric acid mixture (Gerstenberger and Haase, 1997), and U and Pb isotopic measurements made on a GV IsotopX Phoenix multicollector thermal ionization mass spectrometers equipped with an ion-counting Daly detector. Pb isotopes were measured by peak-jumping all isotopes on the Daly detector for 100 to 160 cycles, and corrected for 0.16 ± 0.03%/a.m.u. (1 sigma error) mass fractionation. Transitory isobaric interferences due to high-molecular weight organics, particularly on $^{204}Pb$ and $^{207}Pb$, disappeared within approximately 30 cycles, while ionization efficiency averaged $10^4$ cps/pg of each Pb isotope. Linearity (to ≥1.4 x $10^6$ cps) and the associated deadtime correction of the Daly detector were monitored by repeated analyses of

NBS982, and have been constant since installation. Uranium was analyzed as $UO_2^+$ ions in static Faraday mode on $10^{11}$ ohm $10^{12}$ ohm resistors for 200-300 cycles, and corrected for isobaric interference of $^{233}U^{18}O^{16}O$ on $^{235}U^{16}O^{16}O$ with an $^{18}O/^{16}O$ of 0.00206. Ionization efficiency averaged 20 mV/ng of each U isotope. U mass fractionation was corrected using the known $^{233}U/^{235}U$ ratio of the EARTHTIME tracer solution.

CA-IDTIMS U-Pb dates and uncertainties were calculated using the algorithms of Schmitz and Schoene (2007), EARTHTIME ET535 tracer solution (Condon et al., in prep.; McLean et al., in prep.) with calibration of $^{235}U/^{205}Pb$ = 100.233, $^{233}U/^{235}U$ = 0.99506, and $^{205}Pb/^{204}Pb$ = 11268, and U decay constants recommended by Jaffey et al. (1971). $^{206}Pb/^{238}U$ ratios and dates were corrected for initial $^{230}Th$ disequilibrium using a Th/U[magma] = 3.0 ± 0.3, resulting in an increase in the $^{206}Pb/^{238}U$ dates of ~0.09 Ma. All common Pb in analyses was attributed to laboratory blank and subtracted based on the measured laboratory Pb isotopic composition and associated uncertainty. U blanks are estimated at 0.013 pg.

Weighted mean $^{206}Pb/^{238}U$ dates were calculated from equivalent dates using Isoplot 3.0 (Ludwig, 2003). Errors on the weighted mean dates are given as ± x / y / z, where x is the internal error based on analytical uncertainties only, including counting statistics, subtraction of tracer solution, and blank and initial common Pb subtraction, y includes the tracer calibration uncertainty propagated in quadrature, and z includes the $^{238}U$ decay constant uncertainty propagated in quadrature. Internal errors should be considered when comparing our dates with $^{206}Pb/^{238}U$ dates from other laboratories that used the same EARTHTIME tracer solution or a tracer solution that was cross-calibrated using EARTHTIME gravimetric standards. Errors including the uncertainty in the tracer calibration should be considered when comparing our dates with those derived from other geochronological methods using the U-Pb decay scheme (e.g., laser ablation ICPMS). Errors including uncertainties in the tracer calibration and $^{238}U$ decay constant (Jaffey et al., 1971) should be considered when comparing our dates with those derived from other decay schemes (e.g., $^{40}Ar/^{39}Ar$, $^{187}Re$-$^{187}Os$). Errors for weighted mean dates and dates from individual grains are given at 2σ

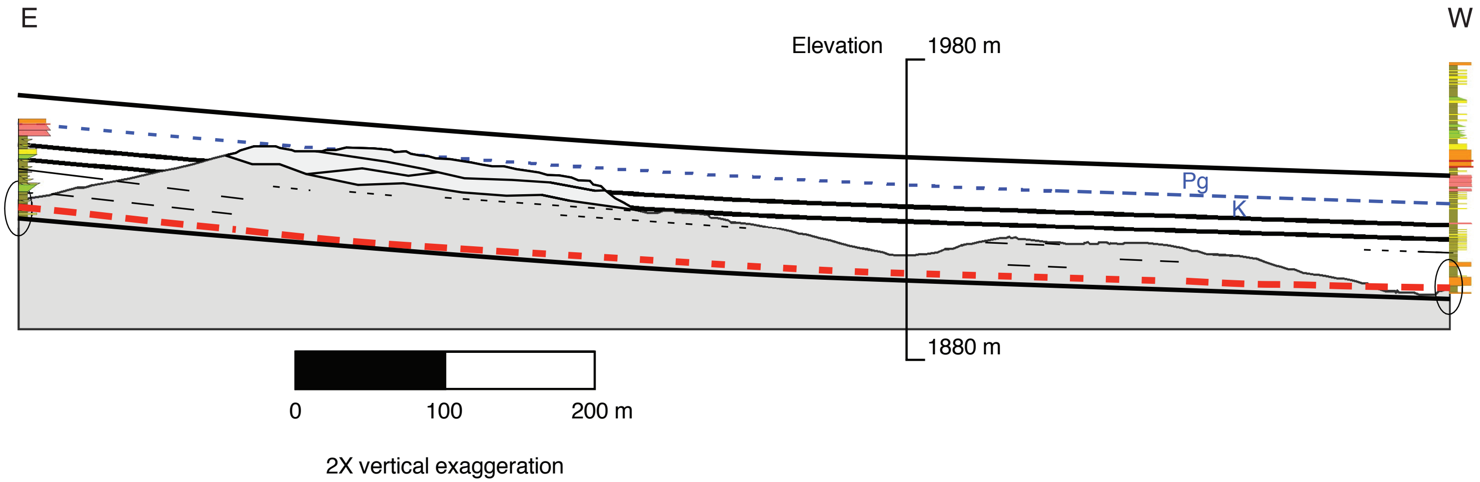


**Figure S1:** Schematic cross-section, Stock Tank section (left/west) to LEAF6 section (right/east). The ground-surface profile is from Google Earth. The cross-section shows a top Cretaceous interval of floodplain and fluvial sedimentary rocks (darker gray). Andesite-bearing pebbly sandstones, in a stack of fluvial channels (lighter gray), cap ridge on left. Multiple lignite/coal seams, each up to 25 cm thick, occur in the interval between the lower pair of black lines. The upper pair of black lines highlights an interval of arkosic ("granite wash") pebbly sandstones in isolated/dispersed channels; this interval brackets the pollen-indicated K-Pg boundary in the LEAF6 section. The dashed red line, near the base of the coaly interval, highlights a volcanic ash bed or beds in the Stock Tank and LEAF6 sections. Both occurrences are 1-2 cm of gray ash sandwiched in 10-20 cm of lignite. KJ1702 (Stock Tank) is ~30 m higher than KJ0958 (LEAF6) at a horizontal offset of ~940 m. Structural dip projections, similarities in stratigraphy, and a tight overlap in U-Pb age dates support the two samples coming either from a single ash or from two separate ashes that are just close in time.

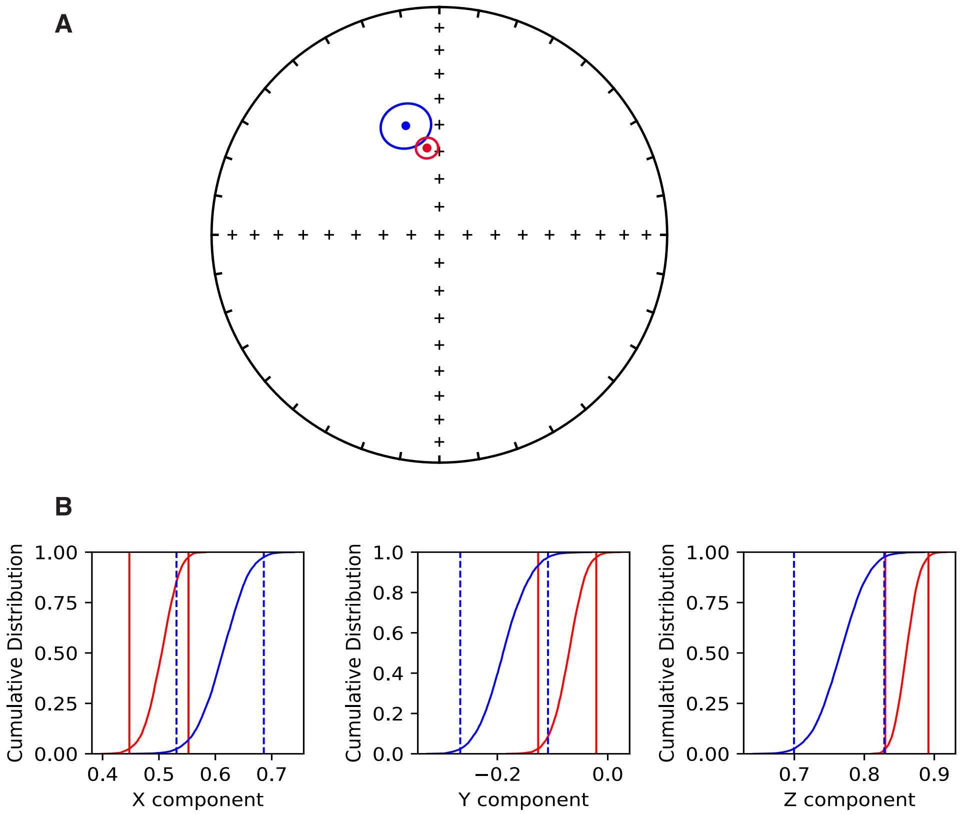


**Figure S2: A.** Mean ChRM directions of alpha normal polarity sites (red) and alpha reversed polarity sites flipped to antipodal direction (blue) with overlapping α95 confidence ellipses. **B.** Results of the parametric bootstrap reversal test of Tauxe et al. (2018) showing overlap of X, Y, and Z components in Cartesian coordinates.

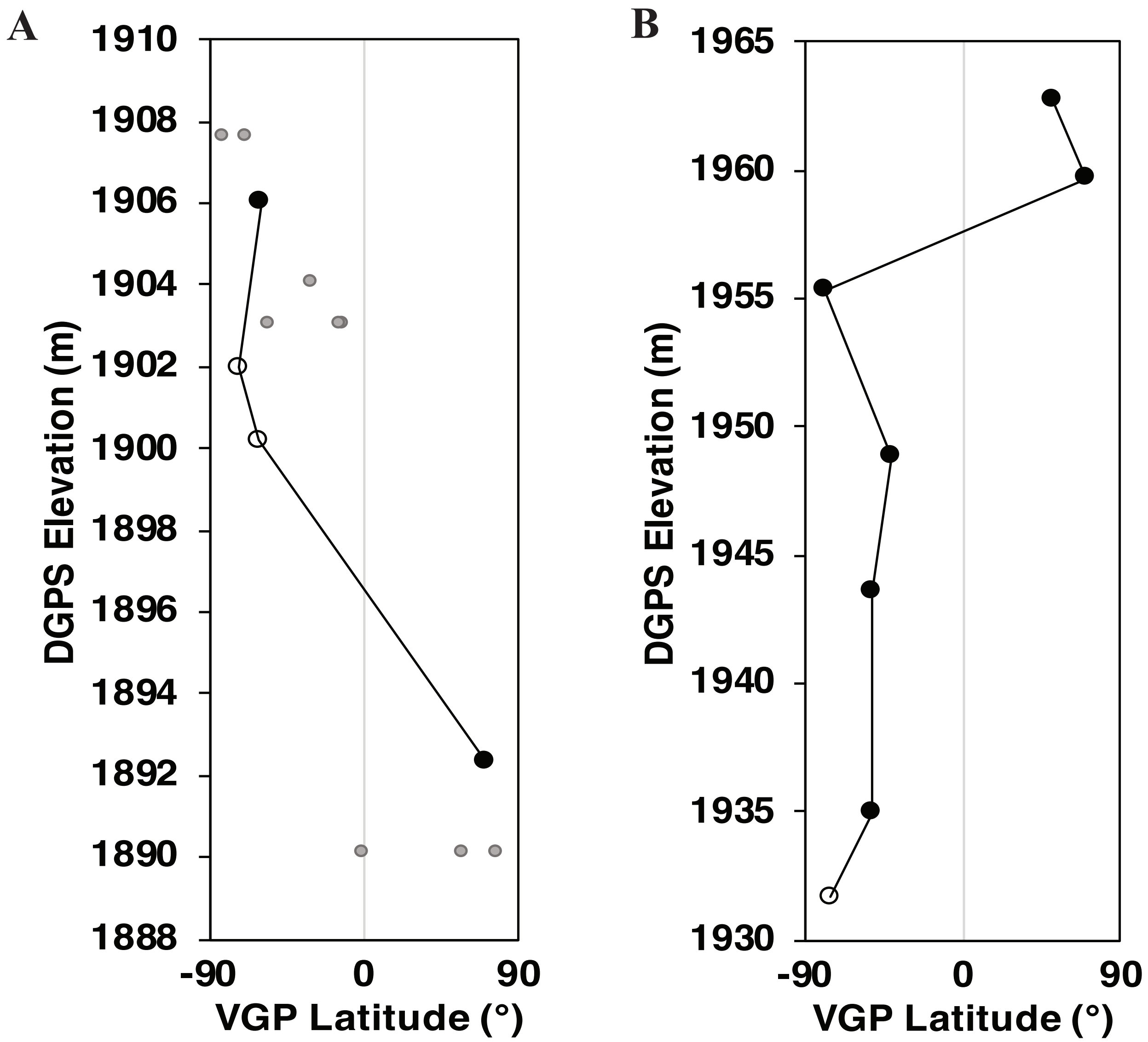

A
B
DGPS Elevation (m)
1910
1908
1906
1904
1902
1900
1898
1896
1894
1892
1890
1888
1965
1960
1955
1950
1945
1940
1935
1930
-90
0
90
VGP Latitude (°)

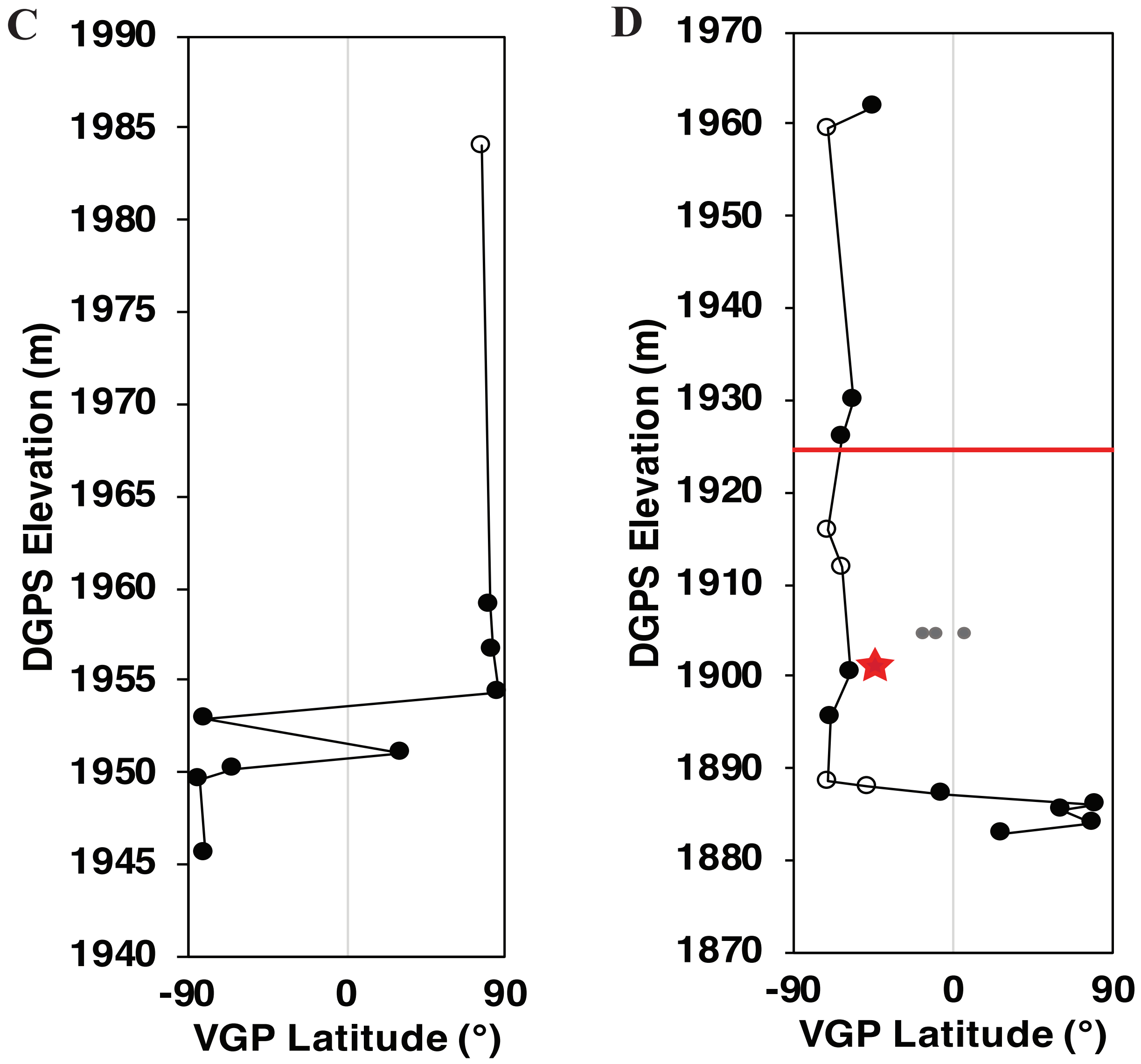
C
1990
1985
1980
1975
1970
1965
1960
1955
1950
1945
1940
DGPS Elevation (m)
-90
0
90
VGP Latitude (°)
D
1970
1960
1950
1940
1930
1920
1910
1900
1890
1880
1870
DGPS Elevation (m)
-90
0
90
VGP Latitude (°)

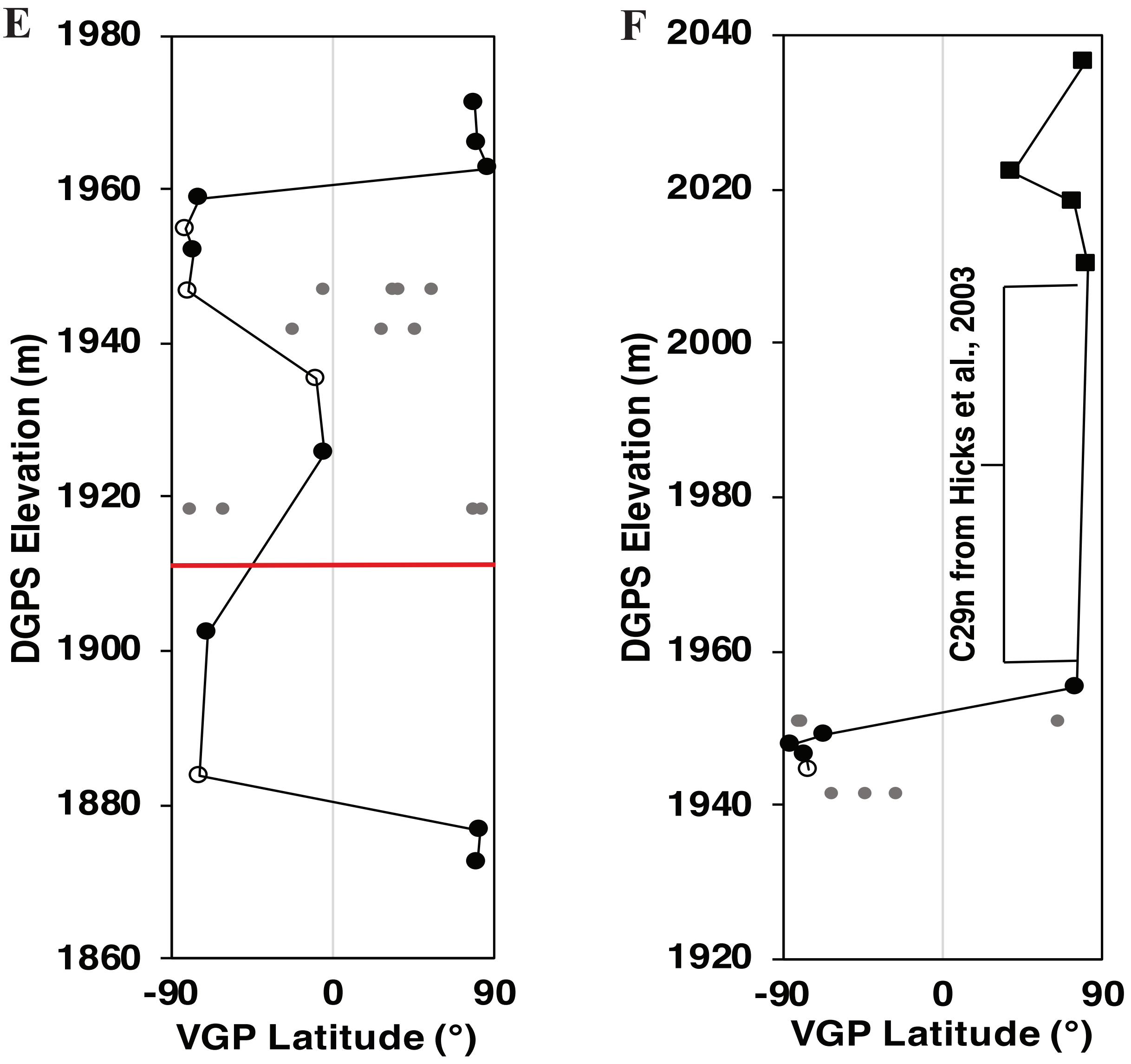

E
1980
1960
1940
1920
1900
1880
1860
DGPS Elevation (m)
-90
0
90
VGP Latitude (°)
F
2040
2020
2000
1980
1960
1940
1920
DGPS Elevation (m)
C29n from Hicks et al., 2003
-90
0
90
VGP Latitude (°)

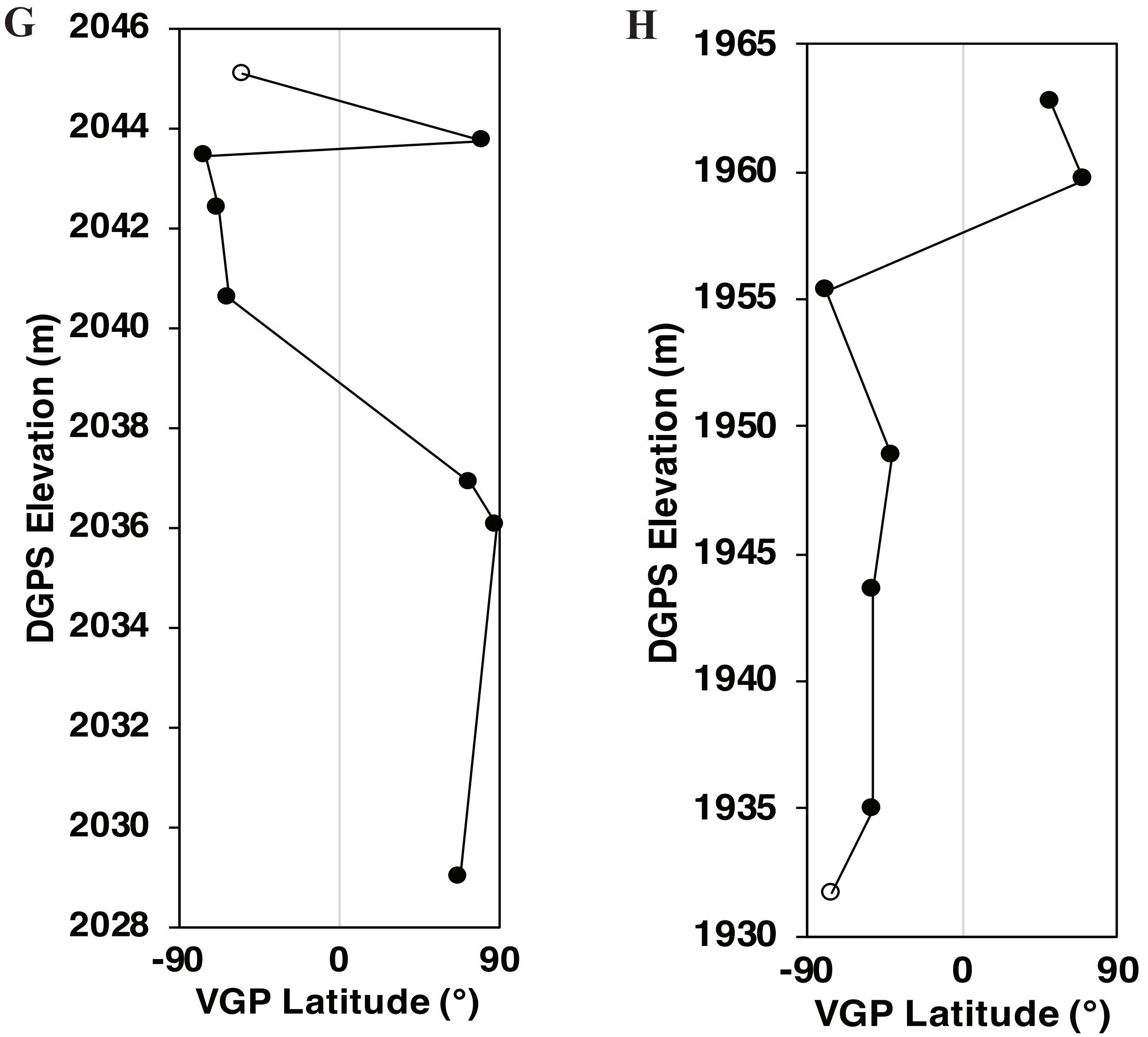
G
2046
2044
2042
2040
2038
2036
2034
2032
2030
2028
DGPS Elevation (m)
-90
0
90
VGP Latitude (°)
H
1965
1960
1955
1950
1945
1940
1935
1930
DGPS Elevation (m)
-90
0
90
VGP Latitude (°)

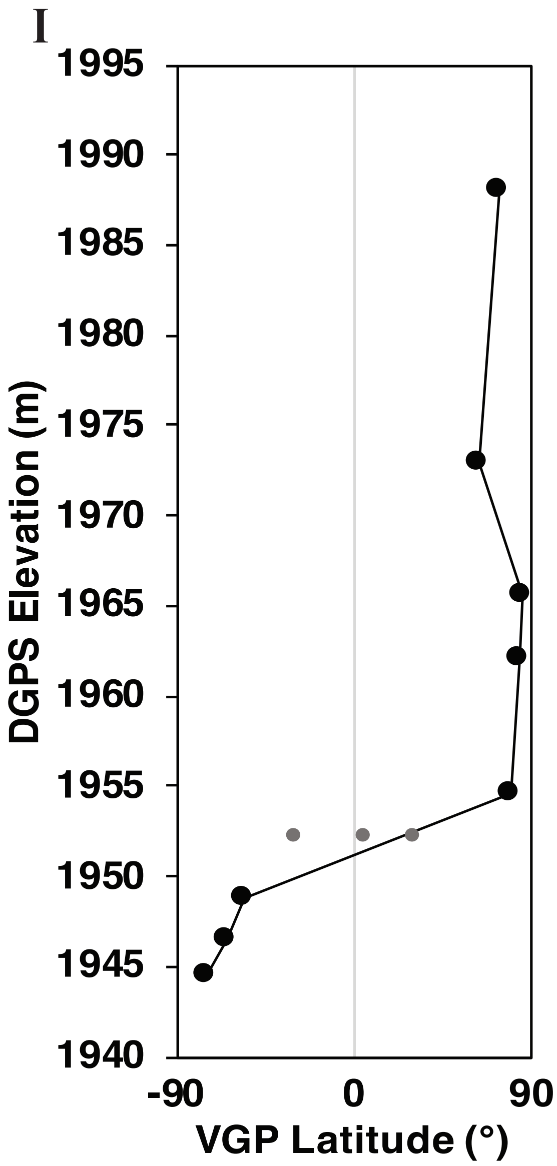


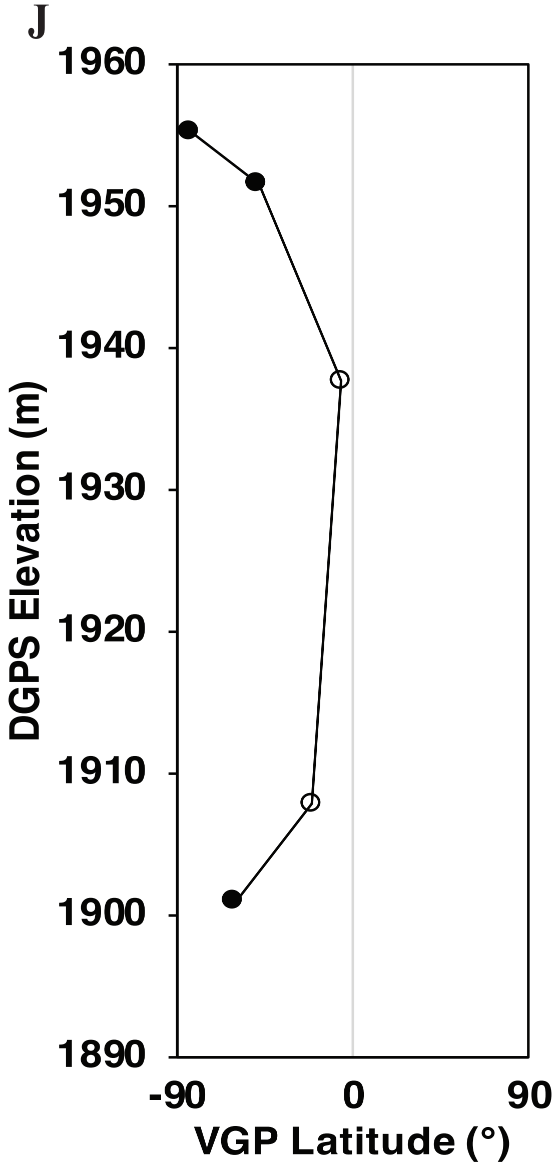


**Figure S3: A-J:** VGP plots for the Lyco Luck, Bambino Canyon, 9776, Leaf 6, 9789, JZ composite, Kunstle, Bishop Wash, Waste Management, and Hawkins sections. Paleomagnetic sites plotted on DGPS elevation (m). Closed circles alpha-level sites, open circles beta level sites. Small, grey circles are samples from sites that did not pass the Watson test. Red line in Leaf 6, 9789, and Bishop Wash sections indicates level of K-Pg boundary within those sections. Red star represents location of the U–Pb dated Feral Llama ash (KJ0958).

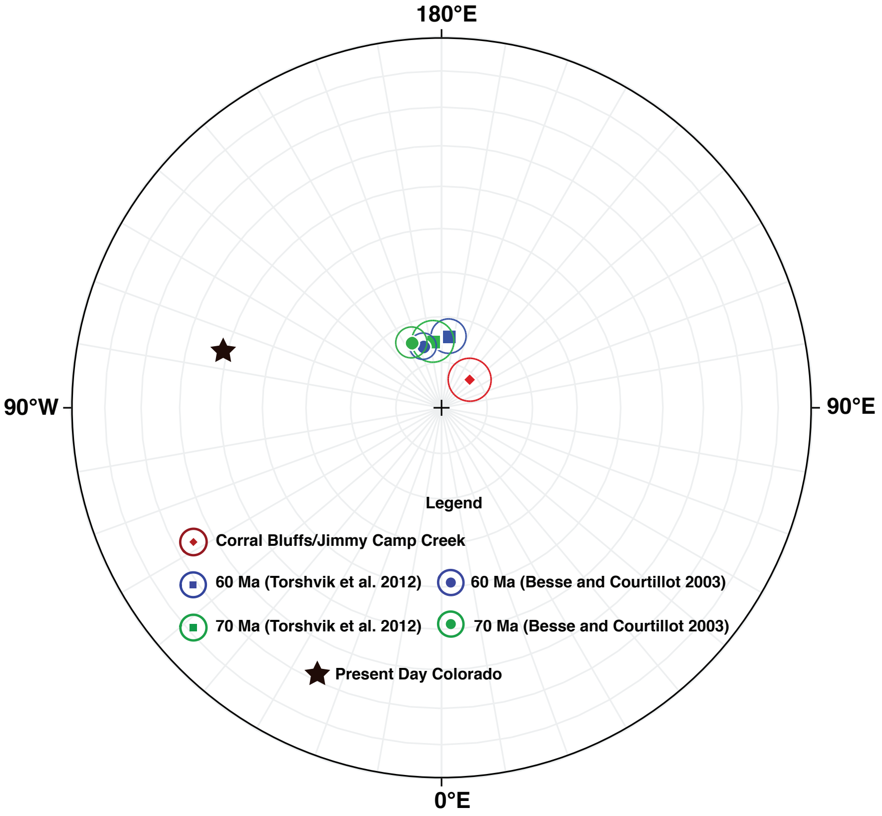


**Figure S4:** Polar equal area projection showing the paleomagnetic pole for this study (red diamond) plotted with its α95 confidence interval. 70Ma (green) and 60Ma (blue) paleomagnetic poles and corresponding α95 confidence intervals for Laurentia (North America and Greenland) from Besse and Courtillot (2003) (circles) and Torsvik et al. (2012) (squares). Black star represents the location of present-day Colorado relative to poles.

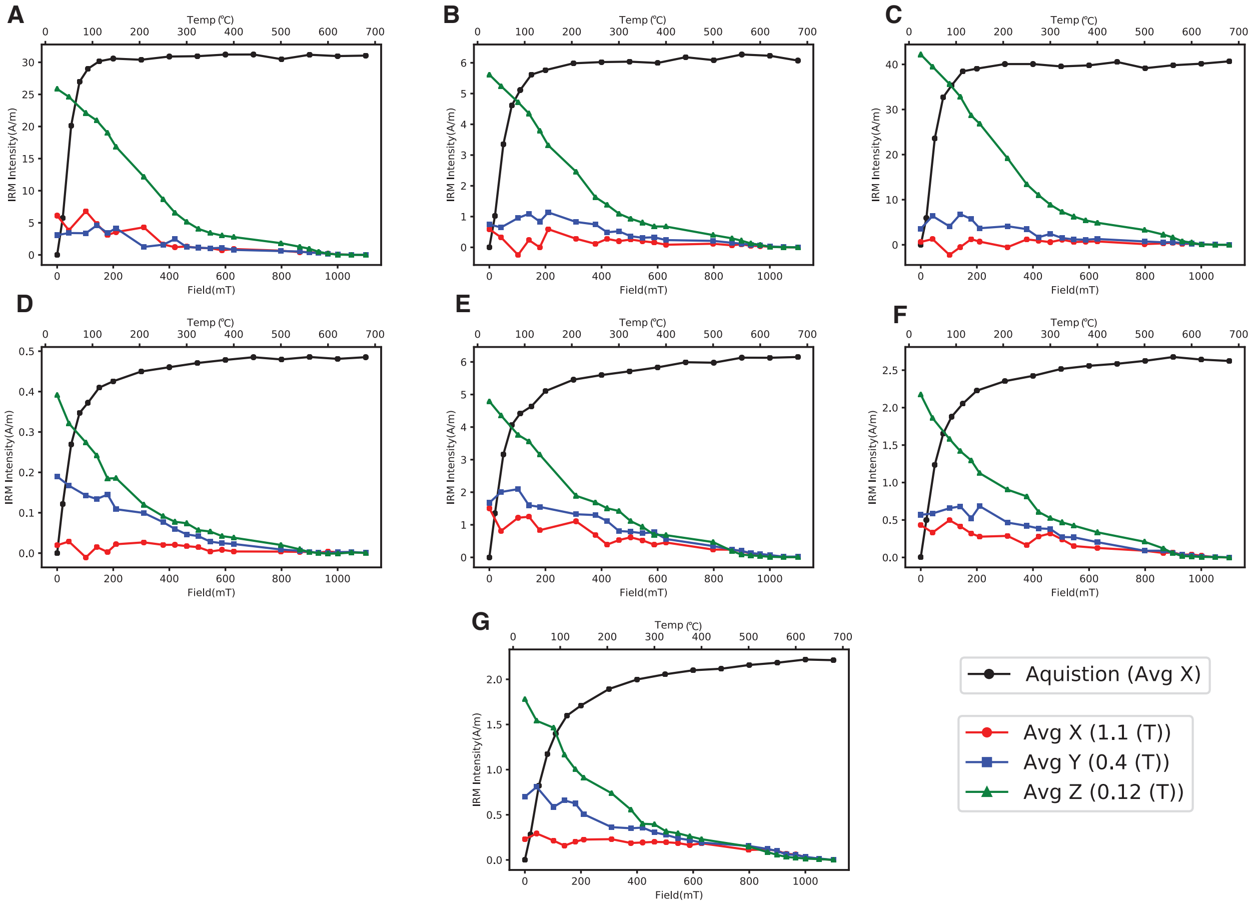


**Figure S5:** IRM acquisition and three-axis demagnetization curves for samples **A:** CB1703A, **B:** CB1711C, **C:** CB1734B, **D:** CB1821D, **E:** CB1824B, **F:** CB1827B, **G:** CB1869B.

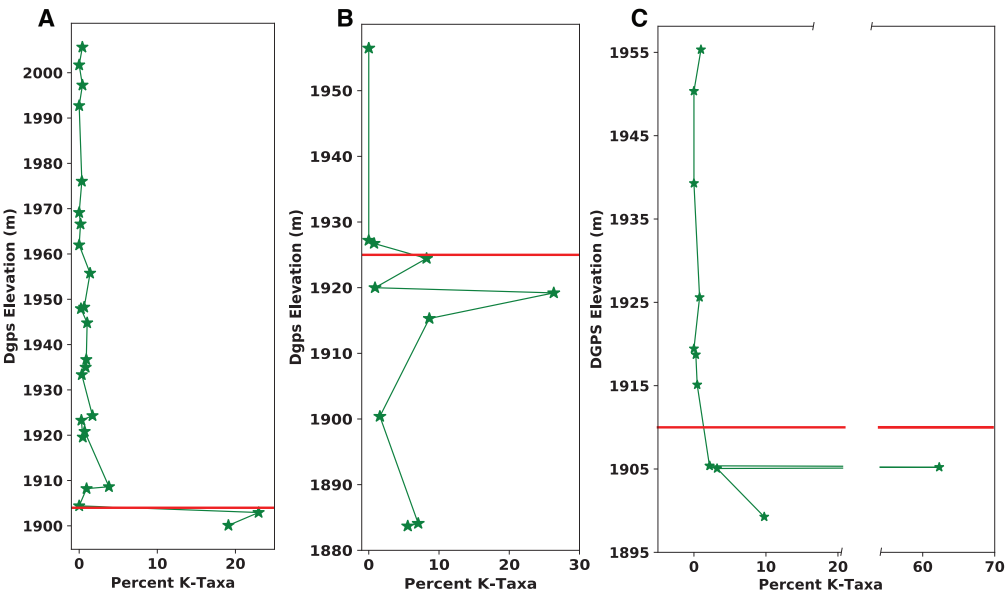


**Figure S6:** Cretaceous pollen counts as a percentage of total palynomorph counts (Percent K-Taxa) per sample plotted by DGPS elevation (m). **A.** Bishop Wash section. **B.** Leaf 6 section **C.** 9789 section. Red line represents placement of the K-Pg boundary in each section.

| Site | Section | Elev (m) | Lon (º) | Lat (º) | α/β | PCA/Fish/GC | Dec (°) | Inc (°) | α95 (°) | n | R | k | VGP Lat (º) | VGP Lon (º) |
|---|---|---|---|---|---|---|---|---|---|---|---|---|---|---|
| CB1701 | Bishop Wash | 1904.3 | -104.59561 | 38.84592 | β | 0,0,4 | 207.0 | -34.2 | 42.8 | 4 | 3.94 | 18.1 | -59.2 | 198.3 |
| CB1702 | Bishop Wash | 1908.2 | -104.59324 | 38.84781 | α | 0,3,0 | 194.1 | -68.5 | 44.7 | 3 | 2.77 | 8.7 | -73.8 | 108.0 |
| CB1703 | Bishop Wash | 1912.1 | -104.59224 | 38.84773 | β | 2,0,1 | 157.3 | -36.4 | 34.5 | 3 | 2.92 | 18.6 | -63.0 | 308.4 |
| CB1704 | Bishop Wash | 1919.0 | -104.59244 | 38.84765 | α | 1,2,0 | 139.5 | -13.8 | 29.9 | 3 | 2.89 | 18.1 | -41.6 | 315.0 |
| CB1705 | Bishop Wash | 1924.2 | -104.59190 | 38.85159 | β | 0,0,3 | 106.6 | -45.4 | NaN | 3 | 2.96 | 12.7 | -28.8 | 358.1 |
| CB1706 | Bishop Wash | 1933.0 | -104.59212 | 38.85552 | β | 0,0,3 | 149.1 | -18.1 | NaN | 3 | 2.89 | 4.7 | -49.5 | 306.8 |
| CB1707 | Bishop Wash | 1935.8 | -104.59144 | 38.85562 | α | 0,3,0 | 164.5 | -37.6 | 14.3 | 3 | 2.97 | 75.7 | -67.7 | 296.7 |
| CB1708 | Bishop Wash | 1936.9 | -104.59077 | 38.85591 | β | 0,0,4 | 176.9 | -53.5 | 6.5 | 3 | 3.00 | 10403.3 | -84.6 | 283.9 |
| CB1709 | Bishop Wash | 1940.6 | -104.59073 | 38.85586 | α | 0,3,0 | 169.9 | -37.1 | 58.6 | 3 | 2.64 | 5.5 | -69.9 | 283.9 |
| CB1875 | Bishop Wash | *1945.2* | -104.59073 | 38.85582 | α | 2,1,0 | 352.7 | 43.5 | 26.2 | 3 | 2.91 | 23.2 | 75.2 | 102.2 |
| CB1710 | Bishop Wash | 1946.1 | -104.59071 | 38.85582 | α | 3,0,0 | 343.2 | 57.4 | 10.0 | 3 | 2.99 | 153.1 | 76.8 | 167.0 |
| CB1711 | Bishop Wash | 1948.1 | -104.59068 | 38.85579 | α | 3,0,0 | 1.5 | 62.0 | 12.9 | 3 | 2.98 | 92.7 | 85.5 | 269.2 |
| CB1712 | Bishop Wash | 1948.3 | -104.58986 | 38.85611 | α | 3,0,0 | 351.3 | 58.3 | 18.7 | 3 | 2.96 | 44.7 | 83.2 | 169.4 |
| CB1713 | Bishop Wash | 1951.3 | -104.58982 | 38.85627 | α | 3,0,0 | 343.8 | 71.3 | 10.3 | 3 | 2.99 | 144.7 | 69.8 | 228.4 |
| CB1714 | Bishop Wash | 1953.5 | -104.58988 | 38.85628 | α | 3,0,0 | 8.9 | 53.4 | 15.7 | 3 | 2.97 | 62.8 | 81.4 | 17.3 |
| CB1715 | Bishop Wash | 1957.3 | -104.58999 | 38.85646 | α | 3,0,0 | 355.0 | 63.1 | 8.4 | 3 | 2.99 | 216.1 | 83.2 | 224.2 |
| CB1716 | Bishop Wash | 1962.1 | -104.58988 | 38.85653 | α | 3,0,0 | 3.8 | 58.3 | 2.8 | 3 | 3.00 | 1984.2 | 87.0 | 341.1 |
| CB1717 | Bishop Wash | 1966.1 | -104.58981 | 38.85663 | α | 3,0,0 | 352.5 | 62.9 | 11.9 | 3 | 2.98 | 108.9 | 82.2 | 212.1 |
| CB1718 | Bishop Wash | 1969.7 | -104.58977 | 38.85684 | α | 3,0,0 | 336.4 | 56.5 | 25.1 | 3 | 2.92 | 25.1 | 71.4 | 167.3 |
| CB1719 | Bishop Wash | 1973.1 | -104.58972 | 38.85697 | α | 3,0,0 | 340.2 | 54.2 | 9.8 | 3 | 2.99 | 158.6 | 73.7 | 157.0 |
| CB1720 | Bishop Wash | 1976.2 | -104.58970 | 38.85702 | α | 3,0,0 | 343.0 | 65.4 | 7.2 | 3 | 2.99 | 293.6 | 75.0 | 205.8 |
| CB1721 | Bishop Wash | 1979.4 | -104.58959 | 38.85709 | α | 3,0,0 | 353.8 | 58.3 | 11.0 | 3 | 2.98 | 126.5 | 85.2 | 168.4 |
| CB1723 | Bishop Wash | 1984.5 | -104.58900 | 38.85736 | α | 0,3,0 | 281.1 | 60.0 | 58.3 | 3 | 2.64 | 5.5 | 31.6 | 194.9 |
| CB1724 | Bishop Wash | 1991.4 | -104.58840 | 38.85725 | α | 3,0,0 | 12.9 | 54.3 | 41.4 | 3 | 2.80 | 9.9 | 78.9 | 2.9 |
| CB1725 | Bishop Wash | 1995.9 | -104.58768 | 38.85799 | α | 3,0,0 | 340.9 | 51.1 | 14.3 | 3 | 2.97 | 75.8 | 73.0 | 146.8 |
| CB1726 | Bishop Wash | 2001.3 | -104.58770 | 38.85809 | α | 3,0,0 | 347.9 | 56.8 | 11.2 | 3 | 2.98 | 121.8 | 80.3 | 160.2 |
| CB1727 | Bishop Wash | 2005.5 | -104.58768 | 38.85823 | α | 1,2,0 | 309.4 | 59.2 | 9.6 | 3 | 2.99 | 167.7 | 51.4 | 183.7 |
| CB1729 | JZB | 1944.5 | -104.62095 | 38.86093 | β | 0,0,3 | 162.1 | -62.3 | 6.2 | 3 | 3.00 | 11527.7 | -75.7 | 10.7 |
| CB1730 | JZB | 1946.5 | -104.62097 | 38.86095 | α | 3,0,0 | 165.3 | -51.7 | 16.4 | 3 | 2.97 | 57.9 | -76.4 | 321.1 |
| CB1731 | JZB | 1947.8 | -104.62095 | 38.86097 | α | 1,2,0 | 178.4 | -54.0 | 18.1 | 3 | 2.96 | 47.4 | -85.5 | 272.5 |
| CB1732 | JZB | 1949.0 | -104.62094 | 38.86099 | α | 1,2,0 | 176.9 | -27.3 | 30.5 | 3 | 2.89 | 17.4 | -65.5 | 262.6 |
| CB1734 | JZB | 1955.3 | -104.62086 | 38.86106 | α | 3,0,0 | 341.9 | 58.1 | 12.9 | 3 | 2.98 | 91.9 | 75.9 | 170.5 |
| CB1740 | Hilaire-West | 2010.1 | -104.60355 | 38.87017 | α | 0,3,0 | 5.0 | 51.6 | 21.3 | 3 | 2.94 | 34.5 | 82.2 | 42.1 |
| CB1741 | Hilaire-West | 2018.3 | -104.60400 | 38.87044 | α | 3,0,0 | 343.6 | 67.8 | 49.5 | 3 | 2.73 | 7.3 | 73.4 | 216.7 |
| CB1742 | Hilaire-West | 2022.1 | -104.60439 | 38.87046 | α | 3,0,0 | 299.7 | 44.5 | 29.2 | 3 | 2.89 | 18.9 | 38.5 | 170.1 |
| CB1743 | Hilaire-West | 2036.2 | -104.60172 | 38.86974 | α | 1,2,0 | 349.3 | 53.5 | 39.0 | 3 | 2.82 | 11.1 | 80.2 | 139.3 |
| DB0902 | Leaf 6 | *1883.0* | -104.63578 | 38.84496 | α | 0,4,0 | 159.5 | 83.3 | 34.1 | 4 | 3.64 | 8.3 | 26.4 | 260.5 |
| CB1735 | Leaf 6 | 1884.1 | -104.63579 | 38.84501 | α | 3,0,0 | 13.1 | 64.6 | 13.8 | 3 | 2.98 | 81.4 | 77.7 | 302.9 |
| CB1736 | Leaf 6 | 1885.5 | -104.63620 | 38.84553 | α | 0,3,0 | 340.0 | 30.2 | 27.0 | 3 | 2.91 | 21.9 | 61.4 | 118.7 |
| DB0903 | Leaf 6 | 1886.0 | -104.63619 | 38.84552 | α/β | 2,2,0 | 5.9 | 49.1 | 18.4 | 4 | 3.88 | 26.0 | 79.9 | 44.7 |
| CB1737 | Leaf 6 | 1887.2 | -104.63700 | 38.84597 | α | 3,0,0 | 165.07 | 61.94 | 35.19 | 3 | 2.85 | 13.3 | -6.9 | 266.3 |
| CB1815 | Leaf 6 | *1888.0* | -104.63617 | 38.84564 | β | 0,0,3 | 182.7 | 3.8 | NaN | 3 | 2.84 | 3.1 | -49.2 | 251.2 |
| CB1738 | Leaf 6 | 1888.6 | -104.63769 | 38.84668 | β | 0,0,3 | 196.0 | -43.1 | NaN | 3 | 2.98 | 28.5 | -70.7 | 206.2 |
| DB0904 | Leaf 6 | 1895.5 | -104.63906 | 38.84793 | α | 2,1,0 | 153.6 | -60.6 | 24.5 | 3 | 2.92 | 26.4 | -69.7 | 1.5 |
| DB0905 | Leaf 6 | 1900.4 | -104.63962 | 38.84921 | α | 2,1,0 | 137.5 | -64.5 | 30.7 | 3 | 2.88 | 17.2 | -58.2 | 13.1 |
| DB0907 | Leaf 6 | 1911.9 | -104.64209 | 38.85116 | β | 1,0,2 | 159.9 | -31.9 | 39.0 | 3 | 2.97 | 28.8 | -62.2 | 300.1 |
| DB0908 | Leaf 6 | *1915.9* | -104.64235 | 38.85136 | β | 0,0,3 | 177.0 | -36.7 | 9.8 | 3 | 3.00 | 4539.4 | -71.4 | 264.3 |
| DB0909 | Leaf 6 | 1926.1 | -104.64304 | 38.85282 | α | 0,0,3 | 173.0 | -22.7 | 28.2 | 3 | 2.90 | 20.2 | -62.3 | 270.3 |
| DB0910 | Leaf 6 | 1930.0 | -104.64350 | 38.85318 | α | 0,0,3 | 148.1 | -36.8 | 52.2 | 3 | 2.70 | 6.7 | -57.1 | 320.8 |
| DB0913 | Leaf 6 | 1959.4 | -104.64322 | 38.85424 | β | 0,0,3 | 166.7 | -40.1 | 58.2 | 3 | 3.00 | 140.9 | -70.4 | 294.5 |
| CB1739 | Leaf 6 | 1961.8 | -104.64319 | 38.85429 | α | 2,1,0 | 126.8 | -46.0 | 19.9 | 3 | 2.95 | 39.4 | -44.6 | 347.3 |
| CB1833 | 9776 | 1945.6 | -104.64253 | 38.86695 | α | 3,0,0 | 193.8 | -58.3 | 24.6 | 3 | 2.92 | 26.2 | -79.3 | 160.2 |
| CB1830 | 9776 | 1949.6 | -104.64282 | 38.86708 | α | 3,0,0 | 176.2 | -52.5 | 23.5 | 3 | 2.93 | 28.5 | -83.5 | 284.3 |
| CB1803 | 9776 | 1950.2 | -104.64283 | 38.86712 | α | 2,1,0 | 153.4 | -41.7 | 35.0 | 3 | 2.85 | 13.5 | -63.0 | 319.8 |
| CB1831 | 9776 | 1951.1 | -104.64280 | 38.86710 | α | 1,2,0 | 296.6 | 33.5 | 56.7 | 3 | 2.66 | 5.8 | 31.9 | 163.9 |
| CB1832 | 9776 | 1952.9 | -104.64279 | 38.86712 | α | 3,0,0 | 166.9 | -59.5 | 38.5 | 3 | 2.82 | 11.3 | -79.8 | 358.0 |
| CB1802 | 9776 | 1954.3 | -104.64259 | 38.86715 | α | 3,0,0 | 356.5 | 57.2 | 5.3 | 3 | 3.00 | 538.4 | 87.1 | 145.3 |
| CB1801 | 9776 | 1956.7 | -104.64268 | 38.86720 | α | 3,0,0 | 359.2 | 51.4 | 6.8 | 3 | 2.99 | 330.9 | 83.2 | 81.0 |
| CB1745 | 9776 | 1959.0 | -104.64328 | 38.86761 | α | 3,0,0 | 353.3 | 52.6 | 9.6 | 3 | 2.99 | 164.7 | 82.2 | 121.4 |
| CB1746 | 9776 | 1984.1 | -104.64317 | 38.86859 | β | 2,0,1 | 16.8 | 57.0 | 10.0 | 3 | 2.99 | 209.6 | 76.8 | 345.6 |
| CB1873 | Lyco-Luck | *1892.3* | -104.66481 | 38.85047 | α | 3,0,0 | 355.6 | 36.1 | 14.5 | 3 | 2.97 | 73.6 | 70.8 | 87.9 |
| CB1820 | Lyco-Luck | 1900.2 | -104.66004 | 38.85321 | β | 0,0,3 | 193.2 | -22.9 | 31.8 | 3 | 3.00 | 443.4 | -60.6 | 228.2 |
| CB1807 | Lyco-Luck | *1902.0* | -104.65764 | 38.85345 | β | 0,0,3 | 201.5 | -62.5 | NaN | 3 | 2.95 | 10.0 | -73.2 | 141.3 |
| CB1748 | Lyco-Luck | *1906.0* | -104.65755 | 38.85441 | α | 1,2,0 | 177.5 | -17.9 | 29.9 | 3 | 2.89 | 18.1 | -60.2 | 260.2 |
| CB1823 | Bambino Canyon | 1931.7 | -104.58900 | 38.85736 | β | 0,0,3 | 197.5 | -63.3 | 25.0 | 3 | 3.00 | 710.8 | -75.7 | 135.3 |

| Site | Section | Elev (m) | Lon (º) | Lat (º) | α/β | PCA/Fish/GC | Dec (°) | Inc (°) | α95 (°) | n | R | k | VGP Lat (º) | VGP Lon (º) |
|---|---|---|---|---|---|---|---|---|---|---|---|---|---|---|
| CB1824 | Bambino Canyon | 1934.9 | -104.58840 | 38.85725 | α | 3,0,0 | 154.3 | -17.7 | 31.2 | 3 | 2.88 | 16.7 | -52.4 | 299.8 |
| CB1825 | Bambino Canyon | 1943.6 | -104.58768 | 38.85799 | α | 1,2,0 | 139.3 | -39.7 | 32.7 | 3 | 2.87 | 15.3 | -51.8 | 332.2 |
| CB1826 | Bambino Canyon | 1948.8 | -104.58770 | 38.85809 | α | 0,3,0 | 117.2 | -56.5 | 29.4 | 3 | 2.89 | 18.6 | -41.5 | 4.1 |
| CB1827 | Bambino Canyon | 1955.3 | -104.58768 | 38.85823 | α | 3,0,0 | 168.4 | -53.9 | 11.9 | 3 | 2.98 | 109.1 | -79.7 | 323.2 |
| CB1809 | Bambino Canyon | | -104.64893 | 38.86377 | β | 0,0,3 | 111.8 | -43.1 | NaN | 3 | 2.95 | 11.1 | -31.9 | 353.4 |
| CB1828 | Bambino Canyon | 1959.6 | -104.64663 | 38.86328 | α | 3,0,0 | 12.3 | 38.7 | 6.5 | 3 | 2.99 | 359.8 | 70.0 | 40.0 |
| CB1829 | Bambino Canyon | 1962.7 | -104.64678 | 38.86333 | α | 3,0,0 | 51.9 | 64.9 | 27.3 | 3 | 2.91 | 21.4 | 51.9 | 315.9 |
| CB1805 | Bambino Canyon | | -104.64938 | 38.86433 | α | 2,1,0 | 17.3 | 59.3 | 11.4 | 3 | 2.98 | 117.2 | 76.6 | 334.5 |
| CB1810 | Hawkins | 1901.0 | -104.58737 | 38.83270 | α | 0,3,0 | 166.2 | -23.4 | 27.2 | 3 | 2.91 | 21.6 | -60.7 | 283.9 |
| CB1811 | Hawkins | 1907.9 | -104.58495 | 38.83414 | β | 0,0,3 | 85.1 | -60.9 | NaN | 3 | 2.67 | 1.5 | -21.7 | 22.5 |
| CB1812 | Hawkins | 1937.6 | -104.57927 | 38.83548 | β | 0,0,3 | 74.7 | -47.5 | 47.9 | 3 | 3.00 | 201.9 | -6.9 | 16.9 |
| CB1814 | Hawkins | 1951.4 | -104.57520 | 38.83759 | α | 0,3,0 | 125.0 | -62.3 | 35.6 | 3 | 2.85 | 13.1 | -49.1 | 10.5 |
| CB1813 | Hawkins | 1955.1 | -104.57549 | 38.83756 | α | 0,3,0 | 184.5 | -51.7 | 54.1 | 3 | 2.68 | 6.3 | -82.6 | 224.6 |
| CB1835 | Waste Management | 1944.5 | -104.58053 | 38.84136 | α | 0,3,0 | 159.2 | -60.3 | 24.7 | 3 | 2.92 | 26.0 | -73.9 | 0.4 |
| CB1866 | Waste Management | 1946.5 | -104.58050 | 38.84138 | α | 2,2,0 | 146.9 | -55.4 | 39.9 | 4 | 3.52 | 6.3 | -63.6 | 349.5 |
| CB1865 | Waste Management | 1948.8 | -104.58043 | 38.84136 | α | 0,3,0 | 146.5 | -35.7 | 33.3 | 3 | 2.86 | 14.8 | -55.4 | 321.7 |
| CB1867 | Waste Management | 1954.6 | -104.58034 | 38.84142 | α | 3,0,0 | 12.8 | 60.4 | 7.3 | 3 | 2.99 | 288.7 | 79.9 | 326.7 |
| CB1857 | Waste Management | 1962.1 | -104.57928 | 38.84170 | α | 3,0,0 | 4.2 | 54.0 | 6.9 | 3 | 2.99 | 318.0 | 84.6 | 36.1 |
| CB1834 | Waste Management | 1965.5 | -104.57942 | 38.84185 | α | 3,0,0 | 5.4 | 56.2 | 11.4 | 3 | 2.98 | 118.2 | 85.3 | 9.5 |
| CB1856 | Waste Management | *1973.0* | -104.58229 | 38.84597 | α | 3,0,0 | 354.3 | 76.6 | 49.7 | 3 | 2.72 | 49.7 | 64.2 | 249.8 |
| CB1855 | Waste Management | *1988.0* | -104.58209 | 38.84647 | α | 2,1,0 | 346.7 | 46.2 | 27.6 | 3 | 2.90 | 21.0 | 74.2 | 123.9 |
| CB1859 | 9789 | 1872.6 | -104.62469 | 38.84311 | α | 4,0,0 | 349.1 | 63.0 | 28.8 | 4 | 3.73 | 11.1 | 80.1 | 203.1 |
| CB1860 | 9789 | 1876.8 | -104.62784 | 38.84634 | α | 0,4,0 | 353.5 | 63.0 | 26.8 | 4 | 3.76 | 12.8 | 82.6 | 216.9 |
| CB1861 | 9789 | 1883.8 | -104.62723 | 38.84787 | β | 1,0,3 | 185.3 | -40.9 | 61.0 | 4 | 3.89 | 9.5 | -74.0 | 237.7 |
| CB1863 | 9789 | 1902.3 | -104.63521 | 38.85091 | α | 3,0,1 | 155.1 | -54.9 | 21.5 | 4 | 3.84 | 19.3 | -69.9 | 343.2 |
| CB1838 | 9789 | 1925.6 | -104.63670 | 38.85554 | α | 0,3,0 | 325.8 | -68.4 | 41.1 | 3 | 2.80 | 10.1 | -5.3 | 95.9 |
| CB1839 | 9789 | 1935.5 | -104.63637 | 38.85745 | β | 1,0,2 | 282.5 | -48.9 | 41.8 | 3 | 3.00 | 261.9 | -9.6 | 134.6 |
| CB1841 | 9789 | 1946.7 | -104.63589 | 38.85813 | β | 0,0,3 | 191.6 | -57.1 | 57.4 | 3 | 3.00 | 144.5 | -80.8 | 169.3 |
| CB1843 | 9789 | 1951.8 | -104.63573 | 38.85839 | α | 0,3,0 | 194.4 | -64.2 | 31.1 | 3 | 2.88 | 16.8 | -77.2 | 126.7 |
| CB1844 | 9789 | 1954.9 | -104.63583 | 38.85841 | β | 0,0,3 | 176.3 | -64.6 | 45.4 | 3 | 3.00 | 223.4 | -81.9 | 56.7 |
| CB1845 | 9789 | 1958.9 | -104.63571 | 38.85855 | α | 3,0,0 | 165.5 | -47.3 | 17.1 | 3 | 2.96 | 53.1 | -74.1 | 308.9 |
| CB1846 | 9789 | 1962.6 | -104.63584 | 38.85866 | α | 3,0,0 | 0.8 | 60.5 | 7.2 | 3 | 2.99 | 298.1 | 87.4 | 268.9 |
| CB1847 | 9789 | 1965.8 | -104.63585 | 38.85870 | α | 3,0,0 | 349.6 | 52.5 | 6.4 | 3 | 2.99 | 375.1 | 79.8 | 133.8 |
| CB1848 | 9789 | 1971.2 | -104.63490 | 38.85886 | α | 3,0,0 | 15.3 | 58.4 | 15.9 | 3 | 2.97 | 60.9 | 78.1 | 339.5 |
| CB1850 | Kunstle | 2029.0 | -104.59666 | 38.86847 | α | 3,0,0 | 332.6 | 57.5 | 23.6 | 3 | 2.93 | 28.3 | 68.6 | 172.0 |
| CB1851 | Kunstle | 2036.0 | -104.59621 | 38.86927 | α | 3,0,0 | 358.8 | 59.5 | 19.2 | 3 | 2.95 | 42.2 | 88.3 | 223.7 |
| CB1868 | Kunstle | 2036.9 | -104.59613 | 38.86965 | α | 3,0,0 | 340.7 | 55.1 | 33.2 | 3 | 2.87 | 14.9 | 74.3 | 159.4 |
| CB1869 | Kunstle | 2040.6 | -104.59588 | 38.86971 | α | 1,2,0 | 155.1 | -37.0 | 27.3 | 3 | 2.91 | 21.4 | -61.9 | 312.2 |
| CB1853 | Kunstle | 2042.4 | -104.59593 | 38.86977 | α | 4,0,0 | 176.1 | -31.4 | 28.5 | 4 | 3.74 | 11.3 | -67.8 | 265.3 |
| CB1870 | Kunstle | 2043.4 | -104.59591 | 38.86977 | α | 0,4,0 | 166.3 | -48.7 | 23.8 | 4 | 3.81 | 15.8 | -75.4 | 310.3 |
| CB1854 | Kunstle | 2043.7 | -104.59591 | 38.86980 | α | 3,0,0 | 348.2 | 56.4 | 8.2 | 3 | 2.99 | 230.0 | 80.5 | 157.5 |
| CB1871 | Kunstle | 2045.1 | -104.59585 | 38.86979 | β | 0,0,4 | 136.5 | -49.0 | 12.4 | 4 | 4.00 | 202.5 | -53.4 | 344.7 |

**Table S1**: Site calculations for sites that passed the Watson test (Watson, 1953). Sites are listed in superposition by section. **Elev–** refers to site DGPS elevation in meters, italicized values represent estimated elevations (see methods) , blank values are for sites where we could not determine elevation; **Lon/Lat–** DGPS Longitude and Latitude of site relative to WGS 84 Datum; **α/β–** Site reliability level (alpha/beta site); **PCA/Fish/GC–** Indicates number of PCAs, Fisher Means, and/or Great Circles (respectively) used to characterize a site; **Dec/Inc-** Declination and inclination of site mean vector; **α95 –** 95 percent confidence interval of site mean; n– number of samples used in site statistics; **R–** Length of the resultant vector; **k–** precision parameter; **VGP Lat/Lon–** site VGP latitude and longitude.

| Sample | Section | Elev (m) | Lon(º) | Lat(º) | NRM Intensity (mA/m) | Method | Dec (°) | Inc (°) | MAD/α95 (°) | n points | Start (mT) | End (mT) |
|---|---|---|---|---|---|---|---|---|---|---|---|---|
| CB1836D | Bishop Wash | 1901.8 | -104.59455 | 38.84271 | 2.18 | Fisher | 57.7 | -24.9 | 4.99 | 18 | 3 | 100 |
| CB1836E | Bishop Wash | 1901.8 | -104.59455 | 38.84271 | 2.40 | Fisher | 138.4 | -56.4 | 5.22 | 6 | 20 | 80 |
| CB1836F | Bishop Wash | 1901.8 | -104.59455 | 38.84271 | 2.35 | Fisher | 138.5 | -35.3 | 4.21 | 18 | 3 | 100 |
| CB1701A | Bishop Wash | 1904.3 | -104.59561 | 38.84592 | 3.72 | GC | 208.0 | -44.5 | 15.00 | 8 | 6 | 27 |
| CB1701B | Bishop Wash | 1904.3 | -104.59561 | 38.84592 | 2.81 | GC | 205.7 | -24.8 | 7.50 | 4 | 12 | 24 |
| CB1701C | Bishop Wash | 1904.3 | -104.59561 | 38.84592 | 2.78 | GC | 217.2 | -38.3 | 38.40 | 8 | 0 | 21 |
| CB1701D | Bishop Wash | 1904.3 | -104.59561 | 38.84592 | 7.70 | GC | 198.5 | -28.6 | 11.20 | 7 | 9 | 30 |
| CB1702B | Bishop Wash | 1908.2 | -104.59324 | 38.84781 | 7.89 | Fisher | 211.6 | -66.3 | 7.09 | 10 | 27 | 70 |
| CB1702C | Bishop Wash | 1908.2 | -104.59324 | 38.84781 | 2.12 | Fisher | 238.7 | -56.5 | 3.52 | 13 | 9 | 55 |
| CB1702D | Bishop Wash | 1908.2 | -104.59324 | 38.84781 | 18.50 | Fisher | 129.2 | -55.8 | 15.68 | 6 | 40 | 70 |
| CB1703A | Bishop Wash | 1912.1 | -104.59224 | 38.84773 | 23.40 | PCA | 166.0 | -21.6 | 14.10 | 14 | 21 | 100 |
| CB1703B | Bishop Wash | 1912.1 | -104.59224 | 38.84773 | 22.40 | PCA | 154.5 | -50.9 | 13.20 | 6 | 250 C° | 500 C° |
| CB1703C | Bishop Wash | 1912.1 | -104.59224 | 38.84773 | 8.38 | GC | 149.2 | -36.0 | 11.70 | 16 | 0 | 55 |
| CB1704A | Bishop Wash | 1919.0 | -104.59244 | 38.84765 | 64.90 | Fisher | 134.5 | -10.7 | 8.48 | 4 | 50 | 70 |
| CB1704B | Bishop Wash | 1919.0 | -104.59244 | 38.84765 | 12.37 | Fisher | 123.6 | -11.9 | 4.49 | 10 | 21 | 60 |
| CB1704C | Bishop Wash | 1919.0 | -104.59244 | 38.84765 | 4.88 | PCA | 161.3 | -17.3 | 11.80 | 18 | 6 | 90 |
| CB1705A | Bishop Wash | 1924.2 | -104.59190 | 38.85159 | 1.41 | GC | 96.0 | -48.7 | 15.40 | 10 | 6 | 35 |
| CB1705B | Bishop Wash | 1924.2 | -104.59190 | 38.85159 | 1.31 | GC | 118.1 | -35.9 | 15.00 | 8 | 3 | 24 |
| CB1705D | Bishop Wash | 1924.2 | -104.59190 | 38.85159 | 2.05 | GC | 103.0 | -50.1 | 11.80 | 9 | 3 | 27 |
| CB1706B | Bishop Wash | 1933.0 | -104.59212 | 38.85552 | 6.38 | GC | 160.0 | -15.4 | 7.00 | 6 | 0 | 15 |
| CB1706C | Bishop Wash | 1933.0 | -104.59212 | 38.85552 | 6.41 | GC | 135.2 | -33.7 | 11.40 | 6 | 3 | 18 |
| CB1706D | Bishop Wash | 1933.0 | -104.59212 | 38.85552 | 2.28 | GC | 150.1 | -4.5 | 3.30 | 3 | 21 | 27 |
| CB1707A | Bishop Wash | 1935.8 | -104.59144 | 38.85562 | 58.69 | Fisher | 156.0 | -46.0 | 27.91 | 3 | 80 | 100 |
| CB1707B | Bishop Wash | 1935.8 | -104.59144 | 38.85562 | 49.70 | Fisher | 166.4 | -31.8 | 5.89 | 7 | 45 | 90 |
| CB1707C | Bishop Wash | 1935.8 | -104.59144 | 38.85562 | 35.61 | Fisher | 169.6 | -34.6 | 4.83 | 9 | 40 | 100 |
| CB1708A | Bishop Wash | 1936.9 | -104.59077 | 38.85591 | 14.90 | GC | 176.8 | -53.5 | 14.90 | 10 | 0 | 27 |
| CB1708B | Bishop Wash | 1936.9 | -104.59077 | 38.85591 | 28.78 | GC | 177.2 | -53.1 | 14.10 | 15 | 0 | 50 |
| CB1708D | Bishop Wash | 1936.9 | -104.59077 | 38.85591 | 84.10 | GC | 176.8 | -53.9 | 15.80 | 14 | 0 | 45 |
| CB1709B | Bishop Wash | 1940.6 | -104.59073 | 38.85586 | 10.90 | Fisher | 144.8 | -12.6 | 4.20 | 15 | 6 | 60 |
| CB1709C | Bishop Wash | 1940.6 | -104.59073 | 38.85586 | 10.16 | Fisher | 201.0 | -29.5 | 19.60 | 4 | 40 | 55 |
| CB1709D | Bishop Wash | 1940.6 | -104.59073 | 38.85586 | 3.64 | Fisher | 165.7 | -61.6 | 8.26 | 9 | 24 | 60 |
| CB1875A | Bishop Wash | *1945.2* | -104.59073 | 38.85582 | 15.28 | PCA | 338.0 | 54.3 | 7.80 | 7 | 12 | 40 |
| CB1875B | Bishop Wash | *1945.2* | -104.59073 | 38.85582 | 5.31 | PCA | 351.7 | 47.9 | 9.80 | 8 | 3 | 30 |
| CB1875C | Bishop Wash | *1945.2* | -104.59073 | 38.85582 | 1.21 | Fisher | 3.0 | 27.0 | 3.04 | 6 | 0 | 15 |
| CB1710A | Bishop Wash | 1946.1 | -104.59071 | 38.85582 | 12.00 | PCA | 331.3 | 56.5 | 13.70 | 10 | 24 | 70 |
| CB1710C | Bishop Wash | 1946.1 | -104.59071 | 38.85582 | 4.24 | PCA | 343.1 | 57.7 | 12.80 | 16 | 6 | 70 |
| CB1710D | Bishop Wash | 1946.1 | -104.59071 | 38.85582 | 8.41 | PCA | 355.3 | 56.9 | 7.40 | 17 | 9 | 90 |
| CB1711A | Bishop Wash | 1948.1 | -104.59068 | 38.85579 | 17.40 | PCA | 16.7 | 58.2 | 8.00 | 17 | 12 | 100 |
| CB1711B | Bishop Wash | 1948.1 | -104.59068 | 38.85579 | 17.60 | PCA | 343.4 | 63.7 | 10.40 | 17 | 12 | 100 |
| CB1711C | Bishop Wash | 1948.1 | -104.59068 | 38.85579 | 15.13 | PCA | 1.4 | 62.1 | 9.50 | 16 | 15 | 100 |
| CB1712A | Bishop Wash | 1948.3 | -104.58986 | 38.85611 | 64.88 | PCA | 328.4 | 54.0 | 8.00 | 13 | 24 | 100 |
| CB1712B | Bishop Wash | 1948.3 | -104.58986 | 38.85611 | 208.00 | PCA | 1.0 | 62.3 | 4.40 | 13 | 9 | 55 |
| CB1712D | Bishop Wash | 1948.3 | -104.58986 | 38.85611 | 128.00 | PCA | 6.4 | 55.2 | 5.40 | 6 | 18 | 35 |
| CB1713A | Bishop Wash | 1951.3 | -104.58982 | 38.85627 | 11.10 | PCA | 325.2 | 68.7 | 7.70 | 20 | 3 | 100 |
| CB1713B | Bishop Wash | 1951.3 | -104.58982 | 38.85627 | 7.89 | PCA | 359.5 | 69.0 | 15.40 | 18 | 9 | 100 |
| CB1713C | Bishop Wash | 1951.3 | -104.58982 | 38.85627 | 10.31 | PCA | 347.7 | 74.4 | 10.80 | 16 | 15 | 100 |
| CB1714A | Bishop Wash | 1953.5 | -104.58988 | 38.85628 | 11.70 | PCA | 2.5 | 52.1 | 11.50 | 17 | 12 | 100 |
| CB1714C | Bishop Wash | 1953.5 | -104.58988 | 38.85628 | 13.60 | PCA | 13.3 | 63.5 | 10.60 | 18 | 6 | 90 |
| CB1714D | Bishop Wash | 1953.5 | -104.58988 | 38.85628 | 15.49 | PCA | 11.5 | 44.3 | 10.20 | 17 | 12 | 100 |
| CB1715A | Bishop Wash | 1957.3 | -104.58999 | 38.85646 | 83.50 | PCA | 6.0 | 65.0 | 2.60 | 19 | 6 | 100 |
| CB1715B | Bishop Wash | 1957.3 | -104.58999 | 38.85646 | 70.80 | PCA | 347.4 | 65.0 | 2.80 | 19 | 9 | 100 |
| CB1715D | Bishop Wash | 1957.3 | -104.58999 | 38.85646 | 46.94 | PCA | 352.3 | 58.8 | 2.90 | 17 | 12 | 100 |
| CB1716A | Bishop Wash | 1962.1 | -104.58988 | 38.85653 | 46.10 | PCA | 5.3 | 59.8 | 3.90 | 12 | 9 | 50 |
| CB1716C | Bishop Wash | 1962.1 | -104.58988 | 38.85653 | 40.80 | PCA | 3.5 | 56.5 | 5.50 | 16 | 15 | 100 |
| CB1716D | Bishop Wash | 1962.1 | -104.58988 | 38.85653 | 22.49 | PCA | 2.8 | 58.7 | 6.00 | 18 | 9 | 100 |
| CB1717A | Bishop Wash | 1966.1 | -104.58981 | 38.85663 | 82.07 | PCA | 344.4 | 60.1 | 3.40 | 17 | 3 | 70 |
| CB1717B | Bishop Wash | 1966.1 | -104.58981 | 38.85663 | 34.25 | PCA | 4.4 | 70.0 | 15.90 | 16 | 9 | 80 |
| CB1717C | Bishop Wash | 1966.1 | -104.58981 | 38.85663 | 17.70 | PCA | 352.6 | 57.9 | 11.70 | 16 | 6 | 70 |
| CB1718A | Bishop Wash | 1969.7 | -104.58977 | 38.85684 | 60.00 | PCA | 4.3 | 61.6 | 5.60 | 19 | 6 | 100 |
| CB1718C | Bishop Wash | 1969.7 | -104.58977 | 38.85684 | 14.60 | PCA | 338.6 | 58.4 | 13.50 | 14 | 15 | 80 |
| CB1718E | Bishop Wash | 1969.7 | -104.58977 | 38.85684 | 23.40 | PCA | 316.5 | 44.7 | 15.70 | 17 | 12 | 100 |
| CB1719A | Bishop Wash | 1973.1 | -104.58972 | 38.85697 | 34.10 | PCA | 343.8 | 51.7 | 3.70 | 17 | 12 | 100 |
| CB1719C | Bishop Wash | 1973.1 | -104.58972 | 38.85697 | 13.39 | PCA | 338.7 | 61.2 | 6.20 | 14 | 21 | 100 |
| CB1719D | Bishop Wash | 1973.1 | -104.58972 | 38.85697 | 40.52 | PCA | 338.0 | 49.7 | 2.50 | 17 | 12 | 100 |
| CB1720A | Bishop Wash | 1976.2 | -104.58970 | 38.85702 | 91.00 | PCA | 333.6 | 62.1 | 7.30 | 15 | 18 | 100 |
| CB1720C | Bishop Wash | 1976.2 | -104.58970 | 38.85702 | 55.16 | PCA | 350.1 | 65.7 | 2.90 | 18 | 9 | 100 |
| CB1720D | Bishop Wash | 1976.2 | -104.58970 | 38.85702 | 34.45 | PCA | 346.9 | 67.7 | 3.50 | 17 | 12 | 100 |
| CB1721A | Bishop Wash | 1979.4 | -104.58959 | 38.85709 | 48.70 | PCA | 7.5 | 61.4 | 11.40 | 18 | 9 | 100 |
| CB1721B | Bishop Wash | 1979.4 | -104.58959 | 38.85709 | 26.10 | PCA | 344.8 | 53.8 | 8.10 | 17 | 12 | 100 |
| CB1721C | Bishop Wash | 1979.4 | -104.58959 | 38.85709 | 57.75 | PCA | 351.5 | 58.6 | 3.20 | 16 | 15 | 100 |
| CB1722B | Bishop Wash | 1981.1 | -104.58915 | 38.85728 | 45.60 | PCA | 76.2 | -11.1 | 7.60 | 15 | 18 | 100 |
| CB1722C | Bishop Wash | 1981.1 | -104.58915 | 38.85728 | 44.60 | PCA | 333.5 | 66.4 | 5.90 | 17 | 12 | 100 |
| CB1722D | Bishop Wash | 1981.1 | -104.58915 | 38.85728 | 42.35 | PCA | 342.3 | 58.7 | 5.60 | 17 | 12 | 100 |
| CB1723B | Bishop Wash | 1984.5 | -104.58900 | 38.85736 | 7.80 | Fisher | 332.1 | 62.4 | 6.71 | 11 | 3 | 35 |
| CB1723C | Bishop Wash | 1984.5 | -104.58900 | 38.85736 | 3.79 | Fisher | 231.6 | 34.9 | 8.39 | 12 | 15 | 60 |
| CB1723D | Bishop Wash | 1984.5 | -104.58900 | 38.85736 | 4.75 | Fisher | 309.3 | 56.0 | 5.51 | 13 | 3 | 45 |
| CB1724B | Bishop Wash | 1991.4 | -104.58840 | 38.85725 | 181.50 | PCA | 355.2 | 43.9 | 2.10 | 16 | 6 | 70 |

| Sample | Section | Elev (m) | Lon(º) | Lat(º) | NRM Intensity (mA/m) | Method | Dec (°) | Inc (°) | MAD/α95 (°) | n points | Start (mT) | End (mT) |
|---|---|---|---|---|---|---|---|---|---|---|---|---|
| CB1724C | Bishop Wash | 1991.4 | -104.58840 | 38.85725 | 276.00 | PCA | 71.5 | 65.8 | 3.50 | 19 | 6 | 100 |
| CB1724D | Bishop Wash | 1991.4 | -104.58840 | 38.85725 | 149.10 | PCA | 2.8 | 41.8 | 5.50 | 16 | 15 | 100 |
| CB1725A | Bishop Wash | 1995.9 | -104.58768 | 38.85799 | 334.00 | PCA | 336.6 | 47.4 | 2.30 | 18 | 9 | 100 |
| CB1725B | Bishop Wash | 1995.9 | -104.58768 | 38.85799 | 235.70 | PCA | 334.7 | 59.8 | 2.50 | 18 | 9 | 100 |
| CB1725C | Bishop Wash | 1995.9 | -104.58768 | 38.85799 | 722.10 | PCA | 349.5 | 45.5 | 3.50 | 18 | 9 | 100 |
| CB1726A | Bishop Wash | 2001.3 | -104.58770 | 38.85809 | 91.70 | PCA | 0.2 | 52.5 | 3.50 | 19 | 6 | 100 |
| CB1726C | Bishop Wash | 2001.3 | -104.58770 | 38.85809 | 67.12 | PCA | 343.0 | 59.5 | 6.10 | 17 | 12 | 100 |
| CB1726E | Bishop Wash | 2001.3 | -104.58770 | 38.85809 | 78.25 | PCA | 338.7 | 57.2 | 5.90 | 15 | 18 | 100 |
| CB1727A | Bishop Wash | 2005.5 | -104.58768 | 38.85823 | 563.30 | PCA | 314.9 | 58.4 | 14.90 | 10 | 24 | 70 |
| CB1727D | Bishop Wash | 2005.5 | -104.58768 | 38.85823 | 992.10 | Fisher | 317.8 | 59.6 | 7.68 | 11 | 18 | 60 |
| CB1727E | Bishop Wash | 2005.5 | -104.58768 | 38.85823 | 974.00 | Fisher | 295.6 | 58.5 | 5.13 | 12 | 15 | 60 |
| CB1728A | JZB | 1941.2 | -104.62101 | 38.86086 | 9.32 | Fisher | 229.7 | 13.9 | 3.03 | 18 | 3 | 100 |
| CB1728C | JZB | 1941.2 | -104.62101 | 38.86086 | 6.18 | Fisher | 218.2 | -12.4 | 2.84 | 10 | 6 | 35 |
| CB1728D | JZB | 1941.2 | -104.62101 | 38.86086 | 8.81 | Fisher | 166.9 | -24.4 | 5.82 | 8 | 6 | 27 |
| CB1729B | JZB | 1944.5 | -104.62095 | 38.86093 | 2.64 | GC | 161.4 | -62.0 | 14.90 | 17 | 0 | 100 |
| CB1729C | JZB | 1944.5 | -104.62095 | 38.86093 | 8.56 | GC | 162.1 | -62.6 | 7.70 | 13 | 0 | 40 |
| CB1729E | JZB | 1944.5 | -104.62095 | 38.86093 | 10.71 | GC | 162.8 | -62.4 | 12.90 | 19 | 0 | 80 |
| CB1730A | JZB | 1946.5 | -104.62097 | 38.86095 | 1.85 | PCA | 158.8 | -48.0 | 14.20 | 11 | 24 | 80 |
| CB1730B | JZB | 1946.5 | -104.62097 | 38.86095 | 13.30 | PCA | 162.6 | -44.8 | 7.70 | 16 | 15 | 100 |
| CB1730D | JZB | 1946.5 | -104.62097 | 38.86095 | 26.98 | PCA | 178.3 | -61.4 | 2.10 | 12 | 27 | 100 |
| CB1731B | JZB | 1947.8 | -104.62095 | 38.86097 | 3.05 | Fisher | 189.5 | -58.9 | 6.57 | 19 | 6 | 100 |
| CB1731C | JZB | 1947.8 | -104.62095 | 38.86097 | 1.27 | Fisher | 173.1 | -60.9 | 5.63 | 17 | 12 | 100 |
| CB1731D | JZB | 1947.8 | -104.62095 | 38.86097 | 5.03 | PCA | 174.2 | -41.6 | 12.00 | 15 | 18 | 100 |
| CB1732A | JZB | 1949.0 | -104.62094 | 38.86099 | 3.12 | Fisher | 176.9 | -46.9 | 18.36 | 6 | 21 | 40 |
| CB1732D | JZB | 1949.0 | -104.62094 | 38.86099 | 1.66 | PCA | 168.6 | -23.2 | 14.00 | 17 | 12 | 100 |
| CB1732E | JZB | 1949.0 | -104.62094 | 38.86099 | 2.01 | Fisher | 184.7 | -11.5 | 9.00 |  | 27 | 70 |
| CB1733A | JZB | 1950.6 | -104.62093 | 38.86101 | 5.15 | GC | 180.4 | -48.8 | 15.60 | 14 | 0 | 45 |
| CB1733B | JZB | 1950.6 | -104.62093 | 38.86101 | 6.50 | GC | 187.8 | -48.4 | 19.10 | 8 | 0 | 21 |
| CB1733C | JZB | 1950.6 | -104.62093 | 38.86101 | 6.93 | PCA | 30.2 | 53.8 | 15.60 | 19 | 3 | 90 |
| CB1734A | JZB | 1955.3 | -104.62086 | 38.86106 | 58.93 | PCA | 347.7 | 60.1 | 3.10 | 15 | 3 | 55 |
| CB1734B | JZB | 1955.3 | -104.62086 | 38.86106 | 69.70 | PCA | 353.4 | 56.2 | 3.80 | 17 | 12 | 100 |
| CB1734D | JZB | 1955.3 | -104.62086 | 38.86106 | 75.05 | PCA | 325.1 | 56.0 | 2.50 | 17 | 12 | 100 |
| CB1740C | Hillaire-West | 2010.1 | -104.60355 | 38.87017 | 7.46 | Fisher | 348.7 | 43.7 | 4.43 | 16 | 6 | 70 |
| CB1740D | Hillaire-West | 2010.1 | -104.60355 | 38.87017 | 4.32 | Fisher | 2.8 | 57.1 | 5.32 | 15 | 0 | 50 |
| CB1740F | Hillaire-West | 2010.1 | -104.60355 | 38.87017 | 6.63 | Fisher | 25.9 | 50.9 | 5.01 | 18 | 3 | 40 |
| CB1741A | Hillaire-West | 2018.3 | -104.60400 | 38.87044 | 151.00 | PCA | 335.4 | 51.1 | 3.20 | 13 | 6 | 50 |
| CB1741B | Hillaire-West | 2018.3 | -104.60400 | 38.87044 | 147.00 | PCA | 258.0 | 69.1 | 5.40 | 20 | 0 | 90 |
| CB1741C | Hillaire-West | 2018.3 | -104.60400 | 38.87044 | 73.02 | PCA | 33.1 | 54.3 | 3.70 | 9 | 3 | 27 |
| CB1742D | Hillaire-West | 2022.1 | -104.60439 | 38.87046 | 4.17 | PCA | 294.4 | 52.8 | 7.40 | 18 | 9 | 100 |
| CB1742E | Hillaire-West | 2022.1 | -104.60439 | 38.87046 | 2.35 | PCA | 298.6 | 57.0 | 5.50 | 17 | 9 | 90 |
| CB1742F | Hillaire-West | 2022.1 | -104.60439 | 38.87046 | 8.50 | PCA | 303.8 | 23.2 | 5.70 | 17 | 12 | 100 |
| CB1743A | Hillaire-West | 2036.2 | -104.60172 | 38.86974 | 3.55 | Fisher | 319.8 | 58.0 | 3.49 | 20 | 3 | 100 |
| CB1743D | Hillaire-West | 2036.2 | -104.60172 | 38.86974 | 4.86 | Fisher | 346.5 | 66.6 | 3.45 | 18 | 6 | 90 |
| CB1743E | Hillaire-West | 2036.2 | -104.60172 | 38.86974 | 3.53 | PCA | 8.2 | 30.0 | 10.50 | 16 | 3 | 60 |
| DB0902B | Leaf 6 | *1883* | -104.63578 | 38.84496 | 1.94 | Fisher | 130.3 | 74.8 | 4.64 | 11 | 5 | 30 |
| DB0902C | Leaf 6 | *1883* | -104.63578 | 38.84496 | 2.54 | Fisher | 3.4 | 61.4 | 5.17 | 6 | 3 | 15 |
| DB0902D | Leaf 6 | *1883* | -104.63578 | 38.84496 | 42.70 | Fisher | 139.9 | 66.5 | 3.60 | 11 | 3 | 28 |
| DB0902E | Leaf 6 | *1883* | -104.63578 | 38.84496 | 2.42 | Fisher | 220.1 | 58.4 | 5.60 | 11 | 13 | 45 |
| CB1735A | Leaf 6 | 1884.1 | -104.63579 | 38.84501 | 27.73 | PCA | 7.3 | 67.6 | 7.20 | 14 | 6 | 55 |
| CB1735B | Leaf 6 | 1884.1 | -104.63579 | 38.84501 | 61.06 | PCA | 31.2 | 68.4 | 6.20 | 9 | 3 | 27 |
| CB1735D | Leaf 6 | 1884.1 | -104.63579 | 38.84501 | 89.40 | PCA | 5.3 | 56.4 | 5.30 | 10 | 6 | 35 |
| CB1736B | Leaf 6 | 1885.5 | -104.63620 | 38.84553 | 4.95 | Fisher | 336.2 | 26.9 | 6.62 | 15 | 9 | 70 |
| CB1736C | Leaf 6 | 1885.5 | -104.63620 | 38.84553 | 3.88 | Fisher | 329.2 | 19.5 | 7.34 | 16 | 3 | 60 |
| CB1736D | Leaf 6 | 1885.5 | -104.63620 | 38.84553 | 6.79 | Fisher | 358.5 | 42.6 | 3.94 | 17 | 3 | 70 |
| DB0903A | Leaf 6 | 1886.0 | -104.63619 | 38.84552 | 3.28 | PCA | 11.3 | 59.9 | 13.40 | 10 | 2.5 | 22.5 |
| DB0903B | Leaf 6 | 1886.0 | -104.63619 | 38.84552 | 4.05 | PCA | 11.9 | 64.6 | 12.40 | 11 | 2.5 | 27.5 |
| DB0903C | Leaf 6 | 1886.0 | -104.63619 | 38.84552 | 2.41 | Fisher | 0.6 | 38.5 | 12.59 | 4 | 0 | 8 |
| DB0903D | Leaf 6 | 1886.0 | -104.63619 | 38.84552 | 1.96 | Fisher | 4.6 | 33.0 | 4.20 | 12 | 0 | 28 |
| CB1737A | Leaf 6 | 1887.2 | -104.63700 | 38.84597 | 5.01 | PCA | 177.6 | 46.4 | 10.4 | 16 | 0 | 55 |
| CB1737C | Leaf 6 | 1887.2 | -104.63700 | 38.84597 | 14.20 | PCA | 201.3 | 70.2 | 6.8 | 14 | 0 | 45 |
| CB1737D | Leaf 6 | 1887.2 | -104.63700 | 38.84597 | 6.23 | PCA | 123.5 | 58.2 | 8.3 | 8 | 3 | 24 |
| CB1815B | Leaf 6 | *1888.0* | -104.63617 | 38.84564 | 3.68 | GC | 203.4 | 15.4 | 9.10 | 10 | 0 | 27 |
| CB1815C | Leaf 6 | *1888.0* | -104.63617 | 38.84564 | 20.06 | GC | 183.3 | 1.3 | 4.10 | 4 | 24 | 35 |
| CB1815E | Leaf 6 | *1888.0* | -104.63617 | 38.84564 | 4.89 | GC | 162.0 | -5.8 | 10.10 | 6 | 3 | 18 |
| CB1738D | Leaf 6 | 1888.6 | -104.63769 | 38.84668 | 5.40 | GC | 186.0 | -41.4 | 17.80 | 12 | 0 | 35 |
| CB1738E | Leaf 6 | 1888.6 | -104.63769 | 38.84668 | 4.25 | GC | 206.4 | -44.2 | 14.00 | 9 | 0 | 24 |
| CB1738F | Leaf 6 | 1888.6 | -104.63769 | 38.84668 | 3.51 | GC | 196.0 | -42.8 | 13.40 | 6 | 9 | 24 |
| DB0904A | Leaf 6 | 1895.5 | -104.63906 | 38.84793 | 2.25 | PCA | 166.7 | -61.2 | 12.30 | 19 | 5 | 65 |
| DB0904B | Leaf 6 | 1895.5 | -104.63906 | 38.84793 | 1.60 | Fisher | 119.5 | -56.5 | 4.04 | 12 | 3 | 60 |
| DB0904D | Leaf 6 | 1895.5 | -104.63906 | 38.84793 | 2.17 | PCA | 174.9 | -56.8 | 12.30 | 10 | 15 | 100 |
| DB0905B | Leaf 6 | 1900.4 | -104.63962 | 38.84921 | 1.17 | PCA | 196.3 | -70.8 | 14.20 | 9 | 20 | 100 |
| DB0905C | Leaf 6 | 1900.4 | -104.63962 | 38.84921 | 3.85 | PCA | 125.6 | -60.3 | 6.00 | 8 | 15 | 60 |
| DB0905D | Leaf 6 | 1900.4 | -104.63962 | 38.84921 | 1.63 | Fisher | 120.5 | -52.2 | 9.32 | 7 | 50 | 80 |
| DB0906A | Leaf 6 | 1903.05 | -104.64076 | 38.84945 | 0.62 | Fisher | 180.6 | -23.5 | 6.86 | 10 | 9 | 60 |
| DB0906C | Leaf 6 | 1903.05 | -104.64076 | 38.84945 | 0.36 | Fisher | 150.5 | -0.3 | 21.40 | 5 | 40 | 100 |
| DB0906D | Leaf 6 | 1903.05 | -104.64076 | 38.84945 | 0.37 | Fisher | 217.7 | -49.3 | 10.08 | 19 | 5 | 70 |
| DB0907A | Leaf 6 | 1911.9 | -104.64209 | 38.85116 | 1.74 | GC | 238.1 | 157.5 | -32.5 | 8 | 6 | 35 |

| Sample | Section | Elev (m) | Lon(º) | Lat(º) | NRM Intensity (mA/m) | Method | Dec (°) | Inc (°) | MAD/α95 (°) | n points | Start (mT) | End (mT) |
|---|---|---|---|---|---|---|---|---|---|---|---|---|
| DB0907B | Leaf 6 | 1911.9 | -104.64209 | 38.85116 | 2.82 | PCA | 161.3 | -21.0 | 9.30 | 11 | 12 | 100 |
| DB0907D | Leaf 6 | 1911.9 | -104.64209 | 38.85116 | 0.90 | GC | 160.8 | -42.1 | 12.80 | 8 | 0 | 18 |
| DB0908A | Leaf 6 | *1915.9* | -104.64235 | 38.85136 | 68.40 | GC | 177.0 | -36.7 | 6.50 | 9 | 18 | 45 |
| DB0908B | Leaf 6 | *1915.9* | -104.64235 | 38.85136 | 49.49 | GC | 177.2 | -36.2 | 3.70 | 6 | 9 | 30 |
| DB0908D | Leaf 6 | *1915.9* | -104.64235 | 38.85136 | 32.43 | GC | 176.8 | -37.4 | 5.30 | 6 | 9 | 24 |
| DB0909A | Leaf 6 | 1926.1 | -104.64304 | 38.85282 | 1.43 | Fisher | 159.8 | -9.6 | 5.88 | 13 | 6 | 100 |
| DB0909B | Leaf 6 | 1926.1 | -104.64304 | 38.85282 | 1.46 | Fisher | 189.7 | -25.4 | 5.02 | 12 | 6 | 80 |
| DB0909C | Leaf 6 | 1926.1 | -104.64304 | 38.85282 | 1.69 | Fisher | 170.5 | -31.8 | 6.33 | 11 | 10 | 40 |
| DB0910A | Leaf 6 | 1930.0 | -104.64350 | 38.85318 | 1.96 | Fisher | 134.7 | -14.8 | 15.96 | 5 | 15 | 35 |
| DB0910B | Leaf 6 | 1930.0 | -104.64350 | 38.85318 | 2.07 | Fisher | 188.0 | -39.1 | 5.34 | 8 | 3 | 20 |
| DB0910C | Leaf 6 | 1930.0 | -104.64350 | 38.85318 | 2.66 | Fisher | 124.5 | -47.0 | 6.11 | 8 | 3 | 20 |
| DB0913A | Leaf 6 | 1959.4 | -104.64322 | 38.85424 | 34.30 | GC | 164.3 | -42.1 | 10.30 | 5 | 15 | 35 |
| DB0913B | Leaf 6 | 1959.4 | -104.64322 | 38.85424 | 34.02 | GC | 171.2 | -38.4 | 4.20 | 8 | 3 | 30 |
| DB0913D | Leaf 6 | 1959.4 | -104.64322 | 38.85424 | 41.27 | GC | 165.4 | -41.0 | 5.60 | 9 | 13 | 35 |
| CB1739A | Leaf 6 | 1961.8 | -104.64319 | 38.85429 | 4.17 | Fisher | 144.9 | -50.8 | 2.95 | 16 | 9 | 80 |
| CB1739B | Leaf 6 | 1961.8 | -104.64319 | 38.85429 | 6.53 | PCA | 115.7 | -48.8 | 10.90 | 18 | 9 | 100 |
| CB1739C | Leaf 6 | 1961.8 | -104.64319 | 38.85429 | 18.80 | PCA | 121.7 | -36.5 | 12.20 | 15 | 12 | 80 |
| CB1833B | 9776 | 1945.6 | -104.64253 | 38.86695 | 5.38 | PCA | 173.8 | -64.6 | 11.40 | 12 | 21 | 100 |
| CB1833C | 9776 | 1945.6 | -104.64253 | 38.86695 | 11.45 | PCA | 202.9 | -66.7 | 13.00 | 6 | 25 | 80 |
| CB1833D | 9776 | 1945.6 | -104.64253 | 38.86695 | 8.65 | PCA | 200.3 | -41.8 | 9.90 | 18 | 9 | 100 |
| CB1830A | 9776 | 1949.6 | -104.64282 | 38.86708 | 2.18 | PCA | 163.8 | -62.6 | 4.60 | 8 | 20 | 100 |
| CB1830B | 9776 | 1949.6 | -104.64282 | 38.86708 | 3.48 | PCA | 170.7 | -38.6 | 12.80 | 6 | 40 | 100 |
| CB1830C | 9776 | 1949.6 | -104.64282 | 38.86708 | 2.42 | PCA | 193.6 | -54.3 | 14.10 | 14 | 15 | 80 |
| CB1803A | 9776 | 1950.2 | -104.64283 | 38.86712 | 3.17 | PCA | 148.7 | -47.4 | 16.94 | 16 | 15 | 100 |
| CB1803C | 9776 | 1950.2 | -104.64283 | 38.86712 | 4.11 | PCA | 130.0 | -37.9 | 12.10 | 11 | 30 | 100 |
| CB1803D | 9776 | 1950.2 | -104.64283 | 38.86712 | 7.43 | Fisher | 181.8 | -32.8 | 1.95 | 11 | 27 | 90 |
| CB1831A | 9776 | 1951.1 | -104.64280 | 38.86710 | 2.91 | Fisher | 262.1 | 40.8 | 4.30 | 11 | 0 | 70 |
| CB1831C | 9776 | 1951.1 | -104.64280 | 38.86710 | 3.83 | PCA | 321.1 | 57.9 | 10.90 | 13 | 3 | 60 |
| CB1831D | 9776 | 1951.1 | -104.64280 | 38.86710 | 1.90 | Fisher | 266.9 | 13.4 | 9.74 | 7 | 6 | 30 |
| CB1832A | 9776 | 1952.9 | -104.64279 | 38.86712 | 1.85 | Fisher | 109.6 | -60.8 | 7.92 | 5 | 24 | 40 |
| CB1832D | 9776 | 1952.9 | -104.64279 | 38.86712 | 3.22 | PCA | 180.9 | -47.7 | 11.00 | 7 | 15 | 60 |
| CB1832E | 9776 | 1952.9 | -104.64279 | 38.86712 | 2.16 | PCA | 192.6 | -55.2 | 14.80 | 13 | 24 | 100 |
| CB1802B | 9776 | 1954.3 | -104.64259 | 38.86715 | 93.03 | PCA | 355.1 | 58.9 | 3.40 | 14 | 6 | 55 |
| CB1802C | 9776 | 1954.3 | -104.64259 | 38.86715 | 172.00 | PCA | 355.8 | 53.4 | 3.80 | 11 | 9 | 45 |
| CB1802D | 9776 | 1954.3 | -104.64259 | 38.86715 | 174.00 | PCA | 358.8 | 59.3 | 3.80 | 13 | 9 | 55 |
| CB1801B | 9776 | 1956.7 | -104.64268 | 38.86720 | 77.47 | PCA | 5.1 | 52.7 | 3.80 | 15 | 6 | 60 |
| CB1801C | 9776 | 1956.7 | -104.64268 | 38.86720 | 84.65 | PCA | 354.2 | 53.3 | 3.10 | 13 | 6 | 50 |
| CB1801D | 9776 | 1956.7 | -104.64268 | 38.86720 | 116.60 | PCA | 358.3 | 48.0 | 7.80 | 11 | 27 | 100 |
| CB1744A | 9776 | 1957.5 | -104.64310 | 38.86745 | 210.70 | PCA | 354.6 | 61.8 | 5.70 | 11 | 18 | 60 |
| CB1744B | 9776 | 1957.5 | -104.64310 | 38.86745 | 172.00 | PCA | 341.8 | 65.0 | 5.90 | 13 | 15 | 70 |
| CB1745A | 9776 | 1959.0 | -104.64328 | 38.86761 | 134.40 | PCA | 358.0 | 50.0 | 2.30 | 9 | 6 | 30 |
| CB1745B | 9776 | 1959.0 | -104.64328 | 38.86761 | 110.00 | PCA | 347.3 | 49.0 | 3.70 | 15 | 9 | 70 |
| CB1745C | 9776 | 1959.0 | -104.64328 | 38.86761 | 112.10 | PCA | 354.9 | 58.6 | 2.70 | 15 | 9 | 70 |
| CB1746A | 9776 | 1984.1 | -104.64317 | 38.86859 | 86.30 | PCA | 15.5 | 57.5 | 5.10 | 17 | 12 | 100 |
| CB1746B | 9776 | 1984.1 | -104.64317 | 38.86859 | 69.20 | PCA | 25.6 | 58.8 | 5.30 | 16 | 12 | 100 |
| CB1746C | 9776 | 1984.1 | -104.64317 | 38.86859 | 42.88 | GC | 10.3 | 54.2 | 29.50 | 16 | 15 | 100 |
| CB1872A | Lyco-Luck | *1890.1* | -104.66569 | 38.84979 | 1.37 | Fisher | 11.8 | 50.8 | 7.97 | 11 | 3 | 50 |
| CB1872B | Lyco-Luck | *1890.1* | -104.66569 | 38.84979 | 1.65 | Fisher | 287.7 | -37.4 | 11.11 | 6 | 20 | 50 |
| CB1872D | Lyco-Luck | *1890.1* | -104.66569 | 38.84979 | 1.73 | Fisher | 329.0 | 39.8 | 9.15 | 7 | 0 | 20 |
| CB1873B | Lyco-Luck | *1892.3* | -104.66481 | 38.85047 | 2.36 | PCA | 354.8 | 32.4 | 12.50 | 11 | 3 | 50 |
| CB1873C | Lyco-Luck | *1892.3* | -104.66481 | 38.85047 | 2.30 | PCA | 1.7 | 45.9 | 13.40 | 10 | 3 | 40 |
| CB1873D | Lyco-Luck | *1892.3* | -104.66481 | 38.85047 | 2.88 | PCA | 351.6 | 30.0 | 11.30 | 12 | 6 | 80 |
| CB1820A | Lyco-Luck | 1900.2 | -104.66004 | 38.85321 | 1.25 | GC | 194.5 | -24.5 | 16.40 | 5 | 30 | 80 |
| CB1820C | Lyco-Luck | 1900.2 | -104.66004 | 38.85321 | 0.41 | GC | 193.0 | -22.9 | 12.00 | 4 | 0 | 9 |
| CB1820D | Lyco-Luck | 1900.2 | -104.66004 | 38.85321 | 0.23 | GC | 192.2 | -21.3 | 10.70 | 3 | 60 | 100 |
| CB1807A | Lyco-Luck | *1902.0* | -104.65764 | 38.85345 | 26.26 | GC | 199.1 | -57.4 | 9.70 | 16 | 0 | 55 |
| CB1807B | Lyco-Luck | *1902.0* | -104.65764 | 38.85345 | 8.72 | GC | 174.1 | -69.9 | 13.10 | 18 | 0 | 70 |
| CB1807C | Lyco-Luck | *1902.0* | -104.65764 | 38.85345 | 4.76 | GC | 220.6 | -56.6 | 12.80 | 7 | 0 | 18 |
| CB1806B | Lyco-Luck | *1903* | -104.65737 | 38.85389 | 3.32 | Fisher | 142.5 | 51.6 | 9.96 | 11 | 0 | 30 |
| CB1806C | Lyco-Luck | *1903* | -104.65737 | 38.85389 | 2.10 | Fisher | 146.5 | -32.5 | 8.64 | 12 | 6 | 45 |
| CB1806D | Lyco-Luck | *1903* | -104.65737 | 38.85389 | 0.97 | Fisher | 89.2 | -37.8 | 6.04 | 17 | 6 | 80 |
| CB1749A | Lyco-Luck | *1904* | -104.65747 | 38.85426 | 1.26 | Fisher | 112.4 | -36.7 | 4.85 | 20 | 3 | 100 |
| CB1748A | Lyco-Luck | *1906.0* | -104.65755 | 38.85441 | 1.45 | PCA | 182.0 | -17.3 | 10.50 | 11 | 27 | 90 |
| CB1748C | Lyco-Luck | *1906.0* | -104.65755 | 38.85441 | 1.20 | Fisher | 155.9 | -23.4 | 3.32 | 7 | 6 | 24 |
| CB1748D | Lyco-Luck | *1906.0* | -104.65755 | 38.85441 | 0.93 | Fisher | 193.2 | -11.2 | 7.86 | 12 | 6 | 45 |
| CB1747A | Lyco-Luck | 1907.6 | -104.65741 | 38.85453 | 9.65 | Fisher | 168.1 | -59.1 | 5.87 | 13 | 6 | 50 |
| CB1747B | Lyco-Luck | 1907.6 | -104.65741 | 38.85453 | 1.69 | Fisher | 163.0 | -39.0 | 9.77 | 19 | 6 | 100 |
| CB1823B | Bambino Canyon | 1931.7 | -104.58900 | 38.85736 | 69.21 | GC | 199.1 | -64.6 | 2.90 | 14 | 0 | 45 |
| CB1823C | Bambino Canyon | 1931.7 | -104.58900 | 38.85736 | 49.20 | GC | 195.8 | -62.0 | 7.10 | 11 | 0 | 30 |
| CB1823E | Bambino Canyon | 1931.7 | -104.58900 | 38.85736 | 51.24 | GC | 197.8 | -63.2 | 8.30 | 15 | 3 | 55 |
| CB1824A | Bambino Canyon | 1934.9 | -104.58840 | 38.85725 | 11.30 | PCA | 139.3 | -11.0 | 13.90 | 20 | 3 | 100 |
| CB1824B | Bambino Canyon | 1934.9 | -104.58840 | 38.85725 | 6.74 | PCA | 147.4 | -18.9 | 15.30 | 9 | 24 | 70 |
| CB1824C | Bambino Canyon | 1934.9 | -104.58840 | 38.85725 | 6.67 | PCA | 177.6 | -21.2 | 13.30 | 15 | 18 | 100 |
| CB1825A | Bambino Canyon | 1943.6 | -104.58768 | 38.85799 | 5.03 | Fisher | 161.4 | -42.8 | 3.45 | 15 | 12 | 80 |
| CB1825B | Bambino Canyon | 1943.6 | -104.58768 | 38.85799 | 4.06 | PCA | 110.5 | -37.9 | 13.00 | 17 | 3 | 70 |
| CB1825C | Bambino Canyon | 1943.6 | -104.58768 | 38.85799 | 5.30 | Fisher | 146.4 | -32.7 | 3.12 | 15 | 12 | 80 |
| CB1826A | Bambino Canyon | 1948.8 | -104.58770 | 38.85809 | 1.90 | Fisher | 104.0 | -39.0 | 9.00 | 7 | 40 | 80 |

| Sample | Section | Elev (m) | Lon(º) | Lat(º) | NRM Intensity (mA/m) | Method | Dec (°) | Inc (°) | MAD/α95 (°) | n points | Start (mT) | End (mT) |
|---|---|---|---|---|---|---|---|---|---|---|---|---|
| CB1826C | Bambino Canyon | 1948.8 | -104.58770 | 38.85809 | 2.55 | Fisher | 148.3 | -67.5 | 6.01 | 13 | 18 | 80 |
| CB1826D | Bambino Canyon | 1948.8 | -104.58770 | 38.85809 | 1.05 | Fisher | 114.8 | -59.2 | 9.07 | 5 | 9 | 21 |
| CB1827A | Bambino Canyon | 1955.3 | -104.58768 | 38.85823 | 6.67 | PCA | 173.1 | -54.4 | 8.80 | 13 | 21 | 90 |
| CB1827B | Bambino Canyon | 1955.3 | -104.58768 | 38.85823 | 7.18 | PCA | 178.6 | -53.7 | 7.80 | 14 | 18 | 90 |
| CB1827C | Bambino Canyon | 1955.3 | -104.58768 | 38.85823 | 3.69 | PCA | 154.1 | -52.0 | 7.90 | 12 | 12 | 100 |
| CB1809A | Bambino Canyon | | -104.64893 | 38.86377 | 3.48 | GC | 109.0 | -36.4 | 7.10 | 5 | 0 | 12 |
| CB1809C | Bambino Canyon | | -104.64893 | 38.86377 | 7.59 | GC | 125.1 | -40.2 | 13.90 | 14 | 3 | 50 |
| CB1809D | Bambino Canyon | | -104.64893 | 38.86377 | 7.04 | GC | 99.2 | -51.3 | 12.60 | 9 | 0 | 24 |
| CB1828A | Bambino Canyon | 1959.6 | -104.64663 | 38.86328 | 84.51 | PCA | 8.2 | 38.4 | 3.90 | 13 | 6 | 60 |
| CB1828C | Bambino Canyon | 1959.6 | -104.64663 | 38.86328 | 476.10 | PCA | 16.0 | 41.8 | 4.90 | 9 | 9 | 35 |
| CB1828D | Bambino Canyon | 1959.6 | -104.64663 | 38.86328 | 516.90 | PCA | 13.0 | 35.9 | 4.80 | 13 | 9 | 55 |
| CB1829B | Bambino Canyon | 1962.7 | -104.64678 | 38.86333 | 69.96 | PCA | 33.0 | 70.1 | 4.40 | 8 | 9 | 30 |
| CB1829C | Bambino Canyon | 1962.7 | -104.64678 | 38.86333 | 94.27 | PCA | 32.6 | 74.1 | 11.40 | 10 | 18 | 55 |
| CB1829D | Bambino Canyon | 1962.7 | -104.64678 | 38.86333 | 118.90 | PCA | 68.9 | 46.9 | 7.90 | 14 | 12 | 70 |
| CB1805A | Bambino Canyon | | -104.64938 | 38.86433 | 15.59 | PCA | 22.1 | 55.9 | 5.90 | 5 | 15 | 27 |
| CB1805C | Bambino Canyon | | -104.64938 | 38.86433 | 1.67 | Fisher | 22.0 | 55.8 | 4.29 | 17 | 3 | 60 |
| CB1805F | Bambino Canyon | | -104.64938 | 38.86433 | 120.70 | PCA | 4.3 | 65.6 | 3.30 | 11 | 3 | 35 |
| CB1810A | Hawkins | 1901.0 | -104.58737 | 38.83270 | 6.77 | Fisher | 176.5 | -16.1 | 10.78 | 8 | 3 | 24 |
| CB1810B | Hawkins | 1901.0 | -104.58737 | 38.83270 | 3.31 | Fisher | 165.3 | -13.8 | 16.01 | 6 | 6 | 21 |
| CB1810D | Hawkins | 1901.0 | -104.58737 | 38.83270 | 2.50 | Fisher | 154.4 | -39.9 | 5.50 | 15 | 12 | 80 |
| CB1811B | Hawkins | 1907.9 | -104.58495 | 38.83414 | 27.12 | GC | 102.0 | -36.9 | 18.70 | 6 | 24 | 45 |
| CB1811C | Hawkins | 1907.9 | -104.58495 | 38.83414 | 30.26 | GC | 124.3 | -63.4 | 3.80 | 3 | 40 | 50 |
| CB1811D | Hawkins | 1907.9 | -104.58495 | 38.83414 | 31.80 | GC | 14.9 | -56.8 | 15.80 | 4 | 50 | 70 |
| CB1812A | Hawkins | 1937.6 | -104.57927 | 38.83548 | 35.68 | GC | 77.2 | -46.6 | 14.10 | 12 | 27 | 100 |
| CB1812C | Hawkins | 1937.6 | -104.57927 | 38.83548 | 33.22 | GC | 75.9 | -46.3 | 8.80 | 4 | 50 | 70 |
| CB1812D | Hawkins | 1937.6 | -104.57927 | 38.83548 | 25.28 | GC | 70.7 | -49.4 | 5.50 | 10 | 18 | 50 |
| CB1814A | Hawkins | 1951.4 | -104.57520 | 38.83759 | 2.47 | Fisher | 148.4 | -75.0 | 2.94 | 20 | 3 | 100 |
| CB1814B | Hawkins | 1951.4 | -104.57520 | 38.83759 | 2.10 | Fisher | 92.5 | -68.3 | 8.64 | 14 | 0 | 45 |
| CB1814C | Hawkins | 1951.4 | -104.57520 | 38.83759 | 14.73 | Fisher | 132.0 | -38.7 | 27.41 | 4 | 70 | 100 |
| CB1813B | Hawkins | 1955.1 | -104.57549 | 38.83756 | 5.83 | Fisher | 185.4 | -51.3 | 3.20 | 20 | 3 | 100 |
| CB1813C | Hawkins | 1955.1 | -104.57549 | 38.83756 | 3.38 | Fisher | 221.7 | -33.4 | 2.93 | 20 | 3 | 100 |
| CB1813D | Hawkins | 1955.1 | -104.57549 | 38.83756 | 1.05 | Fisher | 130.3 | -50.6 | 6.43 | 8 | 6 | 27 |
| CB1835B | Waste Management | 1944.5 | -104.58053 | 38.84136 | 1.78 | Fisher | 149.1 | -62.8 | 8.40 | 11 | 6 | 40 |
| CB1835C | Waste Management | 1944.5 | -104.58053 | 38.84136 | 0.30 | Fisher | 178.8 | -72.7 | 7.23 | 11 | 3 | 80 |
| CB1835D | Waste Management | 1944.5 | -104.58053 | 38.84136 | 1.11 | Fisher | 157.6 | -43.9 | 5.57 | 14 | 3 | 50 |
| CB1866A | Waste Management | 1946.5 | -104.58050 | 38.84138 | 8.08 | Fisher | 157.2 | -47.0 | 4.54 | 9 | 20 | 100 |
| CB1866B | Waste Management | 1946.5 | -104.58050 | 38.84138 | 4.28 | Fisher | 166.1 | -42.9 | 3.88 | 8 | 15 | 60 |
| CB1866C | Waste Management | 1946.5 | -104.58050 | 38.84138 | 12.20 | PCA | 59.6 | -48.5 | 7.00 | 10 | 12 | 80 |
| CB1866D | Waste Management | 1946.5 | -104.58050 | 38.84138 | 3.41 | PCA | 173.1 | -47.4 | 11.80 | 12 | 6 | 80 |
| CB1865A | Waste Management | 1948.8 | -104.58043 | 38.84136 | 1.87 | Fisher | 141.8 | -35.5 | 4.37 | 13 | 6 | 100 |
| CB1865B | Waste Management | 1948.8 | -104.58043 | 38.84136 | 0.55 | Fisher | 167.8 | -51.7 | 5.24 | 13 | 6 | 100 |
| CB1865D | Waste Management | 1948.8 | -104.58043 | 38.84136 | 1.25 | Fisher | 136.8 | -17.7 | 2.22 | 7 | 15 | 50 |
| CB1858A | Waste Management | 1952.1 | -104.58042 | 38.84144 | 1.67 | Fisher | 26.9 | -27.2 | 6.82 | 13 | 6 | 80 |
| CB1858C | Waste Management | 1952.1 | -104.58042 | 38.84144 | 2.35 | Fisher | 171.9 | 39.3 | 12.83 | 12 | 18 | 100 |
| CB1858E | Waste Management | 1952.1 | -104.58042 | 38.84144 | 2.48 | Fisher | 280.0 | -5.2 | 7.11 | 9 | 9 | 60 |
| CB1867C | Waste Management | 1954.6 | -104.58034 | 38.84142 | 28.40 | PCA | 12.5 | 62.6 | 4.30 | 9 | 9 | 35 |
| CB1867D | Waste Management | 1954.6 | -104.58034 | 38.84142 | 49.30 | PCA | 5.3 | 57.5 | 2.70 | 10 | 3 | 40 |
| CB1867E | Waste Management | 1954.6 | -104.58034 | 38.84142 | 32.20 | PCA | 21.2 | 60.7 | 4.70 | 7 | 15 | 50 |
| CB1857A | Waste Management | 1962.1 | -104.57928 | 38.84170 | 55.97 | PCA | 359.8 | 55.4 | 4.00 | 11 | 9 | 50 |
| CB1857B | Waste Management | 1962.1 | -104.57928 | 38.84170 | 47.96 | PCA | 12.6 | 53.0 | 7.50 | 6 | 9 | 30 |
| CB1857C | Waste Management | 1962.1 | -104.57928 | 38.84170 | 47.51 | PCA | 359.9 | 53.3 | 6.30 | 12 | 9 | 50 |
| CB1834A | Waste Management | 1965.5 | -104.57942 | 38.84185 | 32.75 | PCA | 358.9 | 58.4 | 9.20 | 13 | 9 | 100 |
| CB1834C | Waste Management | 1965.5 | -104.57942 | 38.84185 | 22.24 | PCA | 14.8 | 60.6 | 14.50 | 16 | 12 | 90 |
| CB1834D | Waste Management | 1965.5 | -104.57942 | 38.84185 | 28.17 | PCA | 3.6 | 49.2 | 5.90 | 17 | 3 | 70 |
| CB1856B | Waste Management | *1973* | -104.58229 | 38.84597 | 100.30 | PCA | 273.3 | 60.3 | 3.30 | 16 | 0 | 55 |
| CB1856C | Waste Management | *1973* | -104.58229 | 38.84597 | 67.94 | PCA | 354.9 | 67.2 | 3.70 | 15 | 3 | 55 |
| CB1856D | Waste Management | *1973* | -104.58229 | 38.84597 | 99.28 | PCA | 65.3 | 59.1 | 7.60 | 9 | 9 | 60 |
| CB1855A | Waste Management | *1988* | -104.58209 | 38.84647 | 8.84 | Fisher | 3.9 | 32.5 | 7.85 | 13 | 18 | 80 |
| CB1855B | Waste Management | *1988* | -104.58209 | 38.84647 | 7.65 | PCA | 339.8 | 45.9 | 10.50 | 11 | 9 | 100 |
| CB1855C | Waste Management | *1988* | -104.58209 | 38.84647 | 8.79 | PCA | 328.7 | 57.4 | 13.40 | 16 | 3 | 60 |
| CB1859A | 9789 | 1872.6 | -104.62469 | 38.84311 | 12.29 | PCA | 290.0 | 57.6 | 13.50 | 7 | 15 | 40 |
| CB1859B | 9789 | 1872.6 | -104.62469 | 38.84311 | 8.88 | PCA | 4.6 | 78.2 | 13.40 | 15 | 9 | 70 |
| CB1859C | 9789 | 1872.6 | -104.62469 | 38.84311 | 4.21 | PCA | 5.0 | 40.7 | 13.10 | 14 | 0 | 100 |
| CB1859E | 9789 | 1872.6 | -104.62469 | 38.84311 | 5.39 | PCA | 10.7 | 57.8 | 13.20 | 11 | 3 | 60 |
| CB1860A | 9789 | 1876.8 | -104.62784 | 38.84634 | 1.87 | Fisher | 265.6 | 69.5 | 10.52 | 13 | 0 | 40 |
| CB1860B | 9789 | 1876.8 | -104.62784 | 38.84634 | 1.48 | Fisher | 22.9 | 55.8 | 6.71 | 9 | 0 | 30 |
| CB1860C | 9789 | 1876.8 | -104.62784 | 38.84634 | 2.14 | Fisher | 1.1 | 52.6 | 5.96 | 6 | 3 | 20 |
| CB1860D | 9789 | 1876.8 | -104.62784 | 38.84634 | 1.50 | Fisher | 352.9 | 53.0 | 8.78 | 5 | 3 | 15 |
| CB1861A | 9789 | 1883.8 | -104.62723 | 38.84787 | 2.63 | GC | 183.6 | -40.2 | 13.70 | 6 | 12 | 40 |
| CB1861B | 9789 | 1883.8 | -104.62723 | 38.84787 | 3.09 | GC | 176.0 | -47.7 | 13.40 | 16 | 3 | 80 |
| CB1861C | 9789 | 1883.8 | -104.62723 | 38.84787 | 3.77 | PCA | 108.3 | -52.8 | 13.70 | 9 | 27 | 100 |
| CB1861D | 9789 | 1883.8 | -104.62723 | 38.84787 | 2.78 | GC | 186.5 | -21.2 | 17.70 | 8 | 3 | 30 |
| CB1863A | 9789 | 1902.3 | -104.63521 | 38.85091 | 1.30 | PCA | 150.4 | -63.0 | 6.20 | 9 | 27 | 70 |
| CB1863B | 9789 | 1902.3 | -104.63521 | 38.85091 | 0.64 | PCA | 162.7 | -44.3 | 22.70 | 8 | 21 | 50 |
| CB1863C | 9789 | 1902.3 | -104.63521 | 38.85091 | 1.27 | PCA | 165.1 | -37.7 | 4.20 | 8 | 20 | 100 |
| CB1863D | 9789 | 1902.3 | -104.63521 | 38.85091 | 0.87 | Fisher | 119.7 | -70.6 | 3.86 | 7 | 25 | 100 |
| CB1864A | 9789 | 1905.3 | -104.63536 | 38.85166 | 4.37 | PCA | 161.3 | -43.6 | 8.80 | 7 | 30 | 100 |

| Sample | Section | Elev (m) | Lon(º) | Lat(º) | NRM Intensity (mA/m) | Method | Dec (°) | Inc (°) | MAD/α95 (°) | n points | Start (mT) | End (mT) |
|---|---|---|---|---|---|---|---|---|---|---|---|---|
| CB1864C | 9789 | 1905.3 | -104.63536 | 38.85166 | 0.65 | Fisher | 140.9 | -21.4 | 3.51 | 9 | 6 | 100 |
| CB1864D | 9789 | 1905.3 | -104.63536 | 38.85166 | 4.73 | PCA | 145.0 | -11.5 | 9.80 | 9 | 24 | 100 |
| CB1837A | 9789 | 1918.2 | -104.63599 | 38.85486 | 25.58 | PCA | 358.6 | 63.6 | 6.60 | 14 | 0 | 45 |
| CB1837B | 9789 | 1918.2 | -104.63599 | 38.85486 | 9.02 | GC | 171.6 | -48.8 | 15.30 | 17 | 0 | 60 |
| CB1837D | 9789 | 1918.2 | -104.63599 | 38.85486 | 19.61 | GC | 161.3 | -26.7 | 7.60 | 3 | 50 | 80 |
| CB1837E | 9789 | 1918.2 | -104.63599 | 38.85486 | 39.18 | PCA | 353.8 | 48.5 | 4.50 | 7 | 12 | 40 |
| CB1838A | 9789 | 1925.6 | -104.63670 | 38.85554 | 0.74 | Fisher | 59.0 | -72.8 | 2.62 | 15 | 6 | 60 |
| CB1838C | 9789 | 1925.6 | -104.63670 | 38.85554 | 0.94 | Fisher | 314.6 | -48.0 | 5.26 | 14 | 9 | 60 |
| CB1838D | 9789 | 1925.6 | -104.63670 | 38.85554 | 0.38 | Fisher | 302.9 | -64.9 | 4.30 | 7 | 15 | 60 |
| CB1839A | 9789 | 1935.5 | -104.63637 | 38.85745 | 9.99 | PCA | 133.7 | -27.6 | 13.50 | 6 | 40 | 100 |
| CB1839B | 9789 | 1935.5 | -104.63637 | 38.85745 | 29.65 | GC | 281.6 | -48.1 | 7.30 | 12 | 9 | 50 |
| CB1839C | 9789 | 1935.5 | -104.63637 | 38.85745 | 14.50 | GC | 279.8 | -48.0 | 4.20 | 7 | 0 | 20 |
| CB1840A | 9789 | 1941.6 | -104.63643 | 38.85744 | 0.42 | Fisher | 141.9 | -31.1 | 9.75 | 15 | 6 | 100 |
| CB1840B | 9789 | 1941.6 | -104.63643 | 38.85744 | 2.08 | Fisher | 288.2 | 33.9 | 16.90 | 6 | 3 | 18 |
| CB1840C | 9789 | 1941.6 | -104.63643 | 38.85744 | 1.03 | PCA | 204.3 | -52.8 | 15.80 | 6 | 30 | 100 |
| CB1841B | 9789 | 1946.7 | -104.63589 | 38.85813 | 38.34 | GC | 193.2 | -53.7 | 10.40 | 10 | 6 | 35 |
| CB1841C | 9789 | 1946.7 | -104.63589 | 38.85813 | 67.06 | GC | 191.6 | -57.2 | 6.90 | 12 | 0 | 35 |
| CB1841D | 9789 | 1946.7 | -104.63589 | 38.85813 | 27.39 | GC | 190.0 | -60.2 | 10.80 | 10 | 6 | 35 |
| CB1842A | 9789 | 1946.7 | -104.63592 | 38.85830 | 1.51 | Fisher | 197.2 | -61.4 | 5.03 | 12 | 3 | 40 |
| CB1842B | 9789 | 1946.7 | -104.63592 | 38.85830 | 1.50 | Fisher | 218.2 | -44.8 | 7.76 | 5 | 6 | 20 |
| CB1842C | 9789 | 1946.7 | -104.63592 | 38.85830 | 1.14 | Fisher | 227.1 | 28.1 | 11.21 | 12 | 6 | 45 |
| CB1842D | 9789 | 1946.7 | -104.63592 | 38.85830 | 1.15 | Fisher | 335.4 | -65.6 | 19.08 | 8 | 9 | 60 |
| CB1843B | 9789 | 1951.8 | -104.63573 | 38.85839 | 5.36 | Fisher | 207.8 | -65.1 | 7.85 | 8 | 12 | 35 |
| CB1843C | 9789 | 1951.8 | -104.63573 | 38.85839 | 3.88 | Fisher | 151.9 | -56.1 | 3.80 | 12 | 6 | 100 |
| CB1843D | 9789 | 1951.8 | -104.63573 | 38.85839 | 2.11 | Fisher | 227.3 | -59.1 | 4.70 | 13 | 3 | 45 |
| CB1844A | 9789 | 1954.9 | -104.63583 | 38.85841 | 5.99 | GC | 179.4 | -65.1 | 17.50 | 13 | 0 | 60 |
| CB1844C | 9789 | 1954.9 | -104.63583 | 38.85841 | 11.34 | GC | 176.1 | -66.6 | 8.30 | 14 | 0 | 45 |
| CB1844E | 9789 | 1954.9 | -104.63583 | 38.85841 | 14.59 | GC | 173.5 | -62.0 | 18.90 | 15 | 0 | 50 |
| CB1845B | 9789 | 1958.9 | -104.63571 | 38.85855 | 2.10 | PCA | 178.0 | -45.9 | 15.30 | 6 | 30 | 100 |
| CB1845C | 9789 | 1958.9 | -104.63571 | 38.85855 | 0.66 | PCA | 147.0 | -46.8 | 15.00 | 7 | 30 | 100 |
| CB1845D | 9789 | 1958.9 | -104.63571 | 38.85855 | 3.68 | PCA | 171.0 | -47.0 | 10.40 | 12 | 21 | 100 |
| CB1846A | 9789 | 1962.6 | -104.63584 | 38.85866 | 113.20 | PCA | 7.1 | 62.9 | 2.30 | 12 | 9 | 50 |
| CB1846D | 9789 | 1962.6 | -104.63584 | 38.85866 | 29.48 | PCA | 352.8 | 57.3 | 2.80 | 9 | 9 | 60 |
| CB1846E | 9789 | 1962.6 | -104.63584 | 38.85866 | 52.52 | PCA | 3.9 | 60.7 | 3.80 | 13 | 9 | 55 |
| CB1847B | 9789 | 1965.8 | -104.63585 | 38.85870 | 127.70 | PCA | 351.7 | 53.7 | 2.10 | 9 | 6 | 50 |
| CB1847D | 9789 | 1965.8 | -104.63585 | 38.85870 | 101.00 | PCA | 346.2 | 55.4 | 3.10 | 16 | 9 | 80 |
| CB1847E | 9789 | 1965.8 | -104.63585 | 38.85870 | 58.45 | PCA | 350.7 | 48.2 | 3.60 | 15 | 12 | 80 |
| CB1848B | 9789 | 1971.2 | -104.63490 | 38.85886 | 3.98 | PCA | 17.2 | 52.7 | 6.20 | 9 | 15 | 100 |
| CB1848C | 9789 | 1971.2 | -104.63490 | 38.85886 | 4.25 | PCA | 33.2 | 62.0 | 7.50 | 16 | 9 | 80 |
| CB1848D | 9789 | 1971.2 | -104.63490 | 38.85886 | 4.28 | PCA | 357.1 | 58.0 | 5.90 | 9 | 12 | 40 |
| CB1850B | Kunstle | 2029.0 | -104.59666 | 38.86847 | 4.87 | PCA | 316.8 | 45.4 | 10.40 | 16 | 15 | 100 |
| CB1850C | Kunstle | 2029.0 | -104.59666 | 38.86847 | 4.56 | PCA | 330.5 | 59.7 | 14.00 | 13 | 18 | 80 |
| CB1850D | Kunstle | 2029.0 | -104.59666 | 38.86847 | 4.99 | PCA | 0.5 | 63.4 | 10.50 | 13 | 3 | 100 |
| CB1851B | Kunstle | 2036.0 | -104.59621 | 38.86927 | 3.04 | PCA | 346.0 | 63.7 | 13.30 | 12 | 18 | 70 |
| CB1851C | Kunstle | 2036.0 | -104.59621 | 38.86927 | 3.42 | PCA | 1.4 | 46.4 | 8.00 | 9 | 9 | 60 |
| CB1851D | Kunstle | 2036.0 | -104.59621 | 38.86927 | 2.45 | PCA | 8.9 | 67.5 | 14.60 | 10 | 12 | 40 |
| CB1868C | Kunstle | 2036.9 | -104.59613 | 38.86965 | 2.90 | PCA | 351.2 | 67.3 | 9.60 | 15 | 3 | 100 |
| CB1868D | Kunstle | 2036.9 | -104.59613 | 38.86965 | 6.96 | PCA | 343.7 | 65.8 | 9.00 | 12 | 9 | 100 |
| CB1868E | Kunstle | 2036.9 | -104.59613 | 38.86965 | 5.79 | PCA | 334.5 | 30.9 | 5.70 | 14 | 3 | 600 |
| CB1869A | Kunstle | 2040.6 | -104.59588 | 38.86971 | 2.16 | Fisher | 150.6 | -22.7 | 2.51 | 10 | 3 | 40 |
| CB1869B | Kunstle | 2040.6 | -104.59588 | 38.86971 | 1.69 | Fisher | 150.6 | -33.9 | 3.66 | 13 | 6 | 100 |
| CB1869D | Kunstle | 2040.6 | -104.59588 | 38.86971 | 1.28 | PCA | 168.8 | -53.9 | 14.50 | 9 | 15 | 80 |
| CB1853B | Kunstle | 2042.4 | -104.59593 | 38.86977 | 7.87 | Fisher | 214.8 | -24.9 | 4.99 | 9 | 21 | 55 |
| CB1853C | Kunstle | 2042.4 | -104.59593 | 38.86977 | 10.50 | Fisher | 149.7 | -30.7 | 10.62 | 6 | 25 | 80 |
| CB1853D | Kunstle | 2042.4 | -104.59593 | 38.86977 | 1.69 | PCA | 171.7 | -33.3 | 11.20 | 6 | 30 | 100 |
| CB1853E | Kunstle | 2042.4 | -104.59593 | 38.86977 | 3.87 | PCA | 168.3 | -27.8 | 11.40 | 5 | 30 | 80 |
| CB1870B | Kunstle | 2043.4 | -104.59591 | 38.86977 | 14.20 | Fisher | 151.5 | -58.1 | 6.37 | 11 | 18 | 100 |
| CB1870C | Kunstle | 2043.4 | -104.59591 | 38.86977 | 14.00 | Fisher | 152.2 | -41.3 | 10.27 | 7 | 27 | 80 |
| CB1870D | Kunstle | 2043.4 | -104.59591 | 38.86977 | 4.36 | Fisher | 164.5 | -30.8 | 13.82 | 10 | 21 | 100 |
| CB1870E | Kunstle | 2043.4 | -104.59591 | 38.86977 | 6.87 | Fisher | 205.8 | -57.2 | 13.56 | 8 | 27 | 70 |
| CB1854A | Kunstle | 2043.7 | -104.59591 | 38.86980 | 38.19 | PCA | 351.8 | 51.0 | 14.90 | 7 | 18 | 40 |
| CB1854B | Kunstle | 2043.7 | -104.59591 | 38.86980 | 28.31 | PCA | 349.0 | 60.2 | 7.80 | 13 | 9 | 55 |
| CB1854D | Kunstle | 2043.7 | -104.59591 | 38.86980 | 32.53 | PCA | 343.3 | 57.8 | 6.60 | 8 | 12 | 60 |
| CB1871A | Kunstle | 2045.1 | -104.59585 | 38.86979 | 1.05 | GC | 135.5 | -49.3 | 26.90 | 9 | 9 | 50 |
| CB1871B | Kunstle | 2045.1 | -104.59585 | 38.86979 | 1.15 | GC | 134.6 | -46.2 | 14.40 | 9 | 9 | 50 |
| CB1871C | Kunstle | 2045.1 | -104.59585 | 38.86979 | 0.92 | GC | 135.4 | -47.8 | 13.00 | 11 | 6 | 60 |
| CB1871D | Kunstle | 2045.1 | -104.59585 | 38.86979 | 0.51 | GC | 140.9 | -52.6 | 14.10 | 10 | 3 | 50 |

**Table S2:** Sample data for all paleomagnetic sites that exhibited stable demagnetization behavior. Samples are sorted by stratigraphic superposition in each section. **Elev–** Sample DGPS elevation, italicized are estimated values and blank could not be determined (see methods); **Lon/Lat–** Sample DGPS longitude and latitude relative to WGS 84 Datum; **NRM Intensity–** intensity of NRM prior to demagnetization (mA/m); **Meth–** refers to method used to define ChRM (PCA, Fisher mean, great circle); **Dec/Inc–** Sample ChRM declination and inclination for PCA and Fisher means and declination and inclination of point along great circle path closest to site mean;**MAD/α95–** maximum angular deviation (PCA, GC) or α95 (fisher mean) of sample ChRM; **n points–** number of steps used to calculate ChRM; **Start–** starting demagnetization step in (mT) (CB1703B characterized by thermal steps in °C) used in ChRM determination; **End–** ending demagnetization step in (mT) (CB1703B characterized by thermal steps in °C) used in ChRM determination.

| Sample | | CB1703A | | | CB1711C | | | CB1734B | | | CB1821D | | | CB1824B | | | CB1827B | | | CB1869B | |
|---|---|---|---|---|---|---|---|---|---|---|---|---|---|---|---|---|---|---|---|---|---|
| Field (mT) | Avg X | Avg Y | Avg Z | Avg X | Avg Y | Avg Z | Avg X | Avg Y | Avg Z | Avg X | Avg Y | Avg Z | Avg X | Avg Y | Avg Z | Avg X | Avg Y | Avg Z | Avg X | Avg Y | Avg Z |
| 0 | -1.40E-03 | 6.98E-04 | -1.41E-03 | 3.11E-03 | 2.64E-04 | 1.49E-03 | -3.45E-03 | 7.52E-04 | 2.31E-03 | -8.71E-05 | 1.02E-03 | -5.94E-04 | -1.49E-03 | 6.26E-04 | -4.47E-03 | 3.48E-03 | 2.29E-03 | -5.88E-03 | 7.50E-04 | 6.66E-04 | -1.42E-04 |
| 20 | 5.74E+00 | 7.71E-02 | 3.46E-01 | 1.02E+00 | -8.86E-02 | 1.32E-01 | 5.93E+00 | 7.47E-01 | -3.25E-01 | 1.22E-01 | -6.54E-04 | -3.70E-03 | 1.35E+00 | -1.14E-01 | 7.83E-02 | 4.99E-01 | 5.52E-02 | 2.73E-02 | 2.82E-01 | -2.38E-02 | -2.40E-02 |
| 50 | 2.01E+01 | -5.61E-02 | 2.87E+00 | 3.35E+00 | 5.14E-01 | 9.05E-02 | 2.36E+01 | -1.65E+00 | -2.47E+00 | 2.69E-01 | -2.84E-02 | -1.69E-03 | 3.16E+00 | 5.69E-02 | 4.31E-01 | 1.24E+00 | 8.21E-02 | 8.66E-02 | 8.22E-01 | 2.83E-02 | -7.11E-02 |
| 80 | 2.70E+01 | 1.21E+00 | 1.53E+00 | 4.61E+00 | -8.25E-02 | 2.94E-01 | 3.27E+01 | -9.14E-01 | -8.37E-01 | 3.47E-01 | -2.74E-02 | 3.65E-03 | 4.06E+00 | 1.87E-01 | 2.25E-01 | 1.65E+00 | 1.02E-02 | 5.72E-03 | 1.17E+00 | 2.27E-02 | 2.28E-02 |
| 110 | 2.90E+01 | 3.40E+00 | 2.75E+00 | 5.11E+00 | -4.24E-01 | 6.43E-01 | 3.53E+01 | 8.17E+00 | -6.51E+00 | 3.72E-01 | -2.15E-02 | -2.18E-03 | 4.42E+00 | 5.78E-01 | 1.60E-01 | 1.88E+00 | -9.32E-02 | -8.98E-02 | 1.40E+00 | 2.00E-02 | 1.87E-02 |
| 150 | 3.02E+01 | -1.54E+00 | 4.31E-02 | 5.62E+00 | -3.80E-03 | -1.06E-02 | 3.85E+01 | 1.70E-01 | -2.64E+00 | 4.10E-01 | -4.67E-02 | -1.56E-02 | 4.64E+00 | 1.50E-03 | 1.06E+00 | 2.05E+00 | -1.07E-01 | 2.03E-01 | 1.60E+00 | -2.17E-02 | 9.08E-03 |
| 200 | 3.06E+01 | -6.91E-01 | 3.03E+00 | 5.76E+00 | -7.54E-02 | 4.59E-01 | 3.91E+01 | -7.86E-01 | -5.07E+00 | 4.26E-01 | 2.55E-02 | -1.13E-02 | 5.10E+00 | 7.73E-01 | 4.40E-02 | 2.23E+00 | -5.35E-02 | 2.53E-02 | 1.71E+00 | -2.33E-01 | 3.07E-03 |
| 300 | 3.04E+01 | 2.57E+00 | 1.03E+00 | 5.98E+00 | 1.22E-01 | 2.41E-01 | 4.01E+01 | 1.47E+00 | -1.43E+00 | 4.50E-01 | -3.23E-02 | 1.24E-02 | 5.45E+00 | -3.98E-01 | 1.32E-01 | 2.35E+00 | -1.17E-01 | -1.26E-01 | 1.89E+00 | -5.16E-02 | -3.18E-02 |
| 400 | 3.09E+01 | -1.97E-01 | 2.55E+00 | 6.02E+00 | -1.75E-01 | 6.55E-01 | 4.01E+01 | 7.15E-01 | 8.57E-01 | 4.60E-01 | -6.08E-02 | -6.68E-03 | 5.60E+00 | 4.15E-01 | 6.31E-01 | 2.42E+00 | 3.93E-01 | 9.00E-02 | 2.00E+00 | -1.17E-01 | -1.59E-02 |
| 500 | 3.09E+01 | 1.90E-01 | 2.37E+00 | 6.04E+00 | 4.80E-02 | 4.98E-01 | 3.95E+01 | 5.31E+00 | -3.65E+00 | 4.71E-01 | -1.96E-03 | -2.53E-03 | 5.71E+00 | -8.97E-01 | -1.02E-01 | 2.52E+00 | -4.83E-02 | 7.75E-02 | 2.06E+00 | 1.55E-01 | -7.60E-02 |
| 600 | 3.12E+01 | 4.65E-01 | 3.98E+00 | 5.99E+00 | 3.18E-01 | 8.23E-01 | 3.98E+01 | -2.16E-01 | -2.68E+00 | 4.78E-01 | 2.15E-02 | -1.96E-03 | 5.83E+00 | -6.93E-01 | -2.59E-01 | 2.56E+00 | 6.37E-02 | 1.22E-01 | 2.10E+00 | 3.15E-02 | -3.34E-02 |
| 700 | 3.12E+01 | 1.04E-02 | 2.98E+00 | 6.18E+00 | 3.12E-01 | 5.79E-01 | 4.05E+01 | 9.96E-01 | -2.33E+00 | 4.85E-01 | -2.22E-02 | -2.28E-03 | 5.99E+00 | 7.85E-02 | 6.76E-01 | 2.59E+00 | 2.42E-02 | 2.01E-01 | 2.12E+00 | -1.30E-01 | -1.50E-01 |
| 800 | 3.05E+01 | 4.18E+00 | 2.77E+00 | 6.09E+00 | -5.75E-01 | 8.04E-01 | 3.92E+01 | 1.15E+01 | 2.14E+00 | 4.80E-01 | -3.38E-02 | -1.04E-02 | 5.98E+00 | 4.21E-01 | 1.00E+00 | 2.62E+00 | -1.33E-01 | 3.43E-01 | 2.16E+00 | -1.09E-01 | 4.46E-02 |
| 900 | 3.12E+01 | -1.15E+00 | 1.90E+00 | 6.27E+00 | -8.57E-02 | 7.79E-01 | 3.98E+01 | 7.39E-01 | -6.75E+00 | 4.86E-01 | -4.07E-02 | 1.73E-02 | 6.13E+00 | -6.61E-01 | 7.99E-01 | 2.68E+00 | -2.38E-01 | 1.54E-01 | 2.18E+00 | -1.92E-02 | 7.11E-02 |
| 1000 | 3.10E+01 | 8.88E-01 | 3.85E+00 | 6.23E+00 | -7.38E-03 | 9.43E-01 | 4.02E+01 | 7.02E+00 | -3.13E+00 | 4.81E-01 | -2.12E-02 | 2.91E-02 | 6.13E+00 | 8.99E-01 | 4.85E-01 | 2.64E+00 | -2.09E-01 | 3.43E-01 | 2.22E+00 | 1.19E-01 | 1.83E-02 |
| 1100 | 3.10E+01 | -1.99E+00 | 3.41E+00 | 6.07E+00 | 7.05E-01 | 1.08E+00 | 4.07E+01 | 1.25E+00 | -3.07E+00 | 4.85E-01 | -2.90E-02 | -2.95E-02 | 6.15E+00 | 8.24E-02 | 4.13E-01 | 2.62E+00 | 1.19E-01 | 4.55E-01 | 2.21E+00 | 4.05E-02 | -8.83E-02 |
| -100 | -2.67E+01 | -5.01E-01 | -4.88E+00 | -4.24E+00 | 2.50E-01 | -7.25E-01 | -3.38E+01 | -1.06E+00 | -1.19E-01 | -2.91E-01 | -9.98E-03 | -1.56E-02 | -2.72E+00 | 4.32E-01 | -3.85E-01 | -1.25E+00 | 8.75E-02 | -5.52E-01 | -6.62E-01 | -1.19E-01 | -1.44E-03 |
| 1100 | 3.07E+01 | -2.10E+00 | 3.72E+00 | 6.22E+00 | -2.21E-02 | 8.60E-01 | 4.09E+01 | 2.10E+00 | -1.22E+00 | 4.85E-01 | -2.45E-02 | 6.42E-03 | 6.11E+00 | -5.72E-01 | 2.70E-01 | 2.62E+00 | -2.04E-01 | 4.07E-01 | 2.21E+00 | -8.80E-03 | 8.64E-02 |
| -300 | -3.01E+01 | -2.44E+00 | -3.61E+00 | -5.66E+00 | -1.57E-02 | -5.05E-01 | -3.90E+01 | -1.35E+00 | 4.73E-01 | -4.29E-01 | 3.50E-02 | -2.99E-03 | -4.61E+00 | -1.92E-01 | -4.96E-01 | -2.07E+00 | -3.61E-02 | -6.55E-01 | -1.61E+00 | 1.20E-01 | 7.73E-02 |

**Table S3**: IRM acquisition data table. Sample denotes sample name. **Field (mT)–** field corresponds to the intensity of the applied magnetic field. Magnetization of each sample (A/m) axis signified by Avg X, Y, and Z columns.

| Sample | | CB1703A | | | CB1711C | | | CB1734B | | | CB1821D | | | CB1824B | | | CB1827B | | | CB1869B | |
|---|---|---|---|---|---|---|---|---|---|---|---|---|---|---|---|---|---|---|---|---|---|
| **Temp (C°)** | **Avg X** | **Avg Y** | **Avg Z** | **Avg X** | **Avg Y** | **Avg Z** | **Avg X** | **Avg Y** | **Avg Z** | **Avg X** | **Avg Y** | **Avg Z** | **Avg X** | **Avg Y** | **Avg Z** | **Avg X** | **Avg Y** | **Avg Z** | **Avg X** | **Avg Y** | **Avg Z** |
| 25 | 6.12E+00 | 3.08E+00 | 2.58E+01 | 5.89E-01 | 7.41E-01 | 5.61E+00 | 6.28E-01 | 3.57E+00 | 4.22E+01 | 1.96E-02 | 1.90E-01 | 3.91E-01 | 1.51E+00 | 1.68E+00 | 4.79E+00 | 1.09E-07 | 1.43E-07 | 5.48E-07 | 2.30E-01 | 6.99E-01 | 1.78E+00 |
| 50 | 3.84E+00 | 3.41E+00 | 2.46E+01 | 3.22E-01 | 6.48E-01 | 5.24E+00 | 1.27E+00 | 6.38E+00 | 3.95E+01 | 2.88E-02 | 1.67E-01 | 3.21E-01 | 8.12E-01 | 2.01E+00 | 4.35E+00 | 8.37E-08 | 1.48E-07 | 4.69E-07 | 2.93E-01 | 8.09E-01 | 1.54E+00 |
| 86 | 6.80E+00 | 3.38E+00 | 2.21E+01 | -2.46E-01 | 9.53E-01 | 4.72E+00 | -2.24E+00 | 4.08E+00 | 3.56E+01 | -1.06E-02 | 1.43E-01 | 2.74E-01 | 1.22E+00 | 2.10E+00 | 3.76E+00 | 1.25E-07 | 1.65E-07 | 3.98E-07 | 2.13E-01 | 5.87E-01 | 1.46E+00 |
| 109 | 4.85E+00 | 4.60E+00 | 2.09E+01 | 2.36E-01 | 1.09E+00 | 4.35E+00 | -5.20E-01 | 6.78E+00 | 3.28E+01 | 1.53E-02 | 1.34E-01 | 2.42E-01 | 1.25E+00 | 1.60E+00 | 3.56E+00 | 1.05E-07 | 1.72E-07 | 3.58E-07 | 1.58E-01 | 6.59E-01 | 1.16E+00 |
| 132 | 3.09E+00 | 3.42E+00 | 1.90E+01 | -9.33E-04 | 8.36E-01 | 3.79E+00 | 1.24E+00 | 5.72E+00 | 2.87E+01 | 2.77E-03 | 1.45E-01 | 1.85E-01 | 8.34E-01 | 1.55E+00 | 3.15E+00 | 8.07E-08 | 1.32E-07 | 3.27E-07 | 2.03E-01 | 6.25E-01 | 1.00E+00 |
| 150 | 3.55E+00 | 4.16E+00 | 1.68E+01 | 5.90E-01 | 1.14E+00 | 3.32E+00 | 7.01E-01 | 3.67E+00 | 2.69E+01 | 2.20E-02 | 1.09E-01 | 1.86E-01 | 7.89E-01 | 2.36E+00 | 5.25E-01 | 6.98E-08 | 1.73E-07 | 2.84E-07 | 2.26E-01 | 5.06E-01 | 9.10E-01 |
| 209 | 4.30E+00 | 1.25E+00 | 1.22E+01 | 2.80E-01 | 8.26E-01 | 2.46E+00 | -5.37E-01 | 4.12E+00 | 1.92E+01 | 2.67E-02 | 9.95E-02 | 1.20E-01 | 1.11E+00 | 1.32E+00 | 1.89E+00 | 7.24E-08 | 1.17E-07 | 2.28E-07 | 2.30E-01 | 3.63E-01 | 7.39E-01 |
| 250 | 1.62E+00 | 1.59E+00 | 8.67E+00 | 1.13E-01 | 7.45E-01 | 1.63E+00 | 1.19E+00 | 3.47E+00 | 1.34E+01 | 2.05E-02 | 7.67E-02 | 9.14E-02 | 6.89E-01 | 1.30E+00 | 1.69E+00 | 4.14E-08 | 1.06E-07 | 2.05E-07 | 1.87E-01 | 3.50E-01 | 5.57E-01 |
| 275 | 1.21E+00 | 2.47E+00 | 6.58E+00 | 2.75E-01 | 4.90E-01 | 1.38E+00 | 9.10E-01 | 1.66E+00 | 1.10E+01 | 2.07E-02 | 5.99E-02 | 7.77E-02 | 3.91E-01 | 1.11E+00 | 1.51E+00 | 7.05E-08 | 9.78E-08 | 1.53E-07 | 1.93E-01 | 3.58E-01 | 4.01E-01 |
| 300 | 1.31E+00 | 1.22E+00 | 5.16E+00 | 2.01E-01 | 5.22E-01 | 1.10E+00 | 5.88E-01 | 2.42E+00 | 8.90E+00 | 1.73E-02 | 4.64E-02 | 7.40E-02 | 5.30E-01 | 8.09E-01 | 1.43E+00 | 8.06E-08 | 9.58E-08 | 1.32E-07 | 2.00E-01 | 3.05E-01 | 3.94E-01 |
| 325 | 1.08E+00 | 1.14E+00 | 4.06E+00 | 2.54E-01 | 3.61E-01 | 9.31E-01 | 1.14E+00 | 1.50E+00 | 7.31E+00 | 1.47E-02 | 4.20E-02 | 5.69E-02 | 6.16E-01 | 7.87E-01 | 1.11E+00 | 6.05E-08 | 6.88E-08 | 1.18E-07 | 1.96E-01 | 2.77E-01 | 3.15E-01 |
| 350 | 1.04E+00 | 1.03E+00 | 3.42E+00 | 2.00E-01 | 3.13E-01 | 8.05E-01 | 6.15E-01 | 1.19E+00 | 6.25E+00 | 4.04E-03 | 2.84E-02 | 5.36E-02 | 5.23E-01 | 7.41E-01 | 9.38E-01 | 3.83E-08 | 6.77E-08 | 1.07E-07 | 1.86E-01 | 2.40E-01 | 2.95E-01 |
| 375 | 7.20E-01 | 1.11E+00 | 3.01E+00 | 1.57E-01 | 3.27E-01 | 6.79E-01 | 6.95E-01 | 1.14E+00 | 5.41E+00 | 8.20E-03 | 2.46E-02 | 4.21E-02 | 3.90E-01 | 7.69E-01 | 6.94E-01 | -8.57E-08 | -5.27E-08 | -5.18E-08 | 1.65E-01 | 2.20E-01 | 2.61E-01 |
| 400 | 9.14E-01 | 7.82E-01 | 2.78E+00 | 8.44E-02 | 2.35E-01 | 6.76E-01 | 7.50E-01 | 1.28E+00 | 4.89E+00 | 4.10E-03 | 2.26E-02 | 3.83E-02 | 4.57E-01 | 5.63E-01 | 6.82E-01 | 3.19E-08 | 5.14E-08 | 8.45E-08 | 1.84E-01 | 1.92E-01 | 2.30E-01 |
| 500 | 6.59E-01 | 5.99E-01 | 1.83E+00 | 1.17E-01 | 2.11E-01 | 3.99E-01 | 1.49E-01 | 7.44E-01 | 3.28E+00 | 3.83E-03 | 9.05E-03 | 2.03E-02 | 2.39E-01 | 3.47E-01 | 4.69E-01 | 2.26E-08 | 2.25E-08 | 5.28E-08 | 1.11E-01 | 1.55E-01 | 1.49E-01 |
| 540 | 4.28E-01 | 5.30E-01 | 1.27E+00 | 6.55E-02 | 1.47E-01 | 2.93E-01 | 3.16E-01 | 5.48E-01 | 2.26E+00 | 3.09E-03 | 5.73E-03 | 9.48E-03 | 2.37E-01 | 2.39E-01 | 1.98E-01 | 1.49E-08 | 2.26E-08 | 3.02E-08 | 1.15E-01 | 1.22E-01 | 8.69E-02 |
| 560 | 4.53E-01 | 3.59E-01 | 9.71E-01 | 9.87E-02 | 1.18E-01 | 2.18E-01 | 4.50E-01 | 5.86E-01 | 1.64E+00 | 2.90E-03 | 3.06E-03 | 2.67E-03 | 1.92E-01 | 2.01E-01 | 8.58E-02 | 1.70E-08 | 1.51E-08 | 1.46E-08 | 1.03E-01 | 1.00E-01 | 5.72E-02 |
| 580 | 3.09E-01 | 2.97E-01 | 4.90E-01 | 4.95E-02 | 8.35E-02 | 1.27E-01 | 1.81E-01 | 4.09E-01 | 7.94E-01 | 1.18E-03 | 1.22E-03 | 2.83E-04 | 1.17E-01 | 1.31E-01 | 6.20E-02 | 9.12E-09 | 8.78E-09 | 4.60E-09 | 6.74E-02 | 6.01E-02 | 3.32E-02 |
| 600 | 2.05E-01 | 1.91E-01 | 2.07E-01 | 3.98E-02 | 6.66E-02 | 8.04E-02 | 2.35E-01 | 2.51E-01 | 4.82E-01 | 3.89E-03 | -4.04E-04 | -1.37E-03 | 8.96E-02 | 1.07E-01 | 3.85E-02 | 9.83E-09 | 6.48E-09 | 3.28E-09 | 6.05E-02 | 5.69E-02 | 2.42E-02 |
| 621 | 8.46E-02 | 7.66E-02 | 4.24E-02 | 1.82E-02 | 3.02E-02 | 1.39E-02 | 8.03E-02 | 9.74E-02 | 7.81E-02 | 2.14E-03 | 2.45E-03 | -9.55E-04 | 5.70E-02 | 6.99E-02 | 1.81E-02 | 5.85E-09 | 3.35E-09 | 1.67E-09 | 3.11E-02 | 3.35E-02 | 1.51E-02 |
| 650 | 2.45E-02 | 2.85E-02 | 1.12E-02 | 2.69E-03 | 1.07E-02 | 2.79E-03 | 2.47E-02 | 2.97E-02 | 1.18E-02 | 2.17E-03 | 2.93E-03 | 1.64E-03 | 1.08E-02 | 2.10E-02 | 6.32E-03 | 7.41E-10 | 1.74E-09 | 1.12E-09 | 1.07E-02 | 1.55E-02 | 9.01E-03 |
| 680 | 8.26E-04 | 3.54E-04 | 4.89E-04 | 8.39E-04 | -6.15E-05 | 7.09E-04 | -1.12E-05 | 1.64E-04 | 8.01E-04 | 2.97E-04 | 8.78E-04 | 5.59E-04 | 1.39E-02 | 2.44E-02 | 5.46E-03 | 2.66E-11 | 1.65E-10 | 1.71E-11 | 1.12E-04 | 1.11E-04 | 4.84E-04 |

**Table S4:** IRM demagnetization data table. Sample denotes sample name. **Temp (Cº)–** field corresponds to the temperature of the thermal demagnetization step. Magnetization of each sample (A/m) axis signified by Avg X, Y, and Z columns.

| Sample Fractions | Compositions | | | | | | | | Isotopic Ratios | | | | | | corr. coef. | Isotopic Ages (Ma) | | | | | |
|---|---|---|---|---|---|---|---|---|---|---|---|---|---|---|---|---|---|---|---|---|---|
| | Th/U | U (pg) | 206Pb* E-13 mol | 206Pb* mol% | Pb*/Pb(c) | Pb(c) (pg) | 206 Pb/204 Pb | 208 Pb/206 Pb | 207 Pb/206 Pb | err (2σ%) | 207 Pb/235 U | err (2σ%) | 206 Pb/238 U | err (2σ%) | | 207 Pb/206 Pb | err (2σ) | 207 Pb/235 U | err (2σ) | 206 Pb/238 U | err (2σ) |
| (a) | (b) | (c) | (d) | (d) | (d) | (d) | (e) | (f) | (g) | | (g) | | (g) | | | | | | | | |
| Sample KJ09-58: 38°50'57.2"N, 104°38'20.6"W | | | | | | | | | | | | | | | | | | | | | |
| **z3** | 0.12 | 831 | 0.3576 | 98.83 | 25.7 | 0.31 | 1722.8 | 0.037 | 0.04718 | (.82) | 0.06720 | (.86) | 0.010335 | (.09) | 0.412 | 57.2 | 19.7 | 66.04 | 0.55 | **66.28** | **0.06** |
| **z6** | 0.33 | 300 | 0.1291 | 96.04 | 7.7 | 0.40 | 496.0 | 0.104 | 0.04702 | (2.96) | 0.06695 | (3.04) | 0.010332 | (.22) | 0.402 | 48.9 | 70.6 | 65.80 | 1.94 | **66.27** | **0.15** |
| **z2** | 0.14 | 751 | 0.3232 | 98.73 | 23.8 | 0.30 | 1589.8 | 0.044 | 0.04736 | (.91) | 0.06744 | (.95) | 0.010332 | (.09) | 0.425 | 66.5 | 21.8 | 66.27 | 0.61 | **66.26** | **0.06** |
| **z5** | 0.23 | 526 | 0.2261 | 97.80 | 13.8 | 0.38 | 906.2 | 0.072 | 0.04708 | (1.58) | 0.06702 | (1.63) | 0.010329 | (.13) | 0.381 | 52.3 | 37.7 | 65.87 | 1.04 | **66.25** | **0.09** |
| **z1** | 0.38 | 232 | 0.0999 | 96.01 | 7.8 | 0.31 | 501.3 | 0.121 | 0.04737 | (2.72) | 0.06743 | (2.80) | 0.010329 | (.23) | 0.372 | 66.7 | 64.8 | 66.26 | 1.79 | **66.24** | **0.15** |
| **z7** | 1.25 | 249 | 0.1071 | 95.78 | 9.1 | 0.35 | 470.8 | 0.398 | 0.04715 | (3.01) | 0.06712 | (3.11) | 0.010328 | (.25) | 0.425 | 56.0 | 71.8 | 65.96 | 1.98 | **66.24** | **0.16** |
| **z4** | 0.26 | 845 | 0.3634 | 98.48 | 19.9 | 0.42 | 1286.2 | 0.082 | 0.04722 | (1.09) | 0.06719 | (1.11) | 0.010325 | (.09) | 0.296 | 59.2 | 25.9 | 66.03 | 0.71 | **66.22** | **0.06** |

**Table S5: A.** U-Pb isotopic data for the analyzed zircons of the Lower Feral Llama Gulley tuff sample (KJ0958), Leaf 6 section, Denver Basin, Colorado. **(a)** Thermally annealed and pre-treated single zircon. Data used in age calculation are in bold. **(b)** Model Th/U ratio calculated from radiogenic 208Pb/206Pb ratio and the sample age. **(c)** Total sample U. **(d)** Pb* and Pbc represent radiogenic and common Pb, respectively; mol % 206Pb* with respect to total radiogenic and laboratory blank Pb. **(e)** Measured ratio corrected for spike and fractionation only. **(f)** Radiogenic Pb ratio. **(g)** Corrected for fractionation, spike and blank. Also corrected for initial Th/U disequilibrium using radiogenic $^{208}$Pb and Th/U[magma] = 2.8. Mass fractionation correction of 0.25%/amu ± 0.04%/amu (atomic mass unit) was applied to single-collector Daly analyses. All common Pb assumed to be laboratory blank. Total procedural blank less than 0.1 pg for U. Blank isotopic composition: $^{206}$Pb/$^{204}$Pb = 18.42 ± 0.35, $^{207}$Pb/$^{204}$Pb =15.36 ± 0.23, $^{208}$Pb/$^{204}$Pb = 37.46 ± 0.74. Corr. coef. = correlation coefficient. Ages calculated using the decay constants $\lambda_{238}$ = 1.55125E-10 and $\lambda_{235}$ = 9.8485E-10 (Jaffey et al. 1971).

| Analysis | 207Pb*/235U* | ±2σ (%) | 206Pb*/238U | ±2σ (%) | error corr. | 238U/206Pb* | ±2σ (%) | 207Pb*/206Pb* | ±2σ (%) | error corr. | CA-IDTIMS z# | ±2σ (Ma) | 207Pb*/206Pb* | ±2σ (Ma) | 207Pb*/235U | ±2σ (Ma) | 206Pb*/238U* | ±2σ (Ma) |
|---|---|---|---|---|---|---|---|---|---|---|---|---|---|---|---|---|---|---|
| KJ-1702_M_69 | 0.0851 | 17.59 | 0.0094 | 4.95 | 0.28 | 105.829 | 4.95 | 0.0653 | 16.88 | 0 | **z4** | 14.41 | 783 | 355 | 83 | 14 | 61 | 3 |
| KJ-1702_L_36 | 0.0533 | 20.45 | 0.0095 | 5.63 | 0.27 | 105.140 | 5.63 | 0.0406 | 19.66 | 4.915E-16 | **z1** | 7 | -312 | 504 | 53 | 10 | 61 | 3 |
| KJ-1702_M_43 | 0.0656 | 5.92 | 0.0096 | 4.25 | 0.71 | 104.511 | 4.25 | 0.0497 | 4.12 | 0 | **z5** | 3.08 | 182 | 96 | 65 | 4 | 61 | 3 |
| KJ-1702_L_37 | 0.0715 | 19.03 | 0.0098 | 6.74 | 0.35 | 101.756 | 6.74 | 0.0528 | 17.80 | 0 | **z3** | 35.55 | 319 | 404 | 70 | 13 | 63 | 4 |
| KJ-1702_M_46 | 0.0719 | 17.59 | 0.0099 | 7.20 | 0.41 | 101.437 | 7.20 | 0.0529 | 16.05 | 0 | **z8** | 19.13 | 324 | 364 | 70 | 12 | 63 | 5 |
| KJ-1702_M_78 | 0.0782 | 14.04 | 0.0099 | 2.75 | 0.19 | 100.594 | 2.75 | 0.0570 | 13.77 | -3.5947E-16 | **z6** | 3.28 | 492 | 304 | 76 | 10 | 64 | 2 |
| KJ-1702_L_40 | 0.0773 | 12.73 | 0.0100 | 4.16 | 0.32 | 100.160 | 4.16 | 0.0562 | 12.03 | 0 | **z2** | 49.79 | 460 | 267 | 76 | 9 | 64 | 3 |
| KJ-1702_M_50 | 0.0537 | 15.83 | 0.0101 | 5.54 | 0.35 | 99.096 | 5.54 | 0.0386 | 14.83 | 0 | **z7** | 9.31 | -442 | 390 | 53 | 8 | 65 | 4 |
| KJ-1702_L_34 | 0.0714 | 16.76 | 0.0101 | 6.05 | 0.36 | 98.564 | 6.05 | 0.0510 | 15.63 | 0 | | 18 | 241 | 360 | 70 | 11 | 65 | 4 |
| KJ-1702_M_58 | 0.0662 | 11.04 | 0.0103 | 4.05 | 0.36 | 97.472 | 4.05 | 0.0468 | 10.27 | 0 | **z10** | 29.63 | 40 | 246 | 65 | 7 | 66 | 3 |
| KJ-1702_L_25 | 0.0857 | 21.17 | 0.0103 | 6.64 | 0.31 | 97.268 | 6.64 | 0.0605 | 20.10 | 2.032E-16 | **z9** | 35 | 621 | 434 | 84 | 17 | 66 | 4 |
| KJ-1702_L_30 | 0.0746 | 7.90 | 0.0103 | 5.91 | 0.74 | 97.096 | 5.91 | 0.0526 | 5.24 | 0 | | 14 | 310 | 119 | 73 | 6 | 66 | 4 |
| KJ-1702_M_75 | 0.0920 | 12.61 | 0.0104 | 7.30 | 0.58 | 96.448 | 7.30 | 0.0644 | 10.27 | 0 | **z11** | 15.20 | 753 | 217 | 89 | 11 | 66 | 5 |
| KJ-1702_L_14 | 0.0711 | 8.44 | 0.0104 | 4.53 | 0.53 | 96.351 | 4.53 | 0.0497 | 7.12 | 0 | | 16 | 179 | 166 | 70 | 6 | 67 | 3 |
| KJ-1702_L_1 | 0.0651 | 9.70 | 0.0104 | 4.57 | 0.47 | 96.293 | 4.57 | 0.0454 | 8.55 | -1.75E-16 | | 8 | -32 | 207 | 64 | 6 | 67 | 3 |
| KJ-1702_L_32 | 0.0935 | 13.81 | 0.0104 | 6.20 | 0.45 | 96.248 | 6.20 | 0.0653 | 12.34 | 0 | | 33 | 784 | 259 | 91 | 12 | 67 | 4 |
| KJ-1702_M_60 | 0.0802 | 18.76 | 0.0104 | 5.60 | 0.30 | 96.090 | 5.60 | 0.0559 | 17.91 | 0 | **z12** | 33.27 | 448 | 398 | 78 | 14 | 67 | 4 |
| KJ-1702_M_71 | 0.0703 | 12.85 | 0.0104 | 6.56 | 0.51 | 96.083 | 6.56 | 0.0490 | 11.05 | -3.749E-16 | **z13** | 10.09 | 146 | 259 | 69 | 9 | 67 | 4 |
| KJ-1702_M_53 | 0.0739 | 13.27 | 0.0104 | 5.25 | 0.39 | 95.976 | 5.25 | 0.0514 | 12.18 | 2.12432E-16 | | 11.56 | 261 | 280 | 72 | 9 | 67 | 3 |
| KJ-1702_L_24 | 0.0731 | 9.96 | 0.0104 | 6.56 | 0.65 | 95.898 | 6.56 | 0.0508 | 7.50 | -1.39E-16 | | 7 | 233 | 173 | 72 | 7 | 67 | 4 |
| KJ-1702_L_28 | 0.0705 | 20.91 | 0.0105 | 7.91 | 0.38 | 95.152 | 7.91 | 0.0487 | 19.35 | -1.771E-16 | | 31.08 | 131 | 455 | 69 | 14 | 67 | 5 |
| KJ-1702_M_67 | 0.0689 | 13.83 | 0.0105 | 3.87 | 0.27 | 95.150 | 3.87 | 0.0475 | 13.28 | 5.30601E-16 | | 9.56 | 77 | 315 | 68 | 9 | 67 | 3 |
| KJ-1702_L_5 | 0.1132 | 27.71 | 0.0105 | 8.92 | 0.32 | 94.938 | 8.92 | 0.0779 | 26.24 | -4.6353E-16 | | 52.96 | 1145 | 521 | 109 | 29 | 68 | 6 |
| KJ-1702_M_64 | 0.0947 | 18.84 | 0.0107 | 6.24 | 0.33 | 93.089 | 6.24 | 0.0639 | 17.77 | 7.34203E-16 | | 19.67 | 738 | 376 | 92 | 17 | 69 | 4 |
| KJ-1702_M_72 | 0.0957 | 19.38 | 0.0108 | 6.79 | 0.35 | 92.492 | 6.79 | 0.0642 | 18.15 | 4.40441E-16 | | 19.33 | 748 | 383 | 93 | 17 | 69 | 5 |
| KJ-1702_L_21 | 0.0814 | 15.42 | 0.0110 | 4.99 | 0.32 | 90.836 | 4.99 | 0.0536 | 14.59 | 1.86403E-16 | | 24.47 | 356 | 329 | 79 | 12 | 71 | 4 |
| KJ-1702_M_52 | 0.0848 | 8.13 | 0.0110 | 4.08 | 0.49 | 90.760 | 4.08 | 0.0559 | 7.03 | -2.372E-16 | | 4.72 | 446 | 156 | 83 | 6 | 71 | 3 |
| KJ-1702_M_47 | 0.1340 | 22.34 | 0.0110 | 10.34 | 0.46 | 90.749 | 10.34 | 0.0882 | 19.80 | 1.32416E-16 | | 41.61 | 1387 | 380 | 128 | 27 | 71 | 7 |
| KJ-1702_M_57 | 0.0709 | 18.67 | 0.0111 | 6.04 | 0.32 | 89.787 | 6.04 | 0.0462 | 17.67 | 2.54597E-16 | | 16.27 | 6 | 425 | 70 | 13 | 71 | 4 |
| KJ-1702_M_63 | 0.0869 | 19.22 | 0.0112 | 8.11 | 0.42 | 89.633 | 8.11 | 0.0565 | 17.42 | 0 | | 9.95 | 472 | 386 | 85 | 16 | 72 | 6 |
| KJ-1702_L_39 | 0.0863 | 14.16 | 0.0112 | 6.30 | 0.44 | 89.010 | 6.30 | 0.0557 | 12.68 | 1.7015E-16 | | 20.56 | 442 | 282 | 84 | 11 | 72 | 5 |
| KJ-1702_M_79 | 0.0717 | 17.18 | 0.0116 | 6.31 | 0.36 | 86.321 | 6.31 | 0.0449 | 15.98 | 2.69748E-16 | | 7.10 | -61 | 390 | 70 | 12 | 74 | 5 |
| KJ-1702_L_6 | 0.0570 | 26.92 | 0.0117 | 6.01 | 0.22 | 85.232 | 6.01 | 0.0352 | 26.24 | -3.44E-16 | | 9 | -692 | 727 | 56 | 15 | 75 | 4 |
| KJ-1702_L_18 | 0.0712 | 22.89 | 0.0118 | 7.30 | 0.32 | 84.734 | 7.30 | 0.0438 | 21.69 | 0 | | 25.22 | -123 | 535 | 70 | 15 | 76 | 5 |
| KJ-1702_M_59 | 0.0996 | 18.15 | 0.0121 | 9.05 | 0.50 | 82.813 | 9.05 | 0.0598 | 15.74 | 1.90695E-16 | | 14.19 | 596 | 341 | 96 | 17 | 77 | 7 |
| KJ-1702_M_44 | 0.9531 | 25.65 | 0.0692 | 25.63 | 1.00 | 14.449 | 25.63 | 0.0999 | 1.13 | -8.3925E-16 | | 145.24 | 1622 | 21 | 680 | 127 | 431 | 107 |
| KJ-1702_M_62 | 1.2239 | 22.28 | 0.1000 | 21.74 | 0.98 | 10.004 | 21.74 | 0.0888 | 4.87 | 0 | | 112.99 | 1400 | 93 | 812 | 124 | 614 | 127 |
| KJ-1702_L_22 | 2.0053 | 5.51 | 0.1435 | 5.39 | 0.97 | 6.970 | 5.39 | 0.1014 | 1.10 | -2.63E-16 | | 142 | 1649 | 20 | 1117 | 37 | 864 | 44 |
| KJ-1702_L_11 | 1.8249 | 7.78 | 0.1477 | 7.59 | 0.97 | 6.770 | 7.59 | 0.0896 | 1.71 | 6.498E-16 | | 83 | 1417 | 33 | 1054 | 51 | 888 | 63 |
| KJ-1702_M_41 | 2.6345 | 9.19 | 0.1882 | 8.98 | 0.97 | 5.314 | 8.98 | 0.1015 | 1.97 | -5.0712E-16 | | 132.67 | 1652 | 36 | 1310 | 68 | 1112 | 92 |
| KJ-1702_L_12 | 2.4822 | 2.12 | 0.1903 | 1.19 | 0.44 | 5.254 | 1.19 | 0.0946 | 1.76 | -2.27E-16 | | 63 | 1520 | 33 | 1267 | 15 | 1123 | 12 |
| KJ-1702_M_61 | 2.5069 | 12.40 | 0.1913 | 12.25 | 0.99 | 5.228 | 12.25 | 0.0951 | 1.92 | 0 | | 186.31 | 1529 | 36 | 1274 | 90 | 1128 | 127 |
| KJ-1702_M_74 | 2.5732 | 5.28 | 0.1945 | 5.13 | 0.96 | 5.140 | 5.13 | 0.0959 | 1.23 | 0 | | 66.70 | 1546 | 23 | 1293 | 39 | 1146 | 54 |
| KJ-1702_M_80 | 2.8045 | 13.42 | 0.1980 | 13.34 | 0.99 | 5.049 | 13.34 | 0.1027 | 1.46 | -5.0776E-16 | | 52.89 | 1674 | 27 | 1357 | 100 | 1165 | 142 |
| KJ-1702_L_4 | 2.7170 | 6.02 | 0.2105 | 5.23 | 0.86 | 4.751 | 5.23 | 0.0936 | 2.97 | -1.25E-16 | | 101 | 1500 | 56 | 1333 | 45 | 1231 | 59 |
| KJ-1702_L_31 | 2.8284 | 5.18 | 0.2144 | 4.97 | 0.95 | 4.664 | 4.97 | 0.0957 | 1.49 | -2.5494E-16 | | 103.11 | 1541 | 28 | 1363 | 39 | 1252 | 57 |
| KJ-1702_L_35 | 2.9308 | 7.19 | 0.2159 | 7.04 | 0.97 | 4.631 | 7.04 | 0.0984 | 1.44 | 0 | | 111 | 1595 | 27 | 1390 | 54 | 1260 | 81 |
| KJ-1702_L_13 | 2.8241 | 3.99 | 0.2188 | 2.81 | 0.68 | 4.570 | 2.81 | 0.0936 | 2.82 | -1.11E-16 | | 107 | 1500 | 53 | 1362 | 30 | 1276 | 33 |
| KJ-1702_L_9 | 2.7969 | 4.25 | 0.2268 | 2.85 | 0.65 | 4.410 | 2.85 | 0.0894 | 3.15 | 0 | | 84.17 | 1414 | 60 | 1355 | 32 | 1318 | 34 |
| KJ-1702_L_29 | 2.9998 | 5.75 | 0.2364 | 5.16 | 0.89 | 4.230 | 5.16 | 0.0920 | 2.52 | 1.399E-16 | | 45 | 1468 | 48 | 1408 | 44 | 1368 | 64 |
| KJ-1702_L_33 | 3.3817 | 6.53 | 0.2367 | 6.28 | 0.95 | 4.225 | 6.28 | 0.1036 | 1.80 | -1.91E-16 | | 90 | 1690 | 33 | 1500 | 51 | 1369 | 77 |
| KJ-1702_L_26 | 2.8976 | 5.03 | 0.2422 | 3.60 | 0.70 | 4.128 | 3.60 | 0.0868 | 3.52 | 0 | | 60 | 1355 | 68 | 1381 | 38 | 1398 | 45 |
| KJ-1702_L_17 | 3.3037 | 4.47 | 0.2453 | 3.96 | 0.87 | 4.077 | 3.96 | 0.0977 | 2.08 | 0 | | 101 | 1580 | 39 | 1482 | 35 | 1414 | 50 |
| KJ-1702_L_2 | 3.2446 | 7.44 | 0.2460 | 7.21 | 0.96 | 4.065 | 7.21 | 0.0957 | 1.84 | 0 | | 106 | 1541 | 35 | 1468 | 58 | 1418 | 92 |
| KJ-1702_L_19 | 3.6821 | 4.60 | 0.2535 | 3.93 | 0.84 | 3.945 | 3.93 | 0.1054 | 2.39 | 2.062E-16 | | 88 | 1721 | 44 | 1567 | 37 | 1456 | 51 |
| KJ-1702_M_56 | 3.6816 | 2.97 | 0.2536 | 1.96 | 0.61 | 3.943 | 1.96 | 0.1053 | 2.23 | -2.27E-16 | | 52.93 | 1720 | 41 | 1567 | 24 | 1457 | 26 |
| KJ-1702_M_55 | 3.8605 | 4.54 | 0.2610 | 4.13 | 0.89 | 3.831 | 4.13 | 0.1073 | 1.88 | 0 | | 56.39 | 1753 | 34 | 1605 | 37 | 1495 | 55 |
| KJ-1702_M_70 | 4.1923 | 4.00 | 0.2802 | 3.65 | 0.89 | 3.569 | 3.65 | 0.1085 | 1.64 | -1.6512E-16 | | 84.37 | 1775 | 30 | 1673 | 33 | 1592 | 52 |
| KJ-1702_L_38 | 4.9463 | 6.41 | 0.2917 | 3.46 | 0.53 | 3.428 | 3.46 | 0.1230 | 5.40 | 1.86998E-16 | | 182.34 | 1999.82 | 95.86 | 1810 | 54 | 1650 | 50 |
| KJ-1702_L_15 | 4.3063 | 2.80 | 0.2957 | 1.85 | 0.60 | 3.382 | 1.85 | 0.1056 | 2.10 | 0 | | 71 | 1725 | 39 | 1695 | 23 | 1670 | 27 |

**Table S6:** LA_ICPMS U-Pb geochronologic analyses for Sample KJ1702**.** Isotope ratios and ages are not corrected for initial common Pb. Isotope ratio and apparent age errors do not include systematic calibration errors of 0.77% (207Pb/206Pb), 1.04% (206Pb/238U) (all 2-sigma). Isotope ratios and ages corrected using a measured linear secondary standard age bias - 206Pb count rate relationship. Sweep-by-sweep downhole fractionation of U/Pb ratios not corrected via Si/Zr fractionation factor.

| | Compositional Parameters | | | | | | Radiogenic Isotope Ratios | | | | | | | | Isotopic Ages | | | | | |
|---|---|---|---|---|---|---|---|---|---|---|---|---|---|---|---|---|---|---|---|---|
| Sample | $\frac{Th}{U}$ | $^{206}Pb^*$ x$10^{-13}$ mol | mol % $^{206}Pb^*$ | $\frac{Pb^*}{Pb_c}$ | $Pb_c$ (pg) | $\frac{^{206}Pb}{^{204}Pb}$ | $\frac{^{208}Pb}{^{206}Pb}$ | $\frac{^{207}Pb}{^{206}Pb}$ | % err | $\frac{^{207}Pb}{^{235}U}$ | % err | $\frac{^{206}Pb}{^{238}U}$ | % err | corr. coef. | $\frac{^{207}Pb}{^{206}Pb}$ | ± | $\frac{^{207}Pb}{^{235}U}$ | ± | $\frac{^{206}Pb}{^{238}U}$ | ± |
| (a) | (b) | (c) | (c) | (c) | (c) | (d) | (e) | (e) | (f) | (e) | (f) | (e) | (f) | | (g) | (f) | (g) | (f) | (g) | (f) |
| **KJ-1702** | | | | | | | | | | | | | | | | | | | | |
| z1 | 0.467 | 0.1775 | 98.58% | 21 | 0.21 | 1273 | 0.150 | 0.047337 | 0.398 | 0.067414 | 0.445 | 0.010329 | 0.076 | 0.678 | 66.27 | 9.48 | 66.24 | 0.29 | 66.242 | 0.050 |
| z2 | 0.056 | 0.4122 | 97.73% | 11 | 0.79 | 795 | 0.018 | 0.047180 | 0.763 | 0.067177 | 0.820 | 0.010327 | 0.108 | 0.579 | 58.35 | 18.18 | 66.02 | 0.52 | 66.229 | 0.071 |
| z3 | 0.063 | 0.6933 | 99.60% | 66 | 0.23 | 4507 | 0.020 | 0.047330 | 0.137 | 0.067422 | 0.184 | 0.010332 | 0.066 | 0.805 | 65.89 | 3.26 | 66.25 | 0.12 | 66.261 | 0.043 |
| z5 | 0.683 | 0.1151 | 97.03% | 10 | 0.29 | 607 | 0.219 | 0.047208 | 0.944 | 0.067518 | 1.017 | 0.010373 | 0.110 | 0.697 | 59.75 | 22.49 | 66.34 | 0.65 | 66.524 | 0.073 |
| z6 | 0.218 | 0.1817 | 98.32% | 16 | 0.26 | 1077 | 0.070 | 0.047164 | 0.535 | 0.067169 | 0.589 | 0.010329 | 0.076 | 0.744 | 57.52 | 12.75 | 66.01 | 0.38 | 66.244 | 0.050 |
| z7 | 0.122 | 0.1189 | 95.55% | 6 | 0.46 | 405 | 0.039 | 0.046408 | 1.252 | 0.066050 | 1.345 | 0.010322 | 0.126 | 0.762 | 18.88 | 30.04 | 64.94 | 0.85 | 66.201 | 0.083 |
| z8 | 0.117 | 0.1356 | 95.41% | 6 | 0.54 | 393 | 0.037 | 0.046238 | 1.248 | 0.065758 | 1.340 | 0.010315 | 0.136 | 0.709 | 10.02 | 30.00 | 64.67 | 0.84 | 66.151 | 0.089 |
| z9 | 0.100 | 0.3829 | 99.35% | 41 | 0.21 | 2760 | 0.032 | 0.047340 | 0.185 | 0.067423 | 0.231 | 0.010329 | 0.067 | 0.763 | 66.40 | 4.40 | 66.25 | 0.15 | 66.247 | 0.044 |
| z10 | 0.101 | 0.1596 | 97.01% | 9 | 0.41 | 604 | 0.032 | 0.047285 | 0.731 | 0.067384 | 0.797 | 0.010336 | 0.111 | 0.640 | 63.66 | 17.39 | 66.21 | 0.51 | 66.28 | 0.07 |
| z11 | 0.764 | 0.0766 | 97.01% | 10 | 0.20 | 604 | 0.245 | 0.047348 | 0.776 | 0.067701 | 0.843 | 0.010370 | 0.101 | 0.700 | 66.79 | 18.46 | 66.52 | 0.54 | 66.51 | 0.07 |
| z12 | 0.070 | 0.1810 | 98.66% | 20 | 0.20 | 1343 | 0.022 | 0.047322 | 0.381 | 0.067428 | 0.428 | 0.010334 | 0.074 | 0.691 | 65.50 | 9.06 | 66.26 | 0.27 | 66.28 | 0.05 |
| z13 | 0.161 | 0.1433 | 98.17% | 15 | 0.22 | 988 | 0.052 | 0.047345 | 0.479 | 0.067411 | 0.531 | 0.010327 | 0.087 | 0.653 | 66.64 | 11.38 | 66.24 | 0.34 | 66.23 | 0.06 |

**Table S7** U-Th-Pb isotopic data for analyzed zircons from sample KJ1702. **(a)** z1, z2 etc. are labels for single zircon grains or fragments annealed and chemically abraded after Mattinson (2005). **(b)** Model Th/U ratio iteratively calculated from the radiogenic 208Pb/206Pb ratio and 206Pb/238U age. **(c)** Pb* and Pbc represent radiogenic and common Pb, respectively; mol % $^{206}Pb^*$ with respect to radiogenic, blank and initial common Pb. **(d)** Measured ratio corrected for spike and fractionation only. Fractionation estimated at 0.18 +/- 0.03 %/a.m.u. for Daly analyses, based on analysis of NBS-981 and NBS-982. **(e)** Corrected for fractionation, spike, and common Pb; up to 1 pg of common Pb was assumed to be procedural blank: 206Pb/204Pb = 18.042 ± 0.61%; 207Pb/204Pb = 15.537 ± 0.52%; 208Pb/204Pb = 37.686 ± 0.63% (all uncertainties 1-sigma). Excess over blank was assigned to initial common Pb, using the Stacey and Kramers (1975) two-stage Pb isotope evolution model at the nominal sample age. **(f)** Errors are 2-sigma, propagated using the algorithms of Schmitz and Schoene (2007). **(g)** Calculations are based on the decay constants of Jaffey et al. (1971). 206Pb/238U and 207Pb/206Pb ages corrected for initial disequilibrium in 230Th/238U using Th/U [magma] = 3.

| Sample | Section | K-Taxa | Total | % | Elev (m) |
|---|---|---|---|---|---|
| ADB1703-01 | Leaf 6 | 38 | 685 | 5.5 | 1883.7 |
| ADB1703-02 | Leaf 6 | 66 | 938 | 7.0 | 1884.1 |
| ADB1703-03 | Leaf 6 | 10 | 631 | 1.6 | 1900.4 |
| ADB1703-04 | Leaf 6 | 62 | 719 | 8.6 | 1915.3 |
| ADB1703-04c | Leaf 6 | 133 | 505 | 26.3 | 1919.2 |
| ADB1703-04b | Leaf 6 | 5 | 562 | 0.9 | 1920.0 |
| ADB1703-04d | Leaf 6 | 65 | 790 | 8.2 | 1924.5 |
| ADB1703-04e | Leaf 6 | 5 | 667 | 0.7 | 1926.7 |
| ADB1703-05 | Leaf 6 | 0 | 745 | 0.0 | 1927.2 |
| ADB1703-06 | Leaf 6 | 0 | 1173 | 0.0 | 1956.5 |
| ADB1802-02 | 9789 | 62 | 635 | 9.8 | 1899.3 |
| ADB1802-03 | 9789 | 13 | 405 | 3.2 | 1905.1 |
| ADB1802-04 | 9789 | 76 | 122 | 62.3 | *1905.2* |
| ADB1802-05 | 9789 | 13 | 593 | 2.2 | 1905.4 |
| ADB1802-06 | 9789 | 9 | 417 | 2.2 | 1905.4 |
| ADB1802-07 | 9789 | 3 | 664 | 0.5 | 1915.1 |
| ADB1802-09 | 9789 | 2 | 760 | 0.3 | 1918.7 |
| ADB1802-08 | 9789 | 0 | 560 | 0.0 | 1919.4 |
| ADB1802-10 | 9789 | 5 | 638 | 0.8 | 1925.6 |
| ADB1802-11 | 9789 | 0 | 573 | 0.0 | 1939.3 |
| ADB1802-12 | 9789 | 0 | 777 | 0.0 | 1950.4 |
| ADB1802-13 | 9789 | 7 | 718 | 1.0 | 1955.3 |
| ADB1701-0 | Bishop Wash | 128 | 670 | 19.1 | 1900.1 |
| ADB1701-0a | Bishop Wash | 147 | 640 | 23.0 | 1903.0 |
| ADB1701-01 | Bishop Wash | 0 | 532 | 0.0 | 1904.4 |
| ADB1701-02 | Bishop Wash | 6 | 635 | 0.9 | 1908.2 |
| ADB1701-03 | Bishop Wash | 22 | 577 | 3.8 | 1908.6 |
| ADB1701-05b | Bishop Wash | 6 | 856 | 0.7 | 1920.8 |
| ADB1701-05a | Bishop Wash | 4 | 857 | 0.5 | 1919.6 |
| ADB1701-06 | Bishop Wash | 2 | 708 | 0.3 | 1923.3 |
| ADB1701-07 | Bishop Wash | 12 | 704 | 1.7 | 1924.3 |
| ADB1701-08 | Bishop Wash | 2 | 595 | 0.3 | 1933.4 |
| ADB1701-09 | Bishop Wash | 7 | 842 | 0.8 | 1935.0 |
| ADB1701-10 | Bishop Wash | 6 | 659 | 0.9 | 1936.7 |
| ADB1701-13 | Bishop Wash | 8 | 785 | 1.0 | 1944.8 |
| ADB1701-15 | Bishop Wash | 2 | 881 | 0.2 | 1947.9 |
| ADB1701-16 | Bishop Wash | 4 | 621 | 0.6 | 1948.3 |
| ADB1701-18 | Bishop Wash | 8 | 574 | 1.4 | 1955.8 |
| ADB1701-19 | Bishop Wash | 0 | 654 | 0.0 | 1962.0 |
| ADB1701-20 | Bishop Wash | 1 | 580 | 0.2 | 1966.6 |
| ADB1701-21 | Bishop Wash | 0 | 592 | 0.0 | 1969.2 |
| ADB1701-23 | Bishop Wash | 2 | 586 | 0.3 | 1976.1 |
| ADB1701-28 | Bishop Wash | 0 | 714 | 0.0 | 1992.7 |
| ADB1701-29 | Bishop Wash | 3 | 720 | 0.4 | 1997.3 |
| ADB1701-30 | Bishop Wash | 0 | 734 | 0.0 | 2001.7 |
| ADB1701-31 | Bishop Wash | 3 | 735 | 0.4 | 2005.7 |

**Table S8:** Tabulated pollen counts for the Leaf 6, 9789, and Bishop Wash sections. **Sample-** pollen sample number, **Section-** corresponding lithostratigraphic section, **K-Taxa-** individual Cretaceous palynomorph counts in sample, **Total-** total palynomorph counts in sample, **% -** Percent of K taxa, **Elev (m)-** DGPS elevation in meters of sample, italicized value represents synthetic elevation (see methods).
